# Huntingtin is a cell-autonomous regulator of neuropeptide trafficking and clock output

**DOI:** 10.64898/2026.07.22.740122

**Authors:** Agustina Bruno-Vignolo, Cayetana Arnaiz, Florencia Fernandez-Chiappe, Ana Ricciuti, Lautaro Alejandro Duarte, Ivana Ducrey, Ivana Marcela Linenberg, Cora Noemi Pollak, Pedro Lorenzo Ballestero, Marina Propato-Lots, Camila Sofía Falzone, Esteban Javier Beckwith, Tomás Luis Falzone, Nara Ines Muraro

## Abstract

Huntington’s disease is a severe neurodegenerative condition arising from an abnormal CAG repeat expansion in the *HTT* gene, which leads to the production of a mutant Huntingtin protein carrying an extended polyglutamine stretch. Although the field has largely centred on the toxic effects gained by this mutant protein, growing evidence points to the loss of normal wild-type Huntingtin function as an additional contributor to disease progression. Despite this, the cell-intrinsic roles of wild-type Huntingtin in neuronal biology remain poorly defined, in part because disentangling its specific contributions from broader network-level effects has proven technically challenging. To address this knowledge gap, we took advantage of the *Drosophila* huntingtin homolog (*htt*) and selectively manipulated its expression in the small lateral ventral neurons (sLNvs), a discrete cluster of just eight circadian pacemaker neurons that govern behavioral rhythmicity and sleep. Through targeted genetic knock-down, we show that reducing *htt* levels in sLNvs weakens the robustness of free-running circadian rhythms and substantially increases sleep in female flies. These behavioral changes are not rooted in developmental abnormalities, as restricting *htt* knock-down to adult flies reproduces the sleep phenotype across both beam-crossing and video-based locomotion assays. At the cellular level, *htt* loss disrupts dense core vesicle trafficking along sLNv axons, altering the fraction of motile vesicles and their velocity, and abolishing the time-of-day-dependent fluctuations in vesicle dynamics observed in these neurons. Complementary electrophysiological recordings using whole-cell patch-clamp further reveal that *htt* knock-down lowers action potential firing rates without perturbing resting membrane potential. Together, these results identify huntingtin as a cell-autonomous regulator of neuropeptide trafficking, neuronal excitability and circadian output. Beyond advancing our understanding of wild-type huntingtin physiology, this work carries direct relevance for HD therapeutic strategies, particularly those involving huntingtin-lowering approaches, by highlighting functions that may be unintentionally compromised.

## INTRODUCTION

Huntington’s disease (HD) is a neurodegenerative disorder characterized by motor deficits and non-motor symptoms (1), including disrupted circadian rhythms and sleep (2). HD results from an autosomal dominant CAG repeat expansion in exon 1 of the *Huntingtin* (*HTT*) gene, which leads to an elongated polyglutamine (polyQ) tract in the amino-terminal domain (3). The length of this polyQ expansion correlates with disease severity and age at onset (4), and is thought to drive HTT aggregation, ultimately disrupting cellular homeostasis. Given the dominant inheritance pattern of HD, research efforts have predominantly focused on elucidating the gain-of-function (GOF) properties of the mutant protein. The GOF hypothesis proposes that HD pathology is mainly the result of toxicity caused by the polyQ elongated HTT (HTT^polyQ^). This hypothesis is reinforced by the evidence that polyQ expansions in functionally unrelated proteins can independently give rise to neurodegenerative disorders, including various forms of spinocerebral ataxias (5). However, studies also suggest that a loss-of-function (LOF) of wild-type HTT (HTT^WT^) may also be involved in the etiology of HD (6). Consequently, HD may result from a combination of mutant HTT^polyQ^ toxicity with a partial loss of normal HTT^WT^ function, supporting a combined gain-of-function and loss-of-function (GOF+LOF) model. The detrimental consequences of HTT^WT^ LOF are not unexpected, considering its diverse and fundamental functions which include intracellular trafficking, autophagy regulation and transcriptional control (7). Moreover, HTT^WT^ has been reported to exert a neuroprotective function (6). There is currently no cure for HD, however, treatments aiming at downregulating the levels of HTT^polyQ^ are being tested. Critically, it has been shown that these types of therapies may also decrease the levels of HTT^WT^, with potentially deleterious effects due to LOF of its physiological role (8). Therefore, dissecting the role of HTT^WT^ in neuronal physiology is key to separating its LOF consequences from the pathogenic GOF effects.

Due to the numerous experimental advantages it provides, the fruit fly *Drosophila melanogaster* has been exploited as a powerful model system to investigate the mechanisms underlying human neurodegenerative diseases (9). In the case of HD, the over-expression of human HTT^polyQ^ has been used to unravel molecular mechanisms behind its aggregation and toxicity (10). The *Drosophila* huntingtin (*htt*, CG9995, FBgn0027655), which encodes the ortholog of the human HTT protein (11), has been considerably less studied. *Drosophila htt* null mutants are viable but present a shorter life-span and reduced locomotion in aged flies (12). Of note, *Drosophila htt* lacks the polyQ tract that is elongated in humans suffering from HD (11), a molecular feature absent until the evolution of vertebrate HTT (6). Despite this, several examples demonstrate conserved functions between mammalian *HTT* and the *Drosophila* ortholog via molecular replacement strategies (13,14). Evidence from *Drosophila* suggests that fly htt fulfills functions analogous to vertebrate HTT, including the regulation of development (15), autophagy (13,16), and axonal transport of different cargo (14,17,18).

The aim of this study is to elucidate the cell-autonomous contribution of *htt* to neuronal physiology and function. To this end, we used the genetically tractable fruit fly and selectively manipulated the small lateral ventral neurons (sLNvs), a restricted neuronal subset key to circadian control of behaviour and sleep (19–21). This strategy isolates cell-autonomous effects of *htt* from broader network-driven consequences since the sLNvs comprise only 8 neurons in the adult *Drosophila* brain (four per hemibrain). Moreover, this restricted neuronal group allows the assessment of key cellular, electrophysiological and behavioural outputs. These neurons are key for behavioural rhythms, communicating with other clock and non-clock neurons through the release of neuropeptides such as Pigment Dispersing Factor (PDF) (19,22–24), which controls both the timing and architecture of sleep (20,21). Because huntingtin plays a role in the axonal transport of neuropeptide-containing dense core vesicles (DCV), sLNvs represent an attractive neuronal population in which to investigate its relation to neuropeptide trafficking.

Here, we show that selective downregulation of *htt* in the sLNvs impairs the robustness of free-running circadian rhythms and increases sleep in female flies. We further demonstrate that sleep phenotypes are not developmental in origin, as adult-specific *htt* knock-down recapitulates the behavioural effects. To link these behavioural changes to underlying cellular mechanisms, we examined DCV trafficking within sLNv axons and found that *htt* downregulation alters both the proportion of motile vesicles and their transport dynamics, while abolishing normal time-of-day-dependent changes in vesicle trafficking. Finally, whole-cell patch-clamp recordings revealed that *htt* knock-down reduces action potential firing frequency without affecting resting membrane potential. Together, these findings identify huntingtin as an important cell-endogenous regulator of DCV transport, neuronal excitability, and behavioural output in a defined neuronal population.

## RESULTS

### Downregulation of *htt* in sLNvs impairs free-running circadian rhythmicity

Unlike in mice, where HTT LOF mutants are embryonically lethal (25–27), adult *Drosophila* with *htt* LOF mutations remain viable (12), suggesting that *htt* functions predominantly in adulthood rather than during development. Since *htt* is ubiquitously expressed in the fly brain, in this work we opted for an RNA interference (RNAi) approach using the UAS-Gal4 system (28) to achieve cell-specific downregulation, rather than relying on genetic mutants. This strategy enables the dissection of cell-autonomous effects of *htt* from network-driven phenotypes, with the aim of providing a more detailed understanding of its function. We first obtained two UAS-*htt*^RNAi^ lines: UAS-*htt*^RNAi-TRiP.JF01205^ (29) and UAS-*htt*^RNAi-36204GD^. To assess their efficacy for *htt* downregulation, we determined mRNA levels by qPCR in flies expressing each RNAi ubiquitously under the Act5C-Gal4 driver, using the corresponding heterozygous UAS-*htt*^RNAi^ line (without driver) as genotype-matched control. Both RNAi lines reduced *htt* mRNA levels relative to their respective controls; however, only Act5C-Gal4>UAS-*htt*^RNAi-TRiP.JF01205^ showed a statistically significant reduction (∼67%, p<0.05), whereas the knock-down observed with Act5C-Gal4>UAS-*htt*^RNAi-36204GD^ did not reach significance (**Supplementary Figure 1A**). Based on this result, we selected UAS-*htt*^RNAi-TRiP.JF01205^ for subsequent experiments, hereafter referred to as UAS-*htt*^RNAi^.

Functional sLNvs are important to maintain circadian rhythmicity under free-running conditions (19,22). We therefore hypothesised that, if *htt* plays a physiological role in sLNvs, its downregulation may cause circadian disruptions. We utilized the R6-Gal4 enhancer trap line (30) for sLNvs targeted *htt* downregulation. The driver specificity was first validated by co-localizing R6-Gal4-driven nuclear Green Fluorescent Protein (nGFP) with anti-PDF immunostaining (**Supplementary Figure 1B**). Male progeny from R6-Gal4 x UAS-*htt*^RNAi^ crosses were entrained to light:dark (LD) cycles and subsequently transferred to constant darkness (DD) to assess free-running locomotor activity using *Drosophila* Activity Monitors (DAMs, Trikinetics). Heterozygous controls resulting from crosses of R6-Gal4 or UAS-*htt*^RNAi^ to *w^1118^* flies, which is the genetic background of these fly stocks, were used. Downregulation of *htt* in sLNvs causes a decrease in the circadian rhythmicity of locomotor activity as it is appreciated in representative actograms (**Figure 1A**) and periodograms (**Figure 1B**). The free-running population rhythmicity of flies with *htt* knock-down in sLNvs was 59.55%, which was lower than in the genetic controls (86.52% in the UAS-*htt*^RNAi^ heterozygous control and 71.43% in the R6-Gal4 heterozygous control, **Supplementary Table 1**). To assess circadian rhythmicity at the level of individual animals, rather than only at the population level, we calculated the difference between the peak power value and its significance threshold (31). This analysis revealed a statistically significant decrease in this parameter in the flies with sLNvs specific-*htt* downregulation compared to genetic controls (**Figure 1C-D**). The free running-period was not altered by the manipulation (**Figure 1E**), which is consistent with an alteration of the sLNvs output, rather than a disruption of the molecular circadian clock mechanism. Overall, this experiment suggests that *htt* plays a functional role in sLNvs.

**Figure 1.**
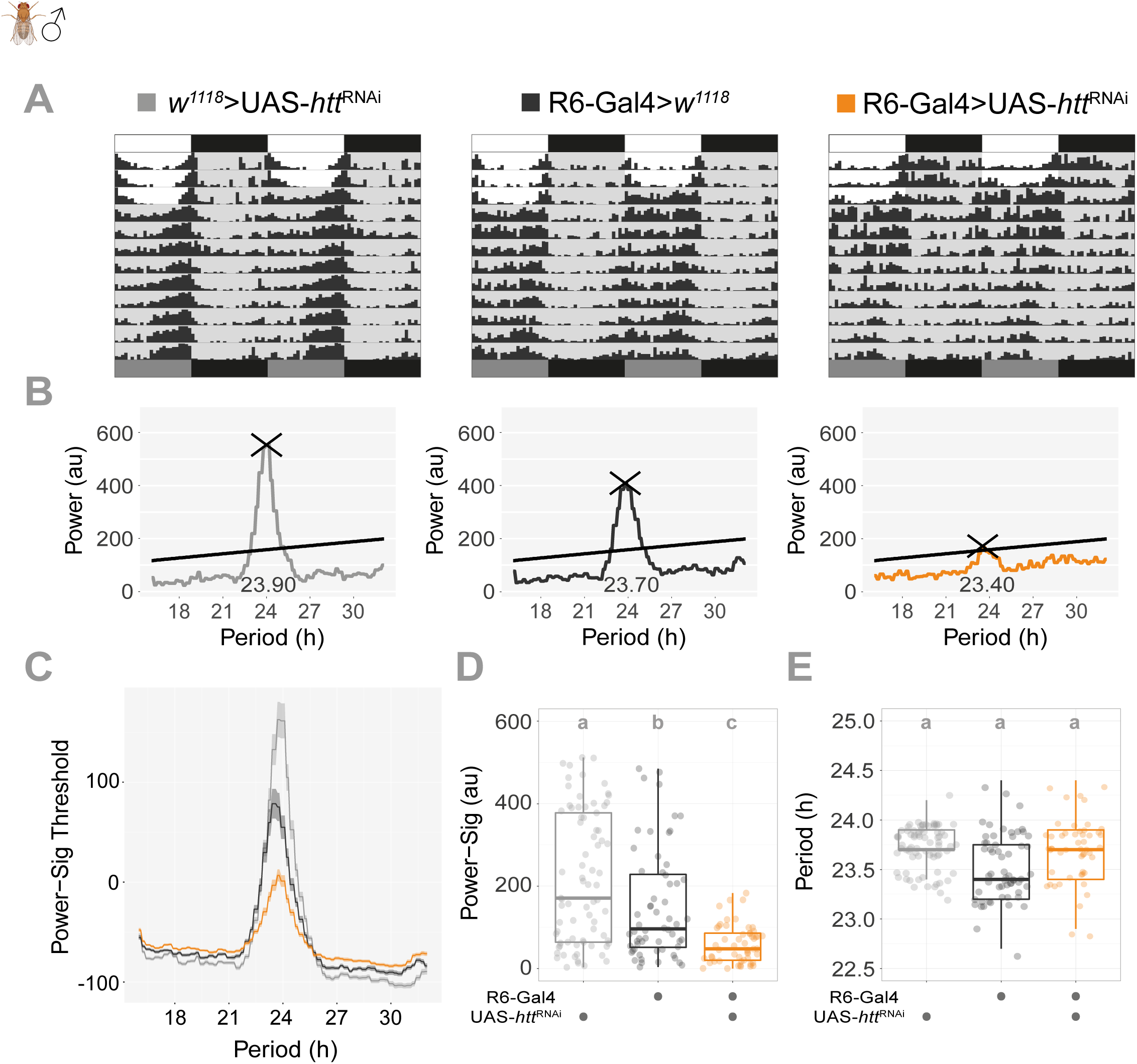
Knock-down of *htt* in sLNvs results in reduced circadian rhythm robustness. **A** Representative actograms of control (*w^1118^*>UAS-*htt*^RNAi^ and R6-Gal4>*w^1118^)* and experimental (R6-Gal4>UAS-*htt*^RNAi^) male flies under constant darkness. Gray shading indicates subjective night. **B.** Individual periodogram analysis corresponding to the representative flies shown in (A). The solid line indicates the significance threshold (*p<0.05*). **C.** Population periodograms showing the average rhythmic power-significance distribution across genotypes. **D.** Boxplots of individual power-significance values showing a significant reduction in circadian rhythm strength in *htt* knockdown flies. **E.** Circadian period in hours for rhythmic individuals. Total sample sizes are n=84-89 flies per genotype from N=3 independent biological replicates. Different letters denote statistically significant differences (*p<0.05*, One-way ANOVA followed by Tukey’s post hoc test).

### Downregulation of *htt* in sLNvs increases sleep

sLNvs are clock neurons involved in sleep regulation, as changes in their physiology can influence not only the timing but also the quantity and quality of sleep (21). To test if endogenous *htt* is involved in the control of sleep by these neurons, we used RNAi mediated downregulation using the R6-Gal4 driver and quantified sleep behaviour using DAMs under LD conditions. Locomotor activity was recorded as infrared beam breaks per minute, with sleep defined by the conventional five-minute immobility criterion (32,33). Downregulation of *htt* in sLNvs produced a marked increase in sleep in female flies (**Figure 2A-B**). The activity index, a measure of how active flies are during the active periods, was not decreased in the experimental group (**Figure 2C**). Assessing the activity index is essential when sleep is inferred from locomotor activity, as an inverse relation between sleep and activity may reflect a spurious increase in sleep arising from underlying motor impairments, rather than true sleep changes. Because the activity index was actually increased in the experimental group, the observed increase in sleep cannot be attributed to reduced locomotion. While quantitatively the sleep increase was present both during the daytime (siesta sleep) and at nighttime (**Figure 2A, D, H**), there were some differences in the effect of *htt* downregulation over the sleep architecture. For instance, the latency to the first daytime sleep episode was shorter on average in the experimental group compared to controls, whereas no differences were observed in the latency to the first nighttime sleep episode (**Figure 2E, I**). Sleep bout analysis showed that the sleep increase was due to longer sleep bout duration throughout the diel cycle (**Figure 2F, J**). Although *htt* knock-down did not affect the number of sleep bouts during daytime sleep (**Figure 2G**), it significantly reduced them at night (**Figure 2K**), consistent with more consolidated nighttime sleep. All behavioural parameters are shown in **Supplementary Table 2**.

**Figure 2.**
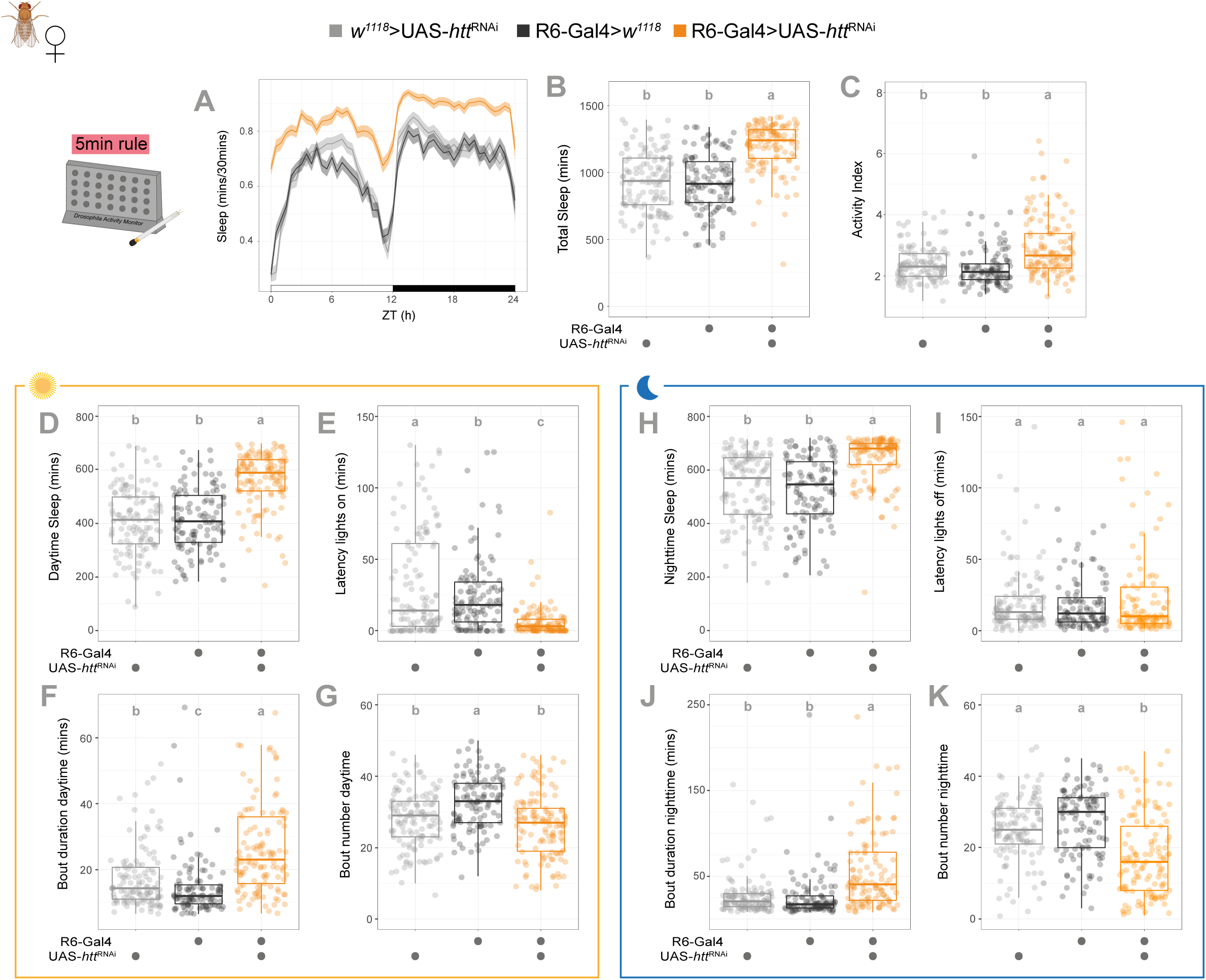
Constitutive downregulation of *htt* in sLNvs increases sleep. **A.** Sleep ethogram. Temporal distribution of sleep in female flies, quantified in 30-minute bins over a 24-hour Light:Dark cycle. White and black bars indicate day (ZT0–12) and night (ZT12–24), respectively. Shaded areas represent the standard deviation. **B.** Total sleep. Boxplot showing the total amount of sleep (minutes) over the 24-hour period. **C.** Activity index. Boxplot showing the average activity per waking minute, indicating that the sleep phenotype is not due to a general decrease in locomotor capacity. **D–G**. Daytime sleep parameters. Quantification of (D) total daytime sleep duration, (E) sleep latency (time to first sleep bout after lights on), (F) mean sleep bout duration, and (G) sleep bout number during the light phase (ZT0–12). **H–K.** Nighttime sleep parameters. Quantification of (H) total nighttime sleep duration, (I) sleep latency (time to first sleep bout after lights off), (J) mean sleep bout duration, and (K) sleep bout number during the dark phase (ZT12–24). Data were recorded using the DAM system, with sleep defined as 5 minutes of continuous inactivity. Sample sizes: n=113-125 flies per genotype, N=5 independent experiments. Different letters indicate statistically significant differences (*p<0.05*) using GLMMs and Sidak’s post-hoc analysis.

In male flies, sleep parameters were not affected by *htt* downregulation in the sLNvs (**Supplementary Figure 2A-C and Supplementary Table 3**), suggesting that female sleep is more sensitive to reduced *htt* expression. We propose that male sleep is less sensitive to this manipulation, rather than male sLNv function being unaffected. This is supported by our circadian behaviour assays, which show that males exhibit a significant reduction in the strength of free-running locomotor rhythms (**Figure 1**), confirming that *htt* knock-down does indeed alter male sLNv function. The observed sexual dimorphism in sleep regulation is consistent with previous reports indicating that sLNvs are integrated into sexually dimorphic neuronal circuits (34–36). Collectively, these results identify endogenous *htt* in the sLNvs as an important regulator of sleep, particularly in female flies.

### Adult-specific knock-down of *htt* in sLNvs is sufficient to increase sleep

To test whether the increase in sleep following *htt* knock-down in sLNvs was due to a developmental effect, we downregulated *htt* in sLNvs exclusively after development, using the Auxin-inducible Gene Expression System (AGES). This genetic tool works by exploiting the plant hormone auxin to trigger the degradation of an auxin-inducible degron-tagged Gal80 repressor, thereby allowing Gal4-mediated transgene expression of the *htt*^RNAi^ only upon auxin administration (37). Adult-specific *htt* knock-down was achieved by combining R6-Gal4 with the AGES system and UAS-*htt*^RNAi^. Experimental flies were exposed to auxin for five days before behavioral assays, together with genetic and vehicle-treated controls. Downregulation of *htt* restricted to adulthood led to an increase in sleep (**Figure 3A-F, Supplementary Table 4**). Notably, the increase in activity index was also present in the adult-specific manipulation (**Figure 3C**). The persistence of these phenotypes following adult-specific manipulation supports an ongoing functional role for *htt* in the adult nervous system.

**Figure 3.**
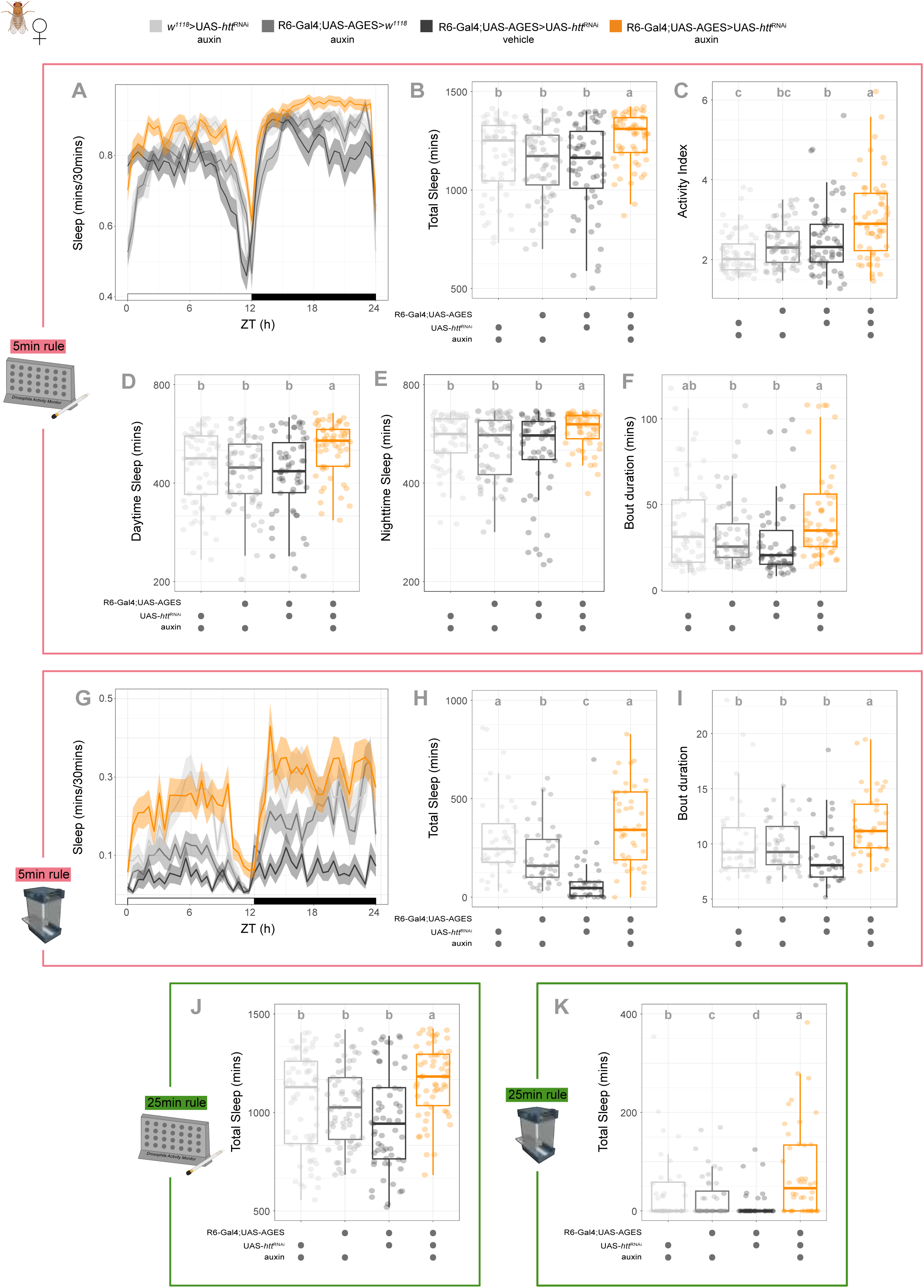
Adult-specific knock-down of *htt* in sLNvs consistently increases sleep across different experimental platforms and sleep definitions. Adult-specific downregulation of *htt* using the AGES systems recapitulated the increased sleep phenotype, demonstrating that the effect is not due to developmental defects. **A–F** Sleep analysis using the Drosophila Activity Monitor (DAM) system and the conventional 5-min sleep definition. **A**. Sleep ethograms. **B**. Total 24-h sleep. **C**. Activity index. **D**. Daytime sleep. **E**. Nighttime sleep. **F**, Mean sleep bout duration over 24 h. **G–I** Sleep analysis using the Ethoscope video-tracking platform and the 5-min sleep definition. **G**. Sleep ethograms. **H**. Total 24-h sleep. **I**. Mean sleep bout duration. **J–K** Total 24-h sleep quantified using a more stringent 25-min inactivity criterion in the **J.** DAM system and **K**. Ethoscope platform. Sample sizes for DAM system (A–F, J): n=59–67 flies per genotype, N=3 independent experiments. Sample sizes for Ethoscopes (G–I, K): n=82–88 flies per genotype, N=3 independent experiments. Different letters indicate statistically significant differences (*p<0.05*) using GLMMs and Sidak’s post-hoc analysis.

To ensure accurate and reliable quantification of sleep, and to avoid the overestimations often associated with traditional beam-crossing systems, we employed ethoscopes, an automated video-tracking platform specifically designed for high-resolution monitoring of *Drosophila* behaviour (38). This system allowed us to detect sleep episodes with high temporal precision and to characterise sleep architecture faithfully. Additionally, differences in basal locomotor activity between *htt* knockdown flies and parental controls, as measured by the DAM system, warranted the use of a more reliable method for sleep inference. When we analysed sleep patterns in adult-specific RNAi knockdown flies targeting sLNvs’ *htt*, we observed a trend towards an increase in total sleep, particularly in females (**Figure 3G-H, Supplementary Table 5**). Using this approach, we found a significant increase in the length of individual sleep bouts in females (**Figure 3I**), with this effect being especially pronounced during the night phase (**Supplementary Table 5**). In addition to the prolonged sleep bouts, we noted a reduction in lights-on sleep latency in the *htt* RNAi-induced flies (**Supplementary Table 5**).

Prompted by these findings, and based on previous work which proposes that long sleep bouts (∼25 minutes) are key for discharging sleep pressure and thus serve as an important target of sleep homeostasis (39), we reanalysed our data using this criterion. This refined analysis revealed more pronounced alterations in sleep parameters in female flies with reduced *htt* expression in the sLNvs when only longer sleep bouts were considered (**Figure 3J, Supplementary Table 6**), suggesting that *htt* knock-down preferentially affects physiologically relevant, deeper sleep. When ethoscope data were analyzed using the 25-minute bout threshold, *htt* knockdown flies exhibited a statistically significant increase in sleep (**Figure 3K, Supplementary Table 7**), which suggests a possible alteration in sleep homeostasis or sleep need.

Consistent with the constitutive knock-down, adult-restricted *htt* downregulation in male flies failed to elicit statistically significant changes in sleep when assessed with both DAMs (**Supplementary Figure 3A-D, Supplementary Tables 8** and **9**) and ethoscopes (**Supplementary Figure 3E-G, Supplementary Tables 10** and **11**). For completeness and to facilitate comparison of effects across manipulations, we also include long-bout sleep analysis for the constitutive manipulations (**Supplementary Figure 4** and **Supplementary Figure 2D, Supplementary Tables 12** and **13**). Taken together, these experiments support a functional role for *htt* in sLNvs, as its downregulation results in an increase in sleep, particularly the more restorative long-episode sleep. Additionally, we confirmed that this potential role for *htt* in the regulation of sleep is specific to female flies.

### Downregulation of *htt* affects dense-core vesicle axonal transport

To investigate the mechanism by which *htt* modulates sLNv-driven behaviors, we quantified the levels of PDF, the principal neuropeptide expressed by these neurons (19,22–24). PDF levels at sLNv axonal terminals exhibit pronounced daily dynamics, peaking in the morning and declining at night (40). Moreover, PDF is considered a wake-promoting signal, as genetic disruption of the coding gene or its receptor is associated with increased sleep (41). Based on this arousal role and the sleep increases observed upon *htt* downregulation, we hypothesized that reduced *htt* expression would lead to decreased PDF immunoreactivity at sLNv axonal terminals.

To test this, we quantified PDF immunoreactivity at *Zeitgeber* Time (ZT) 2 in flies with constitutive *htt* knock-down in the sLNvs and compared it with genetic controls. Visual inspection revealed no gross differences between genotypes; that is, *htt* knock-down did not affect the number of sLNv somata, nor the overall organisation of their projections or axonal terminals. This lack of structural alteration is consistent with a primarily post-developmental effect of the manipulation, as developmental disruptions of PDF signaling are known to produce lasting changes in adult sLNv morphology (42). We quantified the intensity of PDF staining in the sLNvs axonal terminals and found no differences in the *htt* downregulated group (**Figure 4A, Supplementary Table 14**), which was unexpected.

**Figure 4.**
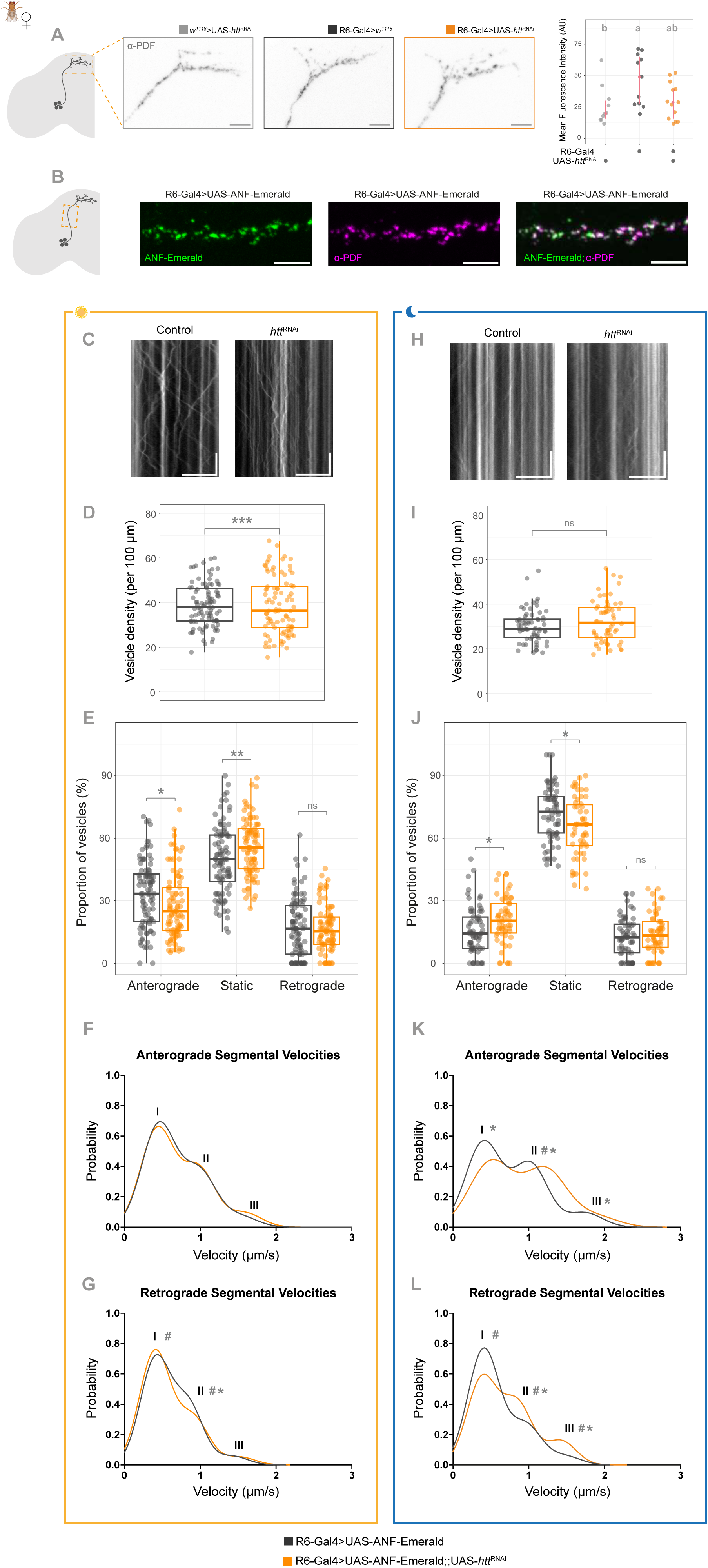
Downregulation of *htt* in sLNvs impairs axonal dense core vesicle transport dynamics without affecting PDF overall levels. **A**. Representative confocal images (left) of sLNvs dorsal axonal projections stained with anti-PDF at ZT2. Quantification (right) of total PDF immunoreactivity (mean fluorescence intensity) at the terminals, showing no significant differences between controls and experimental flies. n=10-13, N=4 independent experiments. Different letters denote statistically significant differences (*p<0.05*, One-way ANOVA followed by Tukey’s post hoc test). Scale bars: 10μm. **B.** Validation of the DVC reporter. Representative confocal images of sLNvs axons expressing UAS-preproANF-Emerald (green) and immunostained for anti-PDF (magenta) at ZT2. Scale bars: 5µm. The merged image (right) shows a highly overlapping distribution of both signals, indicating that the ANF::Emerald reporter effectively labels the PDF-containing secretory compartment in these neurons. **DVC transport dynamics during the Light Phase (ZT1–6)**: **C.** Representative kymograph of vesicle trafficking in sLNvs axons during the day. **D.** Vesicle density (vesicles per 100μm), showing no significant changes in total cargo number. **E.** Vesicle population distribution. Percentage of vesicles classified as anterograde, stationary, or retrograde. **F–G.** Segmental Velocities. Frequency distribution and triple-Gaussian fitting representing slow (I), middle (II), and fast (III) modes of moving vesicles. (F) Anterograde and (G) Retrograde segmental velocities during the day. **DVC transport dynamics during the Dark Phase (ZT13–18)**: **H.** Representative kymograph of vesicle trafficking in sLNvs axons during the night. **I.** Vesicle density (vesicles per 100μm), showing no significant changes in total cargo number. **J.** Vesicle population distribution. Percentage of vesicles classified as anterograde, stationary, or retrograde during nighttime. **K–L.** Segmental Velocities. Frequency distribution and triple-Gaussian fitting of representing slow (I), middle (II), and fast (III) modes of moving vesicles. (K) Anterograde and (L) Retrograde segmental velocities during the night. Asterisks indicate significant differences derived from hierarchical GLMMs (with fly and hemisphere as nested random effects). Asterisks (Mode centre) and numeral symbols (Mode fraction) indicate significance in panels F, G, K and L. \*\*\**p<0.001,* \*\**p<0.01*, \**p<0.05*; ns, non-significant (*p>0.05*). For experiments shown in panels C to L genotypes are control (*R6-Gal4>UAS-preproANF-Emerald*) and experimental (*R6-Gal4>UAS-preproANF-Emerald;;UAS-htt*^RNAi^). Kymograph scale bars: vertical 5 seconds, horizontal 10μm.

Given the lack of detectable changes in this static snapshot of steady-state PDF immunofluorescence signal at sLNv terminals, we turned to a dynamic approach that allowed us to assess the impact of *htt* on neuropeptide vesicle trafficking. It has been reported that wild-type huntingtin promotes axonal vesicular transport (43). This is mediated through interactions with huntingtin-associated protein 1 (HAP1) and the p150 (Glued) subunit of dynactin, in a complex that enhances the transport of vesicles filled with the neuropeptide Brain Derived Neurotrophic Factor (43). In *Drosophila*, *htt* LOF leads to synaptic bouton trafficking defects in larval motor neurons (17,18,44), suggesting that the role of huntingtin in supporting vesicle dynamics is evolutionarily conserved. We therefore hypothesized that *htt* downregulation may affect the axonal transport of DCVs in sLNvs. To test this we expressed the UAS-preproANF-Emerald construct (45) that serves as a fluorescent marker of DCVs. This line enables cell-specific expression of the rat neuropeptide atrial natriuretic factor (ANF) fused to Emerald GFP, and has been used to visualize DCVs in *Drosophila* before (46,47).

To confirm that the ANF::Emerald fusion protein labels PDF-containing DCVs, we drove its expression in sLNvs using the R6-Gal4 driver and validated this by anti-PDF immunostaining. Confocal analysis showed that ANF::Emerald fluorescence largely overlapped with α-PDF immunostaining in punctate structures along sLNv axons, suggesting that both proteins occupy the same subcellular compartment (**Figure 4B, Supplementary Figure 5A**). Expansion microscopy has revealed that individual DCVs within sLNvs could carry more than one type of neuropeptide (72). However, given that the diameter of DCVs (80-200 nm) approaches the diffraction limit of conventional confocal microscopy (∼200 nm), the observed overlap should be interpreted as co-localization rather than definitive evidence of co-packaging. In addition to overlapping fluorescent signals, some ANF::Emerald-positive and PDF-positive fluorescent structures remained spatially distinct, suggesting that a fraction of each cargo is present in separate vesicular populations. At the behavioral level, ANF::Emerald expression alone slightly reduced overall sleep; nevertheless, constitutive *htt* downregulation in flies co-expressing ANF::Emerald still recapitulated the expected increase in sleep (**Supplementary Figure 5B-C** and **Supplementary Table 15**). This indicates that ANF::Emerald expression, required to visualize DCV dynamics in live tissue, did not interfere with our ability to assess the effects of *htt* downregulation on vesicle transport.

To quantify DCV dynamics, we used brain explants of female R6-Gal4>UAS-*htt*^RNAi^; UAS-preproANF-Emerald flies and compared them to R6-Gal4>UAS-preproANF-Emerald as controls. For this analysis, we focused on the ascending axonal projections of the sLNvs rather than on their terminal regions. This choice allowed us to quantify both anterograde and retrograde transport, which would not be feasible at the axonal terminals, where the complex and disorganized projection pattern makes it difficult to distinguish vesicles moving toward the terminal from those returning toward the soma. Sixty-second movies were generated to extract the dynamics of fluorescent ANF vesicles (**Movies 1, 2**) and transformed to kymographs (**Figure 4C, H**) from which trajectories were tracked semiautomatically using a custom-made MATLAB application (48). To capture time-of-day differences, experiments were carried out in the morning (ZT1–6) and during the early night (ZT13–18). ANF::Emerald DCVs were identified moving anterogradely (right descending trajectories), retrogradely (left descending trajectories) or stationary (vertical trajectories) within kymographs. The density of DCVs in sLNv axons was lower in *htt*^RNAi^ than in controls (**Figure 4D**). Moreover, dynamic analysis revealed that *htt* knock-down significantly reduced the proportion of anterograde vesicles while increasing the fraction of stationary vesicles relative to controls (**Figure 4E**), indicative of impaired axonal vesicle movement and a marked defect in DCV axonal flux (**Supplementary Table 16**). Next, the anterograde and retrograde segmental velocities of ANF::Emerald vesicles were extracted from control and *htt* knockdown movies and plotted following a multimodal distribution with a 3-gaussian mixture model, which can be associated with the number of active processive motors in a competition-cooperation model (49). When applied to the mobile vesicle population for each genotype and direction, this approach estimates the probability of encountering vesicles moving at slow (peak I), intermediate (peak II) or fast (peak III) segmental velocities. Each peak reflects variable configurations of active anterograde and retrograde motors that convey the movement of cargos at slower or faster velocities. This analysis revealed that, during the day, *htt* knock-down did not affect anterograde segmental velocities (**Figure 4F**), but increased the retrograde I mode fraction and reduced both the retrograde II mode centre and fraction, collectively indicating a decrease in overall retrograde transport modes (**Figure 4G, Supplementary Table 17**).

We then examined vesicle transport dynamics in the same experimental groups during the night phase (ZT13–18). Overall, both vesicle density and the dynamic properties of ANF::Emerald were reduced during this period relative to early morning. Contrary to daytime observations, total vesicle densities did not differ significantly between *htt* knock-down and control conditions (**Figure 4I**). Analysis of vesicle dynamics at night revealed the opposite pattern to that observed during the day: control flies displayed a lower proportion of anterograde vesicles and a higher proportion of stationary vesicles than *htt* knockdown flies (**Figure 4J**). When segmental velocities were compared using the 3-gaussian mixture model, we observed significant changes in anterograde and retrograde dynamics towards lower I mode fractions and faster mode centres suggesting overall increases in velocities after *htt* knock-down compared to control during the night (**Figure 4K, L, Supplementary Table 17**).

Taken together, our DCV axonal transport experiments reveal that control flies are able to modulate axonal transport of DCV within different times of the day since the proportion of anterograde particles drops more than two-folds from ∼33% during the day to ∼15% at night. This day-night difference was largely abolished by *htt* knock-down, as anterograde vesicle fractions were only slightly higher during the day than at night (∼27% and ∼22%, respectively). This distinction is particularly meaningful given that sLNvs encode time-of-day information through temporally regulated neuropeptide release, with greater output in the morning than at night. Reducing *htt* expression therefore appears to compromise the daily modulation of sLNv signaling capacity to downstream circuits.

### Downregulation of *htt* disrupts neuronal activity

Axonal vesicle transport and neuronal electrical activity are tightly interconnected and mutually dependent processes. Neuronal activity regulates the machinery responsible for vesicle transport, recycling, and degradation (50), whereas the targeting of voltage-gated ion channels to specific axonal domains that shape neuronal excitability depends on vesicular transport mechanisms (51). In the context of HD models, there have been reports of several electrophysiological alterations, in different neuronal types and contexts (52–54). However, since those models were affecting large populations of neurons, it is difficult to discern cell-endogenous from circuital mechanisms. Having established that *htt* knock-down disrupts vesicular trafficking, we sought to determine whether neuronal activity was concomitantly affected in our system, in which the genetic manipulation is restricted to a small subset of neurons.

We performed whole-cell patch-clamp recordings in brain explants following previously established methods (55). For this experiment, we recorded from the lLNvs, which are larger than the sLNvs and therefore better suited to electrophysiological approaches. Similar to sLNvs, the lLNvs function as arousal neurons and signal via the expression of the PDF neuropeptide (56). We downregulated *htt* using RNAi in LNvs using the *Pdf*-Gal4 driver combined with a *Pdf*-RFP element, where the *Pdf* gene promoter directly drives the expression of the Red Fluorescent Protein (57) and allows the visualisation of the target cells. As control we used an heterozygous parental line corresponding to the line *Pdf*-Gal4; *Pdf*-RFP crossed to *w^1118^*. To determine whether *htt* downregulation affects the electrical properties of lLNvs, we performed whole-cell patch-clamp recordings from identified neurons during the daytime and measured their spontaneous membrane activity.

Our electrophysiological recordings show that lLNvs exhibit their characteristic membrane oscillations, a hallmark of neuropeptidergic neurons, and that *htt* knock-down does not significantly alter the organisation of action potential firing (**Figure 5A**). Membrane potential, measured at inter-burst troughs, was similarly unaffected (**Figure 5B**). In contrast, action potential firing frequency was significantly reduced upon *htt* downregulation (**Figure 5C**). Burst frequency, which depends on cholinergic afferent input from the optic lobes rather than cell-autonomous mechanisms (58–60), showed no significant differences following *htt* knock-down, as would be anticipated for an externally driven parameter (**Figure 5D**). These results indicate that *htt* is required to sustain the high action potential firing rates characteristic of daytime LNvs activity. The absence of any effect on the inter-burst membrane potential may argue against a significant role for htt-mediated transport in the delivery of leak channels that set resting conductance. However, the reduction in firing rate following *htt* knock-down could suggest that the trafficking of voltage-gated conductances may be compromised.

**Figure 5.**
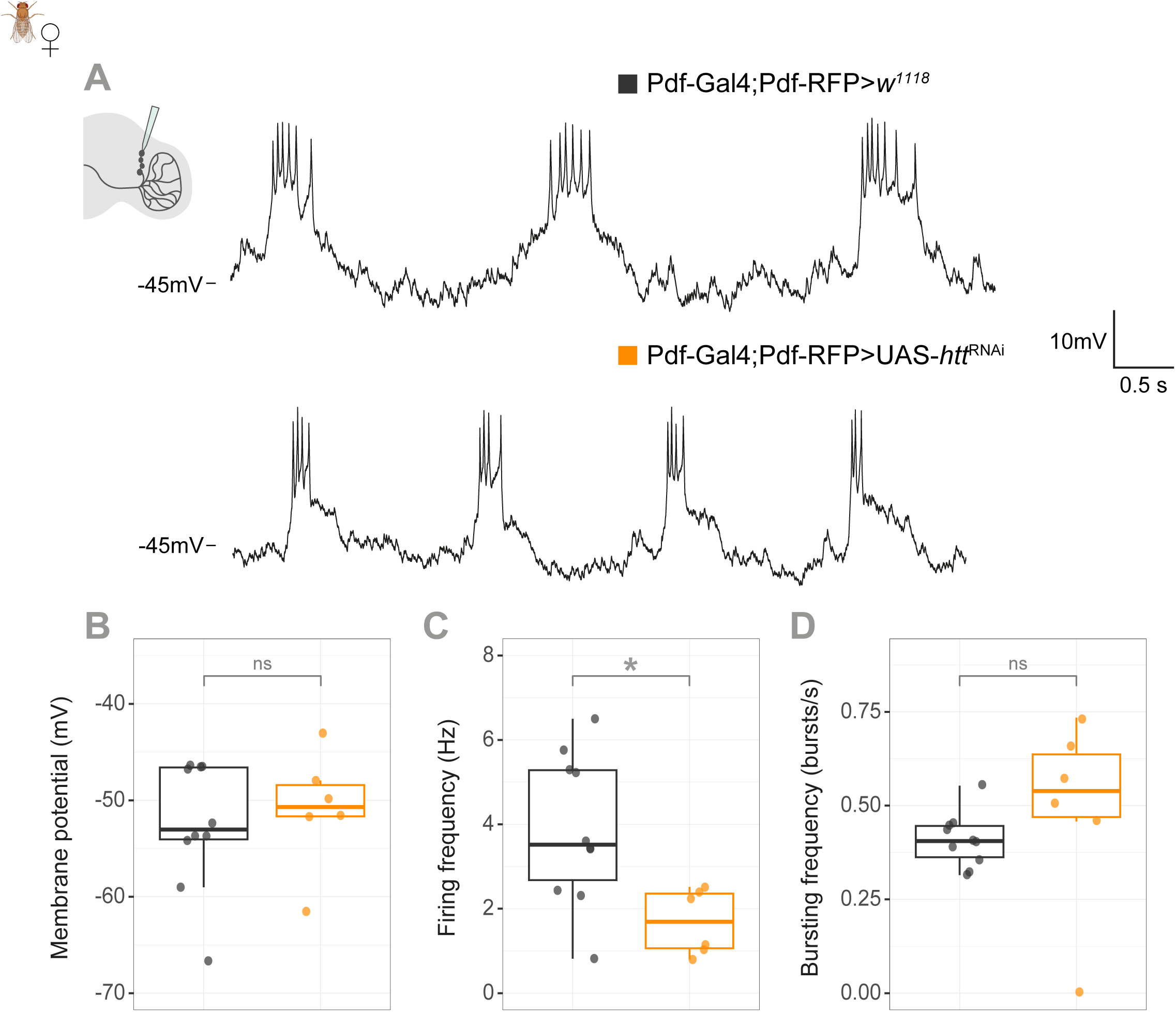
Whole-cell patch-clamp recordings reveal changes in the electrical properties of lLNvs upon *htt* downregulation. **A.** Representative voltage traces of control (*Pdf*-Gal4;*Pdf*-RFP>*w^1118^*) and experimental (*Pdf*-Gal4;*Pdf*-RFP>UAS-*htt*^RNAi^) lLNvs. Neurons were identified by RFP expression and recorded in current-clamp mode. **B.** Membrane potential measured as the trough between bursts, quantified in millivolts (mV), mean values ± SD are: control −52.6 ± 6.6 mV; experimental −50.9 ± 6.1 mV. **C.** Action potential firing frequency, mean values ± SD are: control 3.88 ± 1.79 Hz; experimental: 1.69 ± 0.77 Hz. **D.** Bursting frequency, quantification of the frequency of firing clusters (bursts per second) mean values ± SD are: control 0.41 ± 0.07 Hz; experimental: 0.49 ± 0.26 Hz. All recordings were performed between ZT4 and ZT12. To ensure consistency across samples, all quantifications represent the one-minute interval starting 26 minutes post-dissection. Sample sizes: control n=9, experimental n=6. Statistical significance was determined by Welch’s two-sample T-test (\**p<0.05*; ns, non-significant *p> 0.05*).

## DISCUSSION

Our findings contribute to two important areas of huntingtin research. First, they address physiological functions of endogenous huntingtin, a topic of particular relevance given that many HD therapeutic strategies currently under investigation aim to reduce *huntingtin* expression without fully discriminating between the mutant and the wild-type protein (61). A better understanding of the physiological roles of huntingtin is therefore essential for anticipating the consequences of such approaches. Second, our study employs cell type-specific downregulation of *huntingtin* in a restricted neuronal population, allowing the examination of its functions with minimal interference from the widespread circuit-level adaptations and compensatory mechanisms that often accompany broader genetic manipulations. In mammals, cell type-specific downregulation of *huntingtin* has emerged as an important area of investigation and has provided valuable insights into its roles in processes such as synaptic connectivity, development and neuronal survival (62–66). Nevertheless, most studies to date have manipulated relatively large neuronal populations or broad brain regions, limiting the resolution at which huntingtin function can be dissected within defined sparse populations. Therefore, our study sets the basis for the cell-autonomous exploration of the roles of wild-type huntingtin, demonstrating that its downregulation in only 8 neurons of an otherwise functional brain can lead to discernible behavioural phenotypes.

### *Drosophila htt* and behavioural control

Sleep and circadian rhythm disturbances are common features of HD and contribute substantially to disease burden and reduced quality of life (67,68). Despite this, relatively little is known about the contribution of huntingtin loss-of-function to the regulation of sleep and circadian rhythms, since studies exploring these phenotypes have focused primarily on the toxic gain-of-function effects associated with the mutant protein. Here we show that selective *huntingtin* knock-down in the sLNvs reduces the robustness of free-running locomotor activity rhythms, while leaving the free-running period unchanged. This finding is consistent with a role for huntingtin in mediating neuronal output pathways downstream of the circadian clock, rather than in the core molecular mechanisms that underlie circadian timekeeping.

One of the few studies addressing huntingtin loss-of-function in sleep regulation found that pan-neuronal downregulation of huntingtin in *Drosophila* produces a less consolidated sleep (69). In our study, selective *huntingtin* knock-down in the sLNvs resulted in a contrasting phenotype, with animals exhibiting more consolidated sleep and an overall increase in sleep time, most prominently in the longer sleep episodes that are considered more restorative. Importantly, our findings were replicated across two independent sleep-monitoring platforms and were also observed following adult-specific huntingtin knock-down, demonstrating that huntingtin contributes to sleep regulation independently of its developmental functions.

Notably, the sleep phenotype was restricted to females. Sleep measurements obtained with beam-crossing systems can underestimate activity and potentially produce ceiling effects in male flies. However, we also failed to detect sleep phenotypes in males using the ethoscope platform, which provides high-resolution behavioural tracking and is not subject to these limitations. Together, these findings underscore the importance of considering sex as a biological variable in sleep research and demonstrate that marked sex-specific effects can arise even following genetic manipulations of neuronal populations that are not overtly sexually dimorphic.

### *Drosophila htt*, neuropeptides and DCV axonal transport

Huntingtin functions as a scaffold protein implicated in the trafficking of a wide range of axonal cargoes, including synaptic vesicle precursors, endosomes, autophagosomes, and DCVs, the type of vesicle responsible for neuropeptide transport and release (8). Because the sLNvs exert much of their influence through neuropeptidergic signaling, they represent a particularly relevant neuronal population in which to investigate this function of huntingtin. sLNvs neurons express several neuropeptides, most notably PDF (19,22–24), but also short Neuropeptide F and *Drosophila* insulin-like peptide 2, both of which have also been implicated in sleep regulation (70,71). Notably, individual DCVs within the sLNvs may carry more than one type of neuropeptide (72).

Since huntingtin downregulation did not alter steady-state levels of PDF immunoreactivity, we reasoned that defects in DCV trafficking might occur without producing detectable changes in overall abundance of this specific neuropeptide. We therefore examined DCV axonal transport directly. To minimize interference with endogenous signaling pathways, we employed a fluorescent reporter based on the rat Atrial Natriuretic Factor peptide, an exogenous cargo for which no receptor exists in *Drosophila*. This strategy allowed us to visualize DCV dynamics while reducing the likelihood of perturbing the physiological functions of endogenous neuropeptide signalling. While expression of ANF-GFP has been reported to interfere with certain aspects of neuronal function when driven at high levels using a strong Gal4 driver (73), in our experiments, driving the ANF reporter with R6-Gal4 did not interfere with the increased-sleep phenotype induced by huntingtin knock-down (Supplementary Figure 5B-C), validating this tool to assess DCV transport in sLNvs under these conditions.

Early *in vivo* work in *Drosophila* larvae demonstrated that reduction of endogenous *huntingtin* disrupts fast axonal transport, leading to abnormal accumulations of vesicles and organelles within axons, indicating that normal huntingtin function is required for efficient microtubule-based trafficking (18). These findings support the concept that HD pathology may arise not only from toxic gain-of-function mechanisms associated with polyglutamine expansion, but also from impairment of the physiological transport functions of huntingtin. Loss of huntingtin disrupts the motility and distribution of Rab11-positive vesicles, supporting a role for huntingtin in Rab-dependent membrane trafficking pathways (74). These observations suggested that huntingtin functions as a scaffolding platform coordinating vesicular cargoes with molecular motor complexes during axonal transport. Huntingtin does not regulate all cargoes uniformly, but instead differentially controls the transport of distinct subsets of Rab-containing vesicles (75). Together, these studies support a model in which huntingtin functions as a central organizer of intracellular trafficking, coordinating kinesin– and dynein-dependent transport through selective interactions with Rab-defined vesicular populations (76).

A study in *Drosophila* larval neuromuscular junction reported that loss of *huntingtin* reduced retrograde DCV flux and altered their retention within synaptic boutons, leading to abnormal accumulation of neuropeptide-containing vesicles (17). Bulgari *et al.* concluded that native huntingtin acts as a regulator of DCV transport and neuropeptide storage without explicitly observing axonal dynamics (17). By contrast, using a more sensitive approach, the characterisation of DCV axonal transport within the adult fly brain, we uncovered multiple phenotypes affecting both transport speed and directionality. Given that huntingtin functions as a scaffold protein involved in the regulation of kinesin and dynein molecular motor interactions, analysis of DCV transport is particularly well suited to probe its neuronal role (76,77). We found that huntingtin plays crucial roles in regulating DCV day/night dynamic differences, since its knock-down reduces changes in anterograde to stationary vesicle percentages and significantly impair the anterograde and retrograde DCV segmental velocities compared with control.

A more recent study examined DCV dynamics at the sLNvs synaptic terminals and reported no differences in the velocity of anterograde or retrograde vesicles between ZT3 and ZT15 (78). In contrast, our analysis of DCV trafficking across comparable times revealed a higher proportion of anterograde-moving particles during the morning (ZT1–6), whereas nighttime recordings (ZT13-18) showed a marked increase in the proportion of stationary particles, which were not reported by Klose and colleagues (78). Several methodological differences may account for these apparently divergent findings. First, Klose *et al.* focused their analysis on synaptic terminals, where vesicle trafficking is regulated by both microtubule– and actin-based transport mechanisms (79). In contrast, we analyzed the ascending segment of the sLNv axons before they reach the terminal arborisation, a region in which transport is predominantly microtubule-dependent (80). This distinction may reveal different regulatory mechanisms operating along separate axonal compartments. Second, the approaches used to quantify vesicle movement differ substantially. Klose and colleagues reported mean DCV velocities, providing a global measure of transport dynamics. By contrast, our analysis was based on segmental velocities, allowing individual trajectories to be decomposed into shorter movement events. This approach offers greater resolution and enables the identification of distinct transport states, including stationary periods and movements likely driven by different numbers or combinations of molecular motors. Consequently, our analysis was able to detect changes in the relative distribution of transport behaviours that may not be apparent when considering only average velocity measurements. Here, we demonstrate that axonal DCV transport in the sLNvs undergoes day–night changes in the relative proportions of anterograde, retrograde, and stationary particles. Furthermore, proper *huntingtin* expression is required for these temporal dynamics, as huntingtin knock-down interferes with the normal cycling of axonal transport states in these neurons.

### *Drosophila htt* and neuronal physiology

Direct assessment of spontaneous neuronal activity in the LNvs using whole-cell patch-clamp electrophysiology revealed that *huntingtin* knock-down decreases action potential firing frequency, consistent with recent observations in mammalian systems (81,82). Notably, huntingtin has been implicated in the synchronisation within human cortical neuronal networks (83). Since the LNvs are part of a brain network characterized by coordinated membrane oscillations (58), the *Drosophila* system provides an attractive opportunity to investigate whether huntingtin also contributes to neuronal synchrony in an evolutionarily distant organism. Exploring this possibility lies beyond the scope of the present study but represents an interesting direction for future work.

LNvs are bursting neurons whose action potential firing is organized into rhythmic bursts driven by underlying membrane oscillations (58). The frequency of these oscillations is primarily regulated by cholinergic inputs (59). In this work, we observed a trend toward increased burst frequency in neurons with reduced huntingtin expression. Although this effect did not reach statistical significance, it is nevertheless intriguing and may point to altered cholinergic inputs signaling. While this possibility was not directly examined in the present study, huntingtin has previously been implicated in the trafficking and proper localisation of postsynaptic receptors (64,84). Thus, changes in the processing or distribution of cholinergic receptors could contribute to the potentially altered oscillatory frequency observed following *huntingtin* knock-down in LNvs.

Overall, our results are consistent with a bidirectional relationship between axonal transport and neuronal activity. On the one hand, neuronal activity is known to regulate cargo trafficking and capture since calcium-dependent signaling can arrest moving organelles at active sites (85–87), and synaptic activity can promote capture of transiting vesicles into resident presynaptic pools (47,88). Chronic silencing can also increase presynaptic DCVs accumulation (89). On the other hand, the literature directly supports transport defects as a cause of reduced neuronal function, because anterograde transport is required to supply presynaptic terminals with active-zone material, synaptic vesicle precursors, mitochondria, and ion channels (90,91). Thus, the reduced neuronal firing we observe could be a downstream consequence of impaired cargo delivery, while the altered activity state may in turn reinforce the shift from motile to stationary vesicle pools.

### Final conclusions

This study provides, to our knowledge, the first characterisation of huntingtin loss-of-function within a restricted neuronal population (8 neurons), enabling the identification of phenotypes that arise from cell type-specific reduction of *huntingtin* rather than from broader nervous system manipulations. At the same time, they contribute to the growing knowledge of sLNv function, a neuronal population that has served as a central model in *Drosophila* neuroscience but whose vesicular trafficking dynamics remain poorly understood. By linking huntingtin function to DCV transport and neuronal output in these cells, our work opens new avenues for investigating both the physiological roles of huntingtin and the mechanisms through which circadian neurons regulate behaviour. Our results convey with previous description that loss of huntingtin function disrupts axonal transport by impairing vesicle trafficking and altering bidirectional cargo movement leading to electrophysiological defects that ultimately impair clock output behaviours. Our findings support the hypothesis that loss of huntingtin physiological role plays a crucial regulatory role in HD pathology.

## MATERIALS AND METHODS

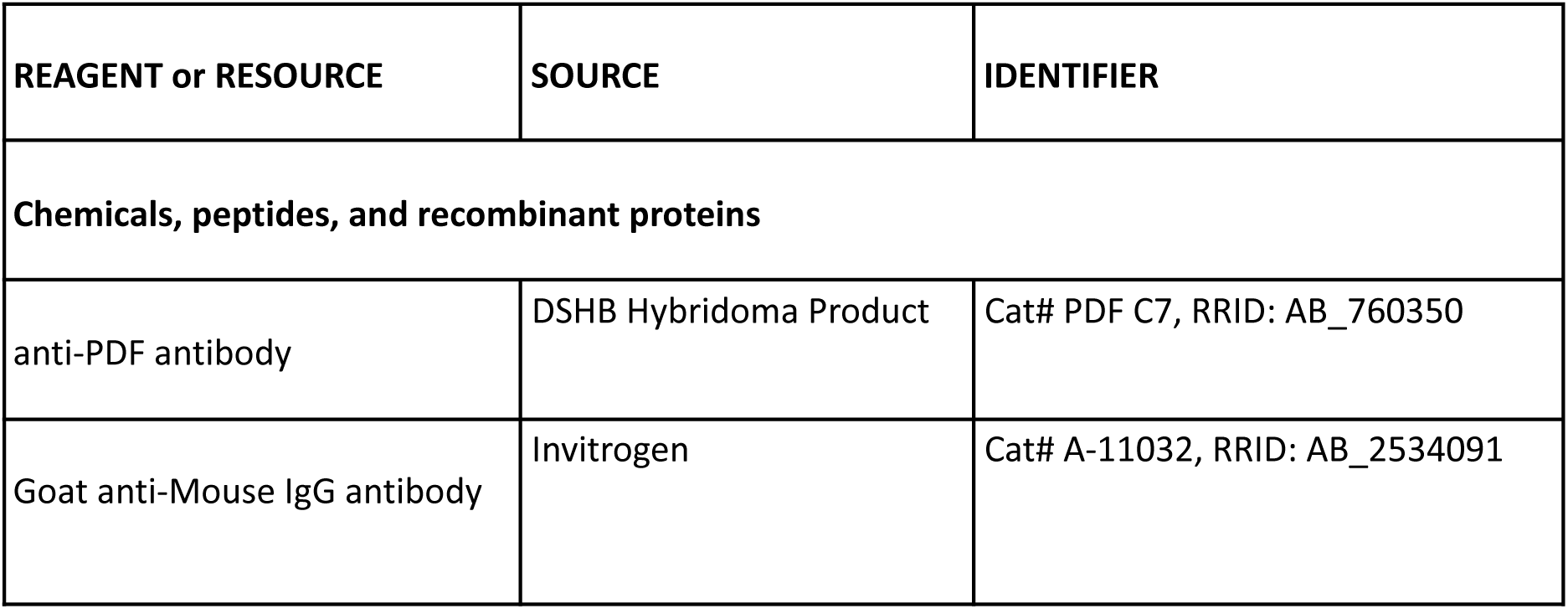
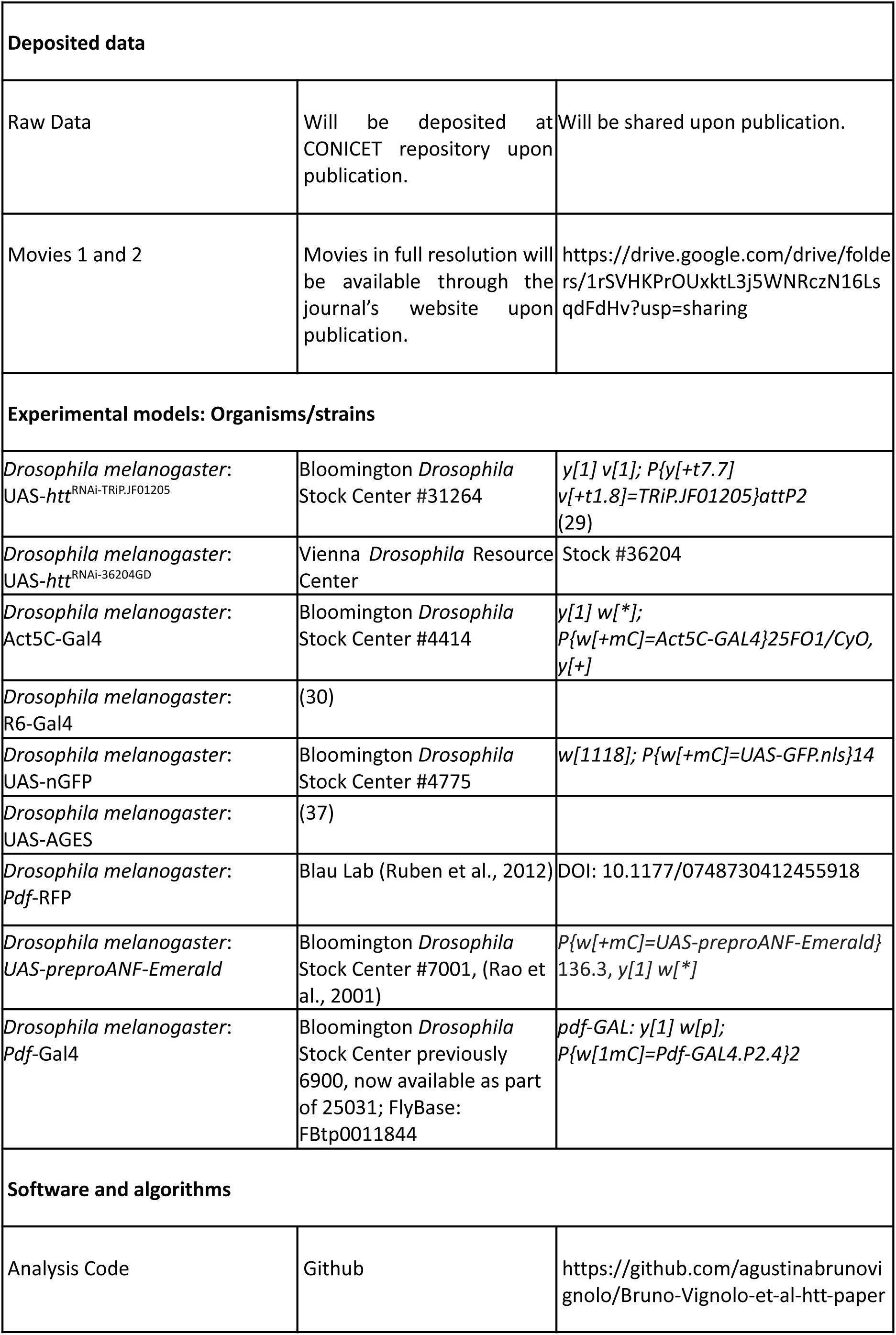
Resource Table.

### Fly Strains

All fly strains used in this study are detailed in the Key Resource Table. Flies were reared and maintained at 25°C on standard cornmeal medium under 12h:12h Light:Dark cycles, unless stated otherwise.

### Adult-specific gene downregulation

Adult-specific downregulation of *htt* was achieved using the Gal4/UAS system combined with an auxin-inducible, degradable Gal80 repressor AID-Gal80 (37). All flies were maintained on standard cornmeal food until 3 days post-eclosion. At this time, all groups except the no-auxin control were transferred to cornmeal food supplemented with auxin (1-naphthaleneacetic acid potassium salt, PhytoTech Labs, N610) which induces degradation of AID-Gal80 and thereby allows Gal4-driven expression of the RNAi construct in the experimental group. First, a 0,25M auxin aqueous solution was freshly prepared and then 2ml of this solution was added for 50 ml of cornmeal food, previously melted in the microwave and cooled down up to 70° C, to reach a final auxin concentration of 10mM. Flies were maintained on auxin-containing food (when applicable) for 5 days prior to behavioural testing and remained on the same diet throughout the experiment.

### Reverse transcription quantitative polymerase chain reaction (RT-qPCR)

Total RNA was obtained from *D. melanogaster* adults of the following genotypes: Act5C-Gal4>*w*^1118^, *w*^1118^>UAS-*htt*^RNAi-TRiP.JF01205^, Act5C-Gal4>UAS-*htt*^RNAi-TRiP.JF01205^, *w*^1118^>UAS-*htt*^RNAi-36204GD^, and Act5C-Gal4>UAS-*htt*^RNAi-36204GD^, by combining chloroform extraction and purification with silica columns (RNeasy Mini kit, Qiagen) according to the manufacturer’s instructions. All genotypes were processed using the same RNA extraction and cDNA synthesis protocol. cDNA was synthesized from the total RNA using SuperScript III Reverse Transcriptase (Invitrogen) with oligo-dT primers. Quantitative PCR reactions were performed on a 96-well format Bio-Rad PCR machine using FastStart Universal SYBR Green Mix (Roche) with primers dm_htt_fw (5’ AGCTCCGATTGTGCCGTCAAGC 3’) and dm_htt_rv (5’ CTTCTTGCCGGCCAAGGACTCG 3’) for *htt*, and dm_rp49_fw (5’ CACTTCATCCGCCACCAGTCGG 3’) and dm_rp49_rv (5’ CTCTTGAGAACGCAGGCGACCG 3’) for *DmRp49* as a reference gene. Relative RNA levels were calculated from quantification cycles (Cq) using the ΔΔCq method, with *htt* levels normalized to *DmRp49*, and all values expressed relative to Act5C-Gal4>*w*^1118^, set as 1. This driver-only control was used solely to establish the normalization baseline; statistical comparisons of knockdown efficiency for each RNAi line were instead performed against its respective genotype-matched heterozygous UAS-*htt*^RNAi^ control (driver absent).

### Free-Running Locomotor Behaviour Analysis

Flies were entrained to a 12:12h Light:Dark (LD) cycle throughout development. Newly eclosed male adults were placed in 65×5 mm glass tubes and their locomotor activity was monitored using the *Drosophila* Activity Monitor (DAM) system (TriKinetics Inc., USA). Locomotor activity was recorded in 1-minute bins, in LD conditions for 3 days, followed by at least 9 days of constant darkness (DD) to assess free-running rhythms. Data processing and analysis were performed in R using the *Rethomics* framework (38). Raw monitor files were imported and curated using the *damr* package. For circadian parameters, period length and rhythmic power were estimated via Chi-square (X^2^) periodogram analysis using the *zeitgebr* package. Individual actograms and population periodograms were generated using *ggetho*. Flies were scored as rhythmic if they displayed a single periodogram peak exceeding the α=0.05 significance threshold.

### Sleep Behaviour Analysis

Female and male flies were socially housed in vials from eclosion at 25°C under a 12:12h LD cycle. At 4–6 days of age, individual flies were transferred to 65×5 mm glass tubes containing standard cornmeal food and loaded into *Drosophila* Activity Monitors (DAM; TriKinetics Inc., USA) where activity was recorded in 1-minute bins; or into the ethoscopes arenas where activity was tracked at 2 frames per second. To account for recovery from CO_2_ anesthesia and environmental acclimation, the first 24 hours of recording were excluded, and sleep parameters were calculated starting on the second day of monitoring. Sleep was operationally defined as periods of continuous inactivity lasting at least 5 minutes (32,33). Additionally, a more stringent 25-minute inactivity threshold was applied as a robustness control for specific experiments (39). Data processing was performed using the rethomics framework (38) in R (92) employing Rstudio (93). The *sleepr* package was used to infer sleep from raw locomotor data and to extract behavioural metrics, including: Total sleep duration (24-h, daytime, and nighttime); Sleep architecture: mean sleep bout duration and sleep bout number; Sleep latency: time to the first sleep bout after “lights-on” (ZT 0) and “lights-off” (ZT 12). The control quantifications for levels of activity are critical to distinguish sleep phenotypes from general locomotor deficits. In DAMs experiments the Activity Index, the average activity counts per waking minute, was used; while in ethoscopes experiments the relative maximal velocity, mean of the maximal velocity recorded in 10 second epochs while the animal is scored as non-sleeping, was implemented. Sleep ethograms (30-minute bins) and quantification plots were generated using the *ggetho* package. All behavioural experiments were conducted in at least two to three independent biological replicates, with 15–32 individuals per genotype per trial, resulting in the total sample sizes reported in the figure legends.

### Immunostaining and Confocal Microscopy

Heads of female flies were cut at ZT2, fixed in paraformaldehyde 4% in 0.2M PB for 30 min at room temperature (RT) and brains were dissected afterward, washed four times in PBS-TritonX-100 0.2%, blocked with 7% normal goat serum for 1 h at RT and incubated with primary antibody anti-PDF (mouse monoclonal, DSHB, Cat# PDF C7, RRID: AB_760350; 1:250), ON at 4°C. After five 10-minute washes in PBS-Triton X-100 0.2%, brains were incubated with the secondary antibody (Goat anti-Mouse IgG, Invitrogen, Cat# A-11032, RRID: AB_2534091; 1:1000) for 2hs. Finally, after five washes in PBS containing 0.2% Triton X-100 (with the final wash in PBS without Triton), brains were mounted on glass slides and gently covered with Vectashield (Vector Laboratories Inc.). A square of nail polish was used as a spacer between the slide and coverslip to prevent compression of the tissue. Confocal images were obtained with air 20x Plan-Apo 20x/0.8 plus digital 4x in a Carl Zeiss LSM 710 Confocal Microscope. All the photographs within the same experiment were taken with the same confocal parameters, and only one brain hemisphere of each brain was used.

### PDF quantitation

For the quantification of PDF intensity at the sLNvs projections, we assembled a maximum intensity z-stack that contains the whole projection (approximate 10-12 images) and constructed a threshold image to create a ROI for measuring immunoreactivity intensity using ImageJ (94). Background intensity was subtracted for each brain and average intensity in arbitrary units was calculated.

### Electrophysiology

Whole-cell patch-clamp recordings were performed as described previously (21,58,95,96). Briefly, experiments in 3-to-9 day old female flies, previously anesthetized with a short incubation on ice. Brain dissection was performed in external recording solution (97), which consisted of the following (in mM): 101 NaCl, 3 KCl, 1 CaCl2, 4 MgCl2, 1.25 NaH2PO4, 5 glucose, and 20.7 NaHCO3, pH 7.2, with an osmolarity of 250 mmol/kg. LNvs were visualized by red fluorescence in *Pdf*-RFP flies, which express a red fluorophore under the Pdf promoter (57), using an Olympus BX51WI upright microscope with 60X water-immersion objective, and ThorLabs LEDD1B and TK-LED (TOLKET S.R.L.) illumination systems. Once the fluorescent cells were identified, cells were visualized under IR-DIC using a DMK23UP1300 Imaging Source camera and IC Capture 2.4 software. lLNvs were distinguished from sLNvs by their size and anatomic position. To allow access of the recording electrode, the superficial glia directly adjacent to LNvs somas was locally digested with protease XIV (10 mg/ml; P5147, Sigma-Aldrich) dissolved in external recording solution and applied with a large open tip (∼20 μm) glass capillary (pulled from borosilicate glass capillaries FG-GBF150-110-7.5 Sutter Instrument with a horizontal puller P-97 Sutter Instrument) (98). Immediately after, the protease solution was quickly washed by perfusion of the external solution using a peristaltic pump (catalog #ISM831, ISMATEC). Recordings were performed using thick-walled borosilicate glass pipettes (FG-GBF150-86-7.5, Sutter Instrument) pulled to 7-8 MΩ using a horizontal puller P-97 (Sutter Instrument) and fire polished to 9-12 MΩ. Recording pipettes were filled with internal solution containing the following (in mM): 102 potassium gluconate, 17 NaCl, 0.085 CaCl2, 0.94 EGTA, and 8.5 HEPES, pH 7.2, with an osmolarity of 235 mmol/kg. Gigaohm seals were accomplished using minimal suction followed by break-in into the whole-cell configuration using gentle suction in voltage-clamp mode with a holding voltage of –60 mV (55). Spontaneous firing was recorded in current clamp (*I=0*) mode. Recordings were made using a Multiclamp 700B amplifier controlled by pClamp 10.4 software via an Axon Digidata 1515 analog-to-digital converter (Molecular Devices), lowpass filtered at 10 kHz, and saved into.abf files. Recordings were performed during the light phase of the day, between ZT4 and ZT12. Analysis of traces was conducted using Clampfit 10.4 software. Spontaneous activity of all cells was analyzed within a restricted time post-dissection (26 minutes), as this parameter is known to influence LNv physiology (58).

### Dense-Core Vesicle Axonal Transport Quantification

Sixty-second movies of ANF::Emerald dense-core vesicles in sLNvs axonal fibres were recorded using an inverted Zeiss AxioObserver Z1 with live imaging system connected to an Axiocam HRm-3, with a temporal resolution of 0.2s, and a spatial resolution of 0.26µm. Flies were anesthetised with a brief incubation of the vial on ice and brain dissections were performed in external recording solution (97). The dissected brain was placed in a glass bottom 35mm petri dish, with the ventral surface facing downward. To determine the field of view and z-plane for each recording, sLNvs axons were identified, and then the longest region where the fibres were in focus, prior to the terminals, was selected. In most cases, movies were obtained from both hemispheres (if sLNvs axonal fibres were visible), and 3 to 4 recordings were taken per side. Recording sessions were strictly scheduled within two temporal windows: the light phase (day), spanning from ZT1 to ZT6, and the dark phase (night), from ZT13 to ZT18. To ensure physiological stability and consistency across samples, all movie acquisitions were performed between 9 and 25 minutes post-dissection. Dissected brains were observed under a 63X/NA1.4 objective and maintained at optimal conditions for live imaging. To analyze vesicle trafficking, kymographs were generated from the recordings with ImageJ using the multiple kymograph plug-in, and movement dynamics of individual vesicles was analyzed using Kymoprocessor, a semiautomated tracking toolbox developed by our group (48).

### Statistical Analysis of Dense-Core Vesicle Axonal Transport

To analyze Dense-Core Vesicle Axonal Transport, we employed Generalized Linear Mixed Models (GLMMs) using the *glmmTMB* package in R. The experimental design included a hierarchical nested structure: multiple movies (3–4 per hemisphere) were recorded from both the left and right sLNvs axonal fibers of each individual fly, with a 1-minute interval between consecutive acquisitions. To account for this dependency in the data and the potential correlation between observations from the same animal or hemisphere, we included fly, hemisphere, and experimental replicate as random effects in the model. Genotype was treated as a fixed effect. Diagnostic plots of the residuals were generated using the DHARMa package to assess model fit, including tests for overdispersion and outliers. Due to differences in variance observed between groups, a dispersion formula (*dispformula = ∼ Genotype*) was incorporated into the model to allow the residual variance to vary by genotype, ensuring the robustness of the standard error estimates. Statistical significance of the fixed effects was determined using Type II Wald chi-square tests (Analysis of Deviance) via the car package. All analyses were performed in R.

### Statistical Analysis of Sleep Behaviour in DAMs

Behavioural parameters were analyzed using generalized linear mixed models via the *glmmTMB* package in R. For each dependent variable, the most appropriate distribution family—including Gaussian (on log-transformed data), Lognormal, Beta, Negative Binomial (nbinom1 or nbinom2), Student’s t, or Gamma—was selected based on the nature of the data and model goodness-of-fit. Genotype was treated as a fixed effect, and replicate was included as a random effect. To account for observed heteroscedasticity between groups, a genotype-specific variance (dispersion) structure was incorporated using the *dispformula* argument. Model assumptions, including homoscedasticity and residual distribution, were validated by inspecting simulated residuals using the DHARMa package. Scaled residuals were compared against the expected distribution using Kolmogorov-Smirnov tests and Levene’s test for homogeneity of variance. Multiple comparisons between genotypes were performed using Sidak or Tukey post-hoc tests. Estimated marginal means (EMMs) and 95% confidence intervals were back-transformed to the original response scale for reporting. This approach ensures that the reported effect sizes and their uncertainty are biologically interpretable while maintaining the statistical rigor of the underlying non-Gaussian models. To facilitate the visualisation of significant differences, a compact letter display (CLD) was employed in all figures and tables. Genotypes sharing at least one common letter are not statistically different (*p>0.05*), whereas different letters indicate significant differences (*p<0.05*). Where applicable, data are presented as boxplots, where the horizontal line represents the median, the box indicates the interquartile range (IQR, 25th to 75th percentiles), and the whiskers extend to the most extreme data points within 1.5 times the IQR. Individual data points are overlaid as a jittered scatter to show the underlying distribution and sample size.

### Other Statistical Analyses

Statistical analyses were performed using R. For circadian assays, data were analyzed using a one-way Analysis of Variance (ANOVA) followed by Tukey’s post-hoc test for multiple comparisons. For sleep parameters recorded via the ethoscope platform, non-parametric methods were applied due to data distribution and to facilitate direct comparison with previously published datasets. Differences among treatments for each variable were assessed using a Kruskal-Wallis test, followed by Dunn’s post-hoc test for multiple comparisons between treatments. To analyze gene expression data via qPCR, Generalized Linear Mixed Models (GLMMs) were employed to account for the data structure, and multiple comparisons between genotypes were performed using Sidak’s post-hoc test. Quantification of PDF immunofluorescence intensity was evaluated using a one-way ANOVA followed by Tukey’s post-hoc test for multiple comparisons. To facilitate the visualisation of significant differences across ethoscope parameters, qPCR expression levels, and immunofluorescence quantifications, a compact letter display (CLD) was employed in the corresponding figures and tables. Genotypes sharing at least one common letter are not statistically different (*p>0.05*), whereas different letters indicate statistically significant differences (*p<0.05*). For electrophysiological recordings, statistical significance between groups was determined using Welch’s two-sample t-test.

## SUPPLEMENTARY FIGURE LEGENDS

**Supplementary Figure 1.**
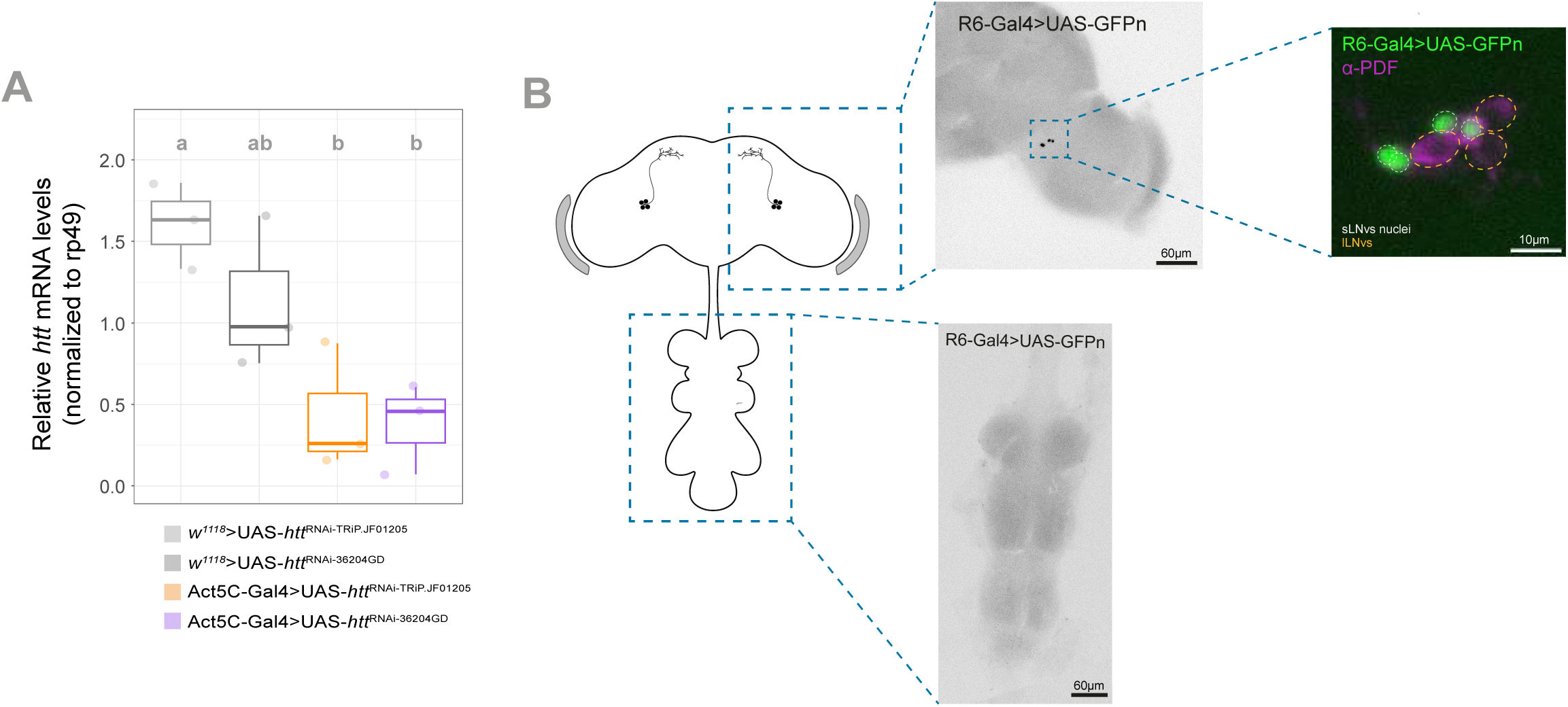
Validation of *htt* knock-down efficiency and R6-Gal4 expression pattern. **A.** Relative *htt* mRNA expression levels across genotypes, normalized to *rp49* and calculated using the 2-ΔΔCt method, with all values expressed relative to the Act5C-Gal4>*w*^1118^ driver-only control (baseline = 1.0, not shown). Grey boxes represent UAS-*htt*^RNAi^ heterozygous controls (driver absent); orange and purple boxes represent the corresponding Act5C-Gal4-driven knockdown lines. Different letters indicate statistically significant differences (*p<0.05*) among all four genotypes, determined using a GLMM followed by Sidak’s post hoc test. **B.** Anatomical specificity of the R6-Gal4 driver. Representative confocal images of an adult female brain expressing UAS-GFPn (green) under the control of R6-Gal4. Anti-PDF immunostaining (red) was used as a counterstain to confirm the identity of the sLNvs.

**Supplementary Figure 2.**
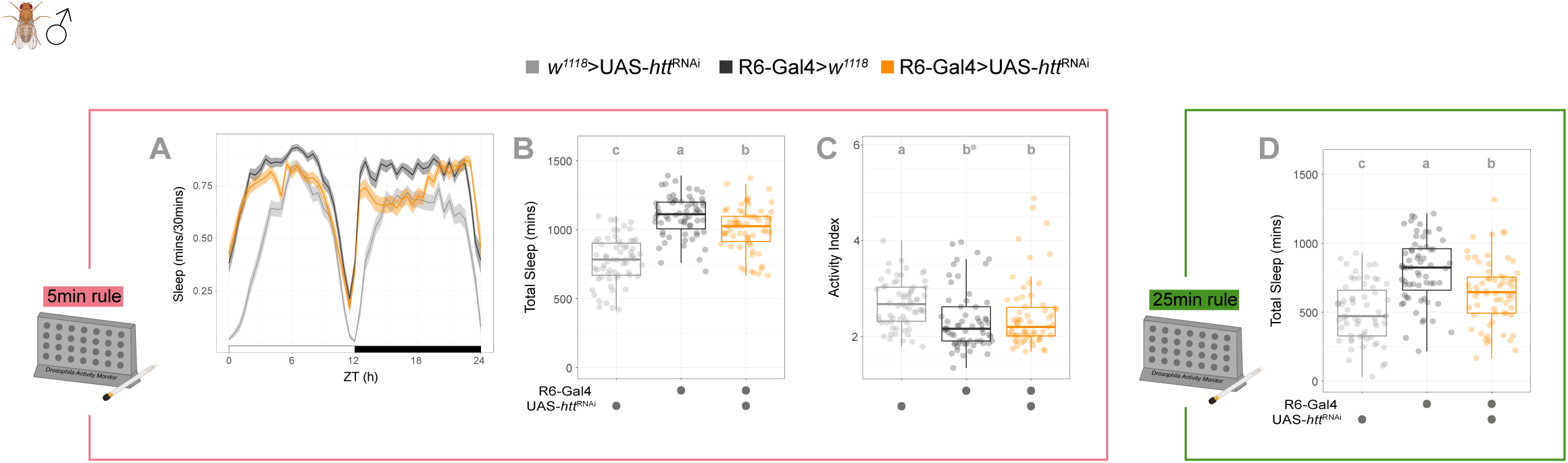
Absence of sleep alterations in male flies upon *htt* knock-down in sLNvs. 5-minute inactivity rule: **A.** Sleep ethogram. Temporal distribution of sleep in male flies over a 12:12h LD cycle. White and black bars indicate day and night, respectively. Shaded areas represent the standard deviation (SD). **B.** Total sleep. Boxplot showing the total amount of sleep (minutes) over 24 hours. **C.** Activity index. Boxplot showing the average activity per waking minute, confirming normal locomotor capacity in male flies. **25-minute inactivity rule: D.** Total 24-h sleep. Quantification of male sleep using a more stringent 25-minute inactivity threshold. Behavioural data were recorded using the Drosophila Activity Monitor system. Sample sizes: n=63-64 male flies per genotype from N=2 independent experiments. Different letters indicate statistically significant differences (*p<0.05*) using GLMMs and Sidak’s post-hoc analysis.

**Supplementary Figure 3.**
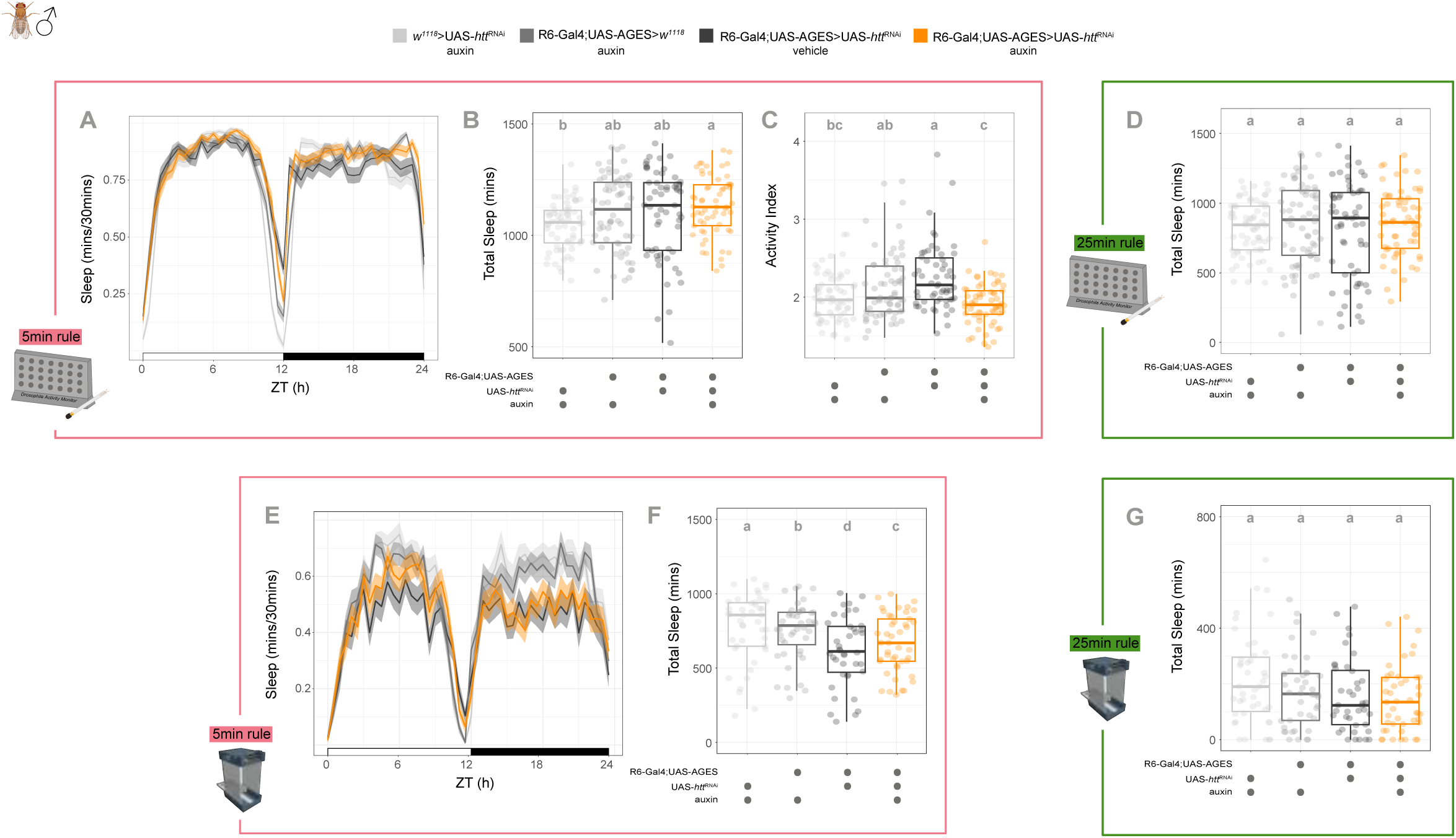
Adult-specific knock-down of *htt* in sLNvs does not affect male sleep behaviour. Adult-specific knock-down in males (DAM system): **A.** Sleep ethogram of control and experimental male flies under 12:12h LD conditions using the AGES system. **B–C.** Boxplots showing (B) total 24-h sleep (5-minute inactivity rule) and (C) activity index. **D.** Total 24-h sleep quantified using a more stringent 25-minute inactivity rule, showing no significant differences. **Adult-specific knock-down in males (Ethoscopes): E.** Sleep ethogram obtained from video-tracking analysis of adult male flies. **F.** Total 24-h sleep (5-minute inactivity rule) recorded via Ethoscopes. **G.** Total 24-h sleep (25-minute inactivity rule), confirming the lack of phenotype in males across platforms. All experiments were performed on adult male flies using the AGES system. Sleep was defined using either a 5-minute (A–C, E–F) or a 25-minute (D, G) inactivity threshold. Sample sizes for the DAM system (A–D) Sample sizes: n=58-66 male flies per genotype from N=3 independent experiments. Sample sizes for Ethoscopes (E-G): n=82–89 flies per genotype, N=3 independent experiments. Different letters indicate statistically significant differences (*p<0.05*) using GLMMs and Sidak’s post-hoc analysis.

**Supplementary Figure 4.**
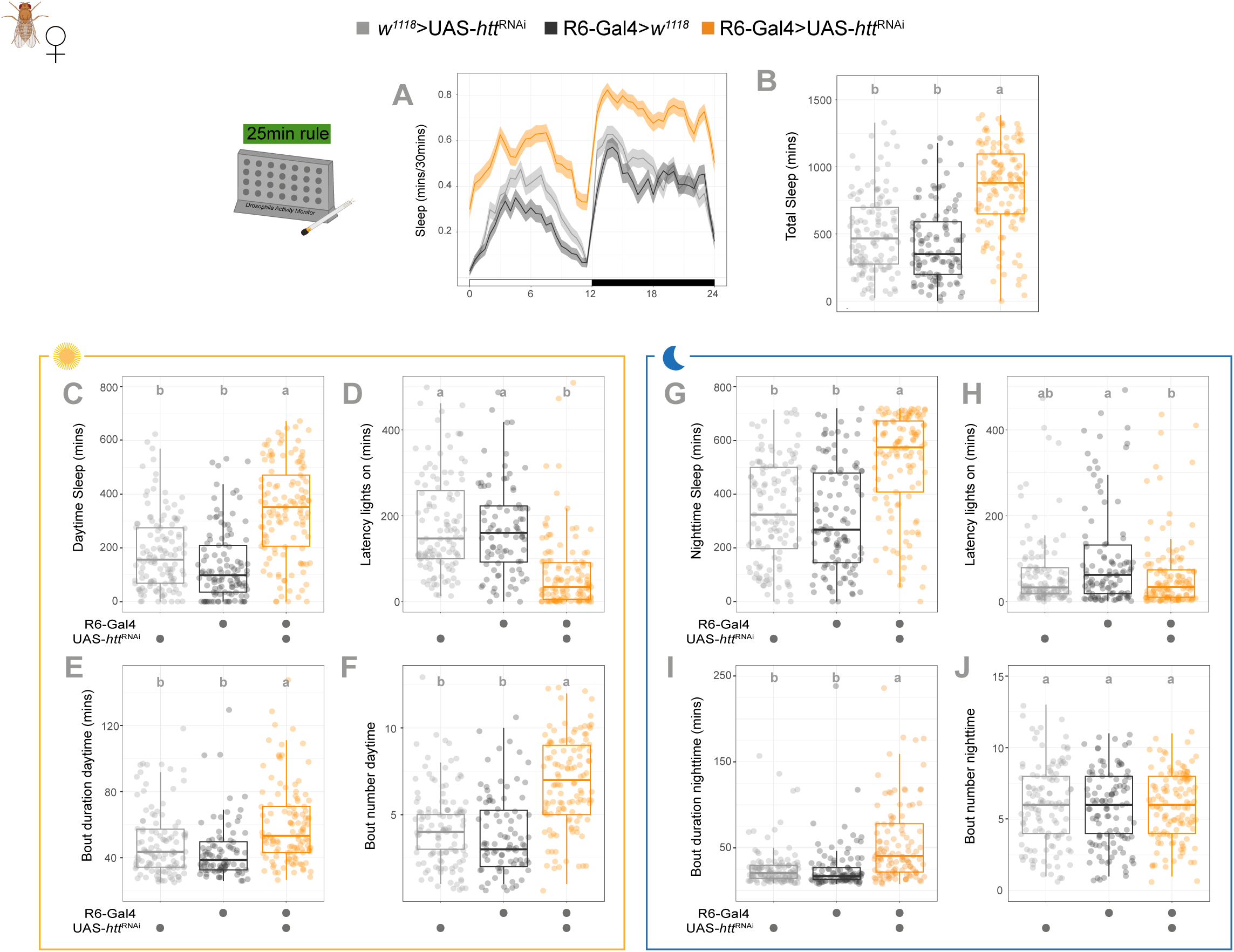
Constitutive downregulation of *htt* in sLNvs promotes sleep in female flies regardless of the sleep definition threshold. **A.** Sleep ethogram. Temporal distribution of sleep in female flies, quantified in 30-minute bins over a 24-hour LD cycle. White and black bars indicate day (ZT 0–12) and night (ZT 12–24), respectively. Shaded areas represent the standard deviation (SD). **B.** Total sleep. Boxplot showing the total amount of sleep (minutes) over the 24-hour period. **C–F.** Daytime sleep parameters. Quantification of (C) total daytime sleep duration, (D) sleep latency (time to first sleep bout after lights on), (E) mean sleep bout duration, and (F) sleep bout number during the light phase (ZT 0–12). **G–J**. Nighttime sleep parameters. Quantification of (G) total nighttime sleep duration, (H) sleep latency (time to first sleep bout after lights off), (I) mean sleep bout duration, and (J) sleep bout number during the dark phase (ZT 12–24). Data were recorded using the Drosophila Activity Monitor (DAM) system, with sleep defined as 25 minutes of continuous inactivity. Sample sizes: n=113-125 flies per genotype, N=5 independent experiments. Different letters indicate statistically significant differences (*p<0.05*) using GLMMs and Sidak’s post-hoc analysis.

**Supplementary Figure 5.**
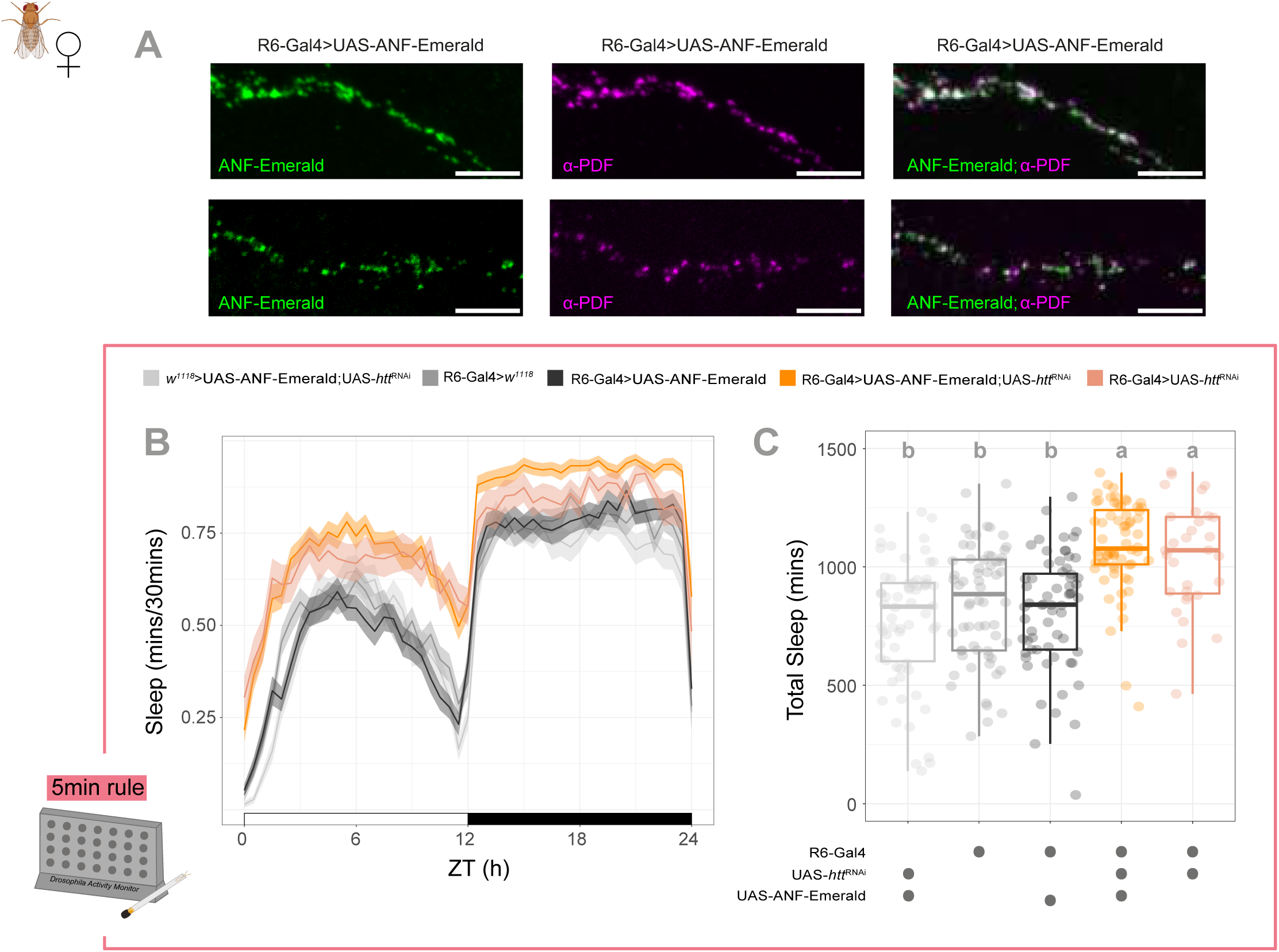
Expression of the vesicle transport reporter prepoANF-Emerald does not alter the *htt* knock-down sleep phenotype. **A.** Validation of the DCV reporter. Two representative sets of confocal images showing sLNvs axons expressing UAS-preproANF-Emerald (green, left) and immunostained for anti-PDF (magenta, middle) at ZT2. The merged image (right) shows a highly overlapping distribution of both signals, consistent with the close spatial association of both markers. Scale bars: 5 µm. **B.** Sleep ethogram. Temporal distribution of sleep in female flies over a 12:12h LD cycle. The graph compares the experimental line used for live-imaging R6-Gal4>preproANF-Emerald;UAS-*htt*^RNAi^ with the primary knockdown line R6-Gal4>UAS-*htt*^RNAi^ and their respective genetic controls. **C.** Total sleep. Boxplot showing the quantification of total 24-h sleep (minutes). Behavioural data were recorded using the *Drosophila* Activity Monitor (DAM) system with a 5-minute inactivity rule. Sample sizes: n=31-64 female flies per genotype from N=2 independent experiments. Different letters indicate statistically significant differences (*p<0.05*) using GLMMs and Sidak’s post-hoc analysis.

## SUPPLEMENTARY TABLE LEGENDS

**Supplementary Table 1.** Circadian rhythm parameters of male flies with *htt* downregulation in sLNvs. Related to Figure 1. Rhythmicity (%) indicates the percentage of rhythmic individuals out of the total flies analyzed. Values for Power-Significance and Period represent mean ± standard deviation (SD). Different letters denote statistically significant differences (*p<0.05*, One-way ANOVA followed by Tukey’s post hoc test). Sample sizes: 84-89 flies per genotype, N=3 independent experiments.

| Genotype | Rhythmicity (%) | Total flies | Rhythmic flies | Power-Significance |  |  | Period |  |  |
| --- | --- | --- | --- | --- | --- | --- | --- | --- | --- |
|  |  |  |  | Mean | SD | Significance | Mean | SD | Significance |
| *w<sup>1118</sup>*>UAS-*htt*<sup>RNAi</sup> | 86.52% | 89 | 77 | 217.66 | 159.44 | a | 23.71 | 0.27 | a |
| R6-Gal4>*w<sup>1118</sup>* | 71.43% | 84 | 60 | 150.62 | 132.87 | b | 23.61 | 0.77 | a |
| R6-Gal4>UAS-*htt*<sup>RNAi</sup> | 59.55% | 89 | 53 | **59.98** | **46.39** | **c** | 23.80 | 0.64 | a |

**Supplementary Table 2.**
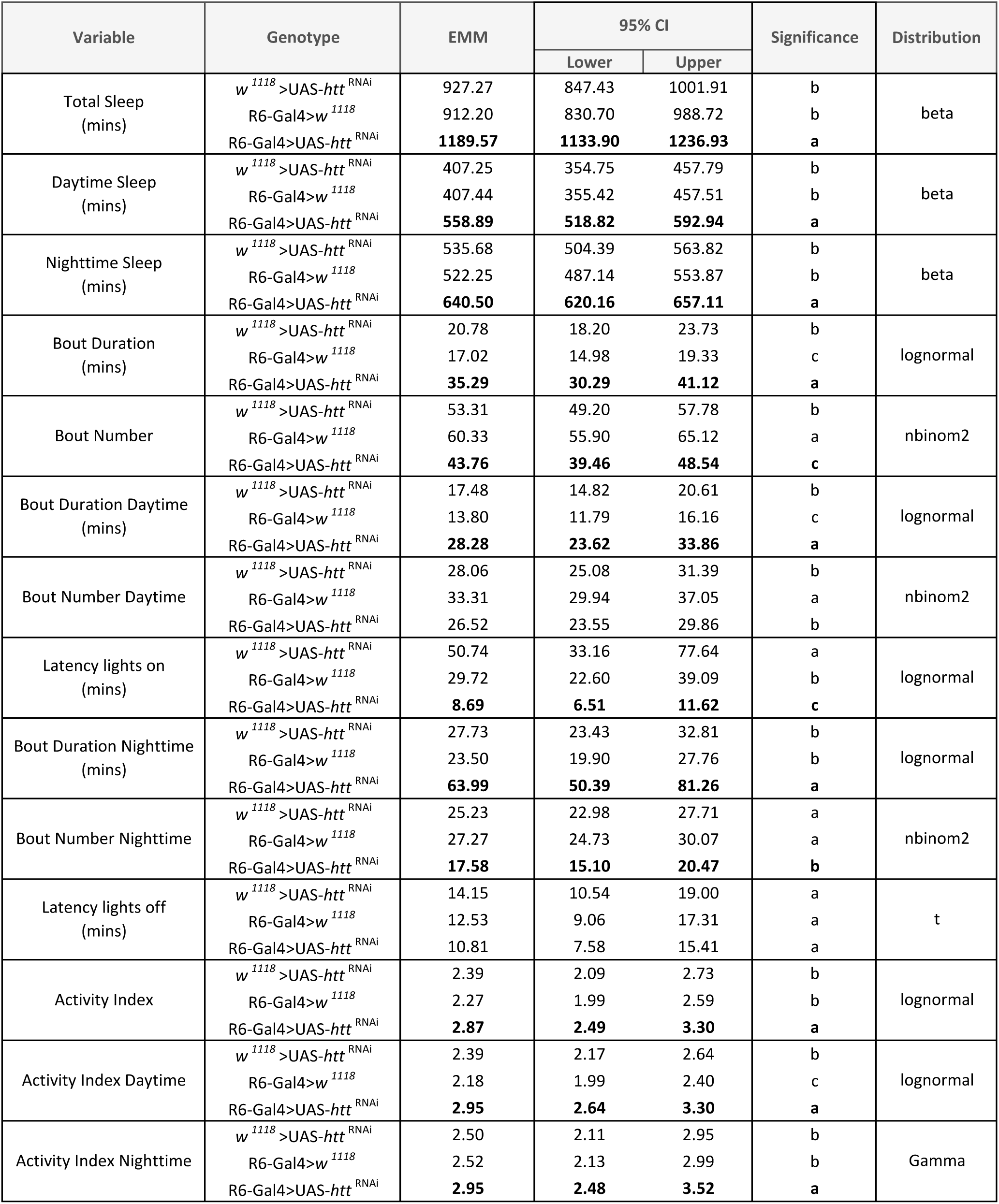
Sleep parameters in female flies with constitutive *htt* downregulation in sLNvs (DAM system, 5-min rule). Related to Figure 2. Values represent Estimated Marginal Means (EMM) and 95% Confidence Intervals (95% CI) derived from Generalized Linear Mixed Models (GLMMs). The underlying error distribution specified for each behavioural variable is indicated in the Distribution column. Different letters in the Significance column denote statistically significant differences (*p<0.05*) among genotypes within each variable, determined by post-hoc pairwise comparisons of EMMs with Sidak’s p-value adjustment. Sample sizes: 113-125 flies per genotype, N=5 independent experiments.

**Supplementary Table 3.** Sleep parameters in male flies with constitutive *htt* downregulation in sLNvs (DAM system, 5-min rule). Related to Supplementary Figure 2. Values represent Estimated Marginal Means (EMM) and 95% Confidence Intervals (95% CI) derived from Generalized Linear Mixed Models (GLMMs). The underlying error distribution specified for each behavioural variable is indicated in the Distribution column. Different letters in the Significance column denote statistically significant differences (*p<0.05*) among genotypes within each variable, determined by post-hoc pairwise comparisons of EMMs with Sidak’s p-value adjustment. Sample sizes: n=63-64 male flies per genotype, N=2 independent experiments.

| Variable | Genotype | EMM | 95% CI |  | Significance | Distribution |
| --- | --- | --- | --- | --- | --- | --- |
|  |  |  | Lower | Upper |  |  |
| Total Sleep (mins) | *w<sup>1118</sup>* >UAS-*htt*<sup>RNAi</sup> | 755.87 | 677.35 | 833.55 | c | beta |
|  | R6-Gal4>*w<sup>1118</sup>* | 1099.13 | 1034.12 | 1156.58 | a |  |
|  | R6-Gal4>UAS-*htt*<sup>RNAi</sup> | 1002.35 | 929.39 | 1069.04 | b |  |
| Daytime Sleep (mins) | *w<sup>1118</sup>* >UAS-*htt*<sup>RNAi</sup> | 347.13 | 318.20 | 376.23 | c | beta |
|  | R6-Gal4>*w<sup>1118</sup>* | 528.87 | 503.18 | 552.54 | a |  |
|  | R6-Gal4>UAS-*htt*<sup>RNAi</sup> | 483.46 | 455.30 | 510.00 | b |  |
| Nighttime Sleep (mins) | *w<sup>1118</sup>* >UAS-*htt*<sup>RNAi</sup> | 407.98 | 328.43 | 483.03 | c | beta |
|  | R6-Gal4>*w<sup>1118</sup>* | 570.05 | 510.01 | 616.41 | a |  |
|  | R6-Gal4>UAS-*htt*<sup>RNAi</sup> | 518.46 | 447.45 | 576.88 | b |  |
| Bout Duration (mins) | *w<sup>1118</sup>* >UAS-*htt*<sup>RNAi</sup> | 21.64 | 19.30 | 24.26 | b | lognormal |
|  | R6-Gal4>*w<sup>1118</sup>* | 31.25 | 27.39 | 35.66 | a |  |
|  | R6-Gal4>UAS-*htt*<sup>RNAi</sup> | 23.13 | 20.57 | 26.01 | b |  |
| Bout Number | *w<sup>1118</sup>* >UAS-*htt*<sup>RNAi</sup> | 37.11 | 34.87 | 39.50 | b | nbinom2 |
|  | R6-Gal4>*w<sup>1118</sup>* | 39.84 | 36.40 | 43.61 | b |  |
|  | R6-Gal4>UAS-*htt*<sup>RNAi</sup> | **47.08** | **44.01** | **50.36** | **a** |  |
| Bout Duration Daytime (mins) | *w<sup>1118</sup>* >UAS-*htt*<sup>RNAi</sup> | 21.32 | 18.91 | 24.04 | b | lognormal |
|  | R6-Gal4>*w<sup>1118</sup>* | 29.21 | 24.63 | 34.66 | a |  |
|  | R6-Gal4>UAS-*htt*<sup>RNAi</sup> | 19.51 | 16.96 | 22.44 | b |  |
| Bout Number Daytime | *w<sup>1118</sup>* >UAS-*htt*<sup>RNAi</sup> | 17.46 | 16.01 | 19.04 | c | nbinom2 |
|  | R6-Gal4>*w<sup>1118</sup>* | 22.54 | 20.02 | 25.37 | b |  |
|  | R6-Gal4>UAS-*htt*<sup>RNAi</sup> | **27.78** | **25.62** | **30.11** | **a** |  |
| Latency lights on (mins) | *w<sup>1118</sup>* >UAS-*htt*<sup>RNAi</sup> | 123.45 | 86.97 | 175.23 | a | lognormal |
|  | R6-Gal4>*w<sup>1118</sup>* | 40.85 | 22.30 | 74.84 | b |  |
|  | R6-Gal4>UAS-*htt*<sup>RNAi</sup> | **16.66** | **9.57** | **29.00** | **c** |  |
| Bout Duration Nighttime (mins) | *w<sup>1118</sup>* >UAS-*htt*<sup>RNAi</sup> | 24.78 | 20.04 | 30.63 | b | lognormal |
|  | R6-Gal4>*w<sup>1118</sup>* | 39.78 | 32.59 | 48.55 | a |  |
|  | R6-Gal4>UAS-*htt*<sup>RNAi</sup> | 30.65 | 25.53 | 36.81 | b |  |
| Bout Number Nighttime | *w<sup>1118</sup>* >UAS-*htt*<sup>RNAi</sup> | 19.63 | 17.63 | 21.86 | a | nbinom2 |
|  | R6-Gal4>*w<sup>1118</sup>* | 17.29 | 15.33 | 19.50 | a |  |
|  | R6-Gal4>UAS-*htt*<sup>RNAi</sup> | 19.30 | 17.38 | 21.44 | a |  |
| Latency lights off (mins) | *w<sup>1118</sup>* >UAS-*htt*<sup>RNAi</sup> | 70.15 | 57.17 | 86.08 | a | t |
|  | R6-Gal4>*w<sup>1118</sup>* | 20.51 | 16.31 | 25.80 | b |  |
|  | R6-Gal4>UAS-*htt*<sup>RNAi</sup> | 20.87 | 16.85 | 25.84 | b |  |
| Activity Index | *w<sup>1118</sup>* >UAS-*htt*<sup>RNAi</sup> | 2.68 | 2.54 | 2.83 | a | lognormal |
|  | R6-Gal4>*w<sup>1118</sup>* | 2.38 | 2.19 | 2.59 | b |  |
|  | R6-Gal4>UAS-*htt*<sup>RNAi</sup> | 2.42 | 2.25 | 2.61 | b |  |
| Activity Index Daytime | *w<sup>1118</sup>* >UAS-*htt*<sup>RNAi</sup> | 2.49 | 2.35 | 2.63 | a | lognormal |
|  | R6-Gal4>*w<sup>1118</sup>* | 2.21 | 1.99 | 2.45 | b |  |
|  | R6-Gal4>UAS-*htt*<sup>RNAi</sup> | 2.36 | 2.13 | 2.61 | ab |  |
| Activity Index Nighttime | *w<sup>1118</sup>* >UAS-*htt*<sup>RNAi</sup> | 2.92 | 2.76 | 3.09 | a | Gamma |
|  | R6-Gal4>*w<sup>1118</sup>* | 2.59 | 2.41 | 2.78 | b |  |
|  | R6-Gal4>UAS-*htt*<sup>RNAi</sup> | 2.45 | 2.31 | 2.59 | b |  |

**Supplementary Table 4.** Sleep parameters in female flies with adult-specific *htt* downregulation in sLNvs (DAM system, 5-min rule). Related to Figure 3 A-F. Values represent Estimated Marginal Means (EMM) and 95% Confidence Intervals (95% CI) derived from Generalized Linear Mixed Models (GLMMs). The underlying error distribution specified for each behavioural variable is indicated in the Distribution column. Different letters in the Significance column denote statistically significant differences (*p<0.05*) among genotypes within each variable, determined by post-hoc pairwise comparisons of EMMs with Sidak’s p-value adjustment. Sample sizes: n=59–67 flies per genotype, N=3 independent experiments.

| Variable | Genotype | Auxin | EMM | 95% CI |  | Significance | Distribution |
| --- | --- | --- | --- | --- | --- | --- | --- |
|  |  |  |  | Lower | Upper |  |  |
| Total Sleep (mins) | *w<sup>1118</sup>* >UAS-*htt*<sup>RNAi</sup> | + | 1172.53 | 1105.51 | 1228.68 | b | beta |
|  | R6-Gal4;UAS-AGES>*w<sup>1118</sup>* | + | 1143.51 | 1071.93 | 1204.24 | b |  |
|  | R6-Gal4;UAS-AGES>UAS-*htt*<sup>RNAi</sup> | - | 1116.15 | 1029.45 | 1189.01 | b |  |
|  | R6-Gal4;UAS-AGES>UAS-*htt*<sup>RNAi</sup> | + | **1265.06** | **1214.52** | **1305.52** | **a** |  |
| Daytime Sleep (mins) | *w<sup>1118</sup>* >UAS-*htt*<sup>RNAi</sup> | + | 548.26 | 508.40 | 582.65 | b | beta |
|  | R6-Gal4;UAS-AGES>*w<sup>1118</sup>* | + | 536.69 | 497.32 | 571.18 | b |  |
|  | R6-Gal4;UAS-AGES>UAS-*htt*<sup>RNAi</sup> | - | 527.38 | 481.54 | 567.21 | b |  |
|  | R6-Gal4;UAS-AGES>UAS-*htt*<sup>RNAi</sup> | + | **606.14** | **574.82** | **631.74** | **a** |  |
| Nighttime Sleep (mins) | *w<sup>1118</sup>* >UAS-*htt*<sup>RNAi</sup> | + | 628.01 | 600.41 | 655.62 | b | beta |
|  | R6-Gal4;UAS-AGES>*w<sup>1118</sup>* | + | 611.66 | 576.74 | 646.59 | b |  |
|  | R6-Gal4;UAS-AGES>UAS-*htt*<sup>RNAi</sup> | - | 590.41 | 547.87 | 632.96 | b |  |
|  | R6-Gal4;UAS-AGES>UAS-*htt*<sup>RNAi</sup> | + | **661.83** | **642.78** | **680.88** | **a** |  |
| Bout Duration (mins) | *w<sup>1118</sup>* >UAS-*htt*<sup>RNAi</sup> | + | 40.30 | 31.49 | 51.58 | ab | lognormal |
|  | R6-Gal4;UAS-AGES>*w<sup>1118</sup>* | + | 33.59 | 27.43 | 41.14 | b |  |
|  | R6-Gal4;UAS-AGES>UAS-*htt*<sup>RNAi</sup> | - | 30.23 | 23.38 | 39.09 | b |  |
|  | R6-Gal4;UAS-AGES>UAS-*htt*<sup>RNAi</sup> | + | 48.21 | 38.19 | 60.85 | a |  |
| Bout Number | *w<sup>1118</sup>* >UAS-*htt*<sup>RNAi</sup> | + | 42.93 | 35.45 | 51.99 | ab | nbinom2 |
|  | R6-Gal4;UAS-AGES>*w<sup>1118</sup>* | + | 42.81 | 36.23 | 50.58 | ab |  |
|  | R6-Gal4;UAS-AGES>UAS-*htt*<sup>RNAi</sup> | - | 51.00 | 42.35 | 61.41 | a |  |
|  | R6-Gal4;UAS-AGES>UAS-*htt*<sup>RNAi</sup> | + | 35.90 | 29.60 | 43.55 | b |  |
| Bout Duration Daytime (mins) | *w<sup>1118</sup>* >UAS-*htt*<sup>RNAi</sup> | + | 31.02 | 24.58 | 39.14 | ab | lognormal |
|  | R6-Gal4;UAS-AGES>*w<sup>1118</sup>* | + | 25.90 | 21.03 | 31.89 | a |  |
|  | R6-Gal4;UAS-AGES>UAS-*htt*<sup>RNAi</sup> | - | 25.27 | 19.32 | 33.05 | a |  |
|  | R6-Gal4;UAS-AGES>UAS-*htt*<sup>RNAi</sup> | + | 38.60 | 29.83 | 49.95 | b |  |
| Bout Number Daytime | *w<sup>1118</sup>* >UAS-*htt*<sup>RNAi</sup> | + | 24.78 | 20.72 | 29.64 | ab | nbinom2 |
|  | R6-Gal4;UAS-AGES>*w<sup>1118</sup>* | + | 27.27 | 23.19 | 32.06 | ab |  |
|  | R6-Gal4;UAS-AGES>UAS-*htt*<sup>RNAi</sup> | - | 29.53 | 24.70 | 35.31 | a |  |
|  | R6-Gal4;UAS-AGES>UAS-*htt*<sup>RNAi</sup> | + | 23.17 | 19.01 | 28.23 | b |  |
| Latency lights on (mins) | *w<sup>1118</sup>* >UAS-*htt*<sup>RNAi</sup> | + | 14.26 | 9.16 | 22.21 | a | lognormal |
|  | R6-Gal4;UAS-AGES>*w<sup>1118</sup>* | + | 16.66 | 8.42 | 32.99 | a |  |
|  | R6-Gal4;UAS-AGES>UAS-*htt*<sup>RNAi</sup> | - | 6.27 | 4.12 | 9.56 | b |  |
|  | R6-Gal4;UAS-AGES>UAS-*htt*<sup>RNAi</sup> | + | 8.38 | 5.58 | 12.58 | ab |  |
| Bout Duration Nighttime (mins) | *w<sup>1118</sup>* >UAS-*htt*<sup>RNAi</sup> | + | 75.49 | 51.86 | 109.91 | ab | lognormal |
|  | R6-Gal4;UAS-AGES>*w<sup>1118</sup>* | + | 65.38 | 46.97 | 91.00 | ab |  |
|  | R6-Gal4;UAS-AGES>UAS-*htt*<sup>RNAi</sup> | - | 50.86 | 34.77 | 74.39 | b |  |
|  | R6-Gal4;UAS-AGES>UAS-*htt*<sup>RNAi</sup> | + | 87.40 | 64.22 | 118.94 | a |  |
| Bout Number Nighttime | *w<sup>1118</sup>* >UAS-*htt*<sup>RNAi</sup> | + | 18.17 | 14.08 | 23.45 | a | nbinom2 |
|  | R6-Gal4;UAS-AGES>*w<sup>1118</sup>* | + | 16.29 | 12.86 | 20.64 | ab |  |
|  | R6-Gal4;UAS-AGES>UAS-*htt*<sup>RNAi</sup> | - | 21.81 | 17.07 | 27.86 | a |  |
|  | R6-Gal4;UAS-AGES>UAS-*htt*<sup>RNAi</sup> | + | 12.73 | 9.89 | 16.38 | b |  |
| Latency lights off (mins) | *w<sup>1118</sup>* >UAS-*htt*<sup>RNAi</sup> | + | 19.60 | 12.76 | 30.11 | a | lognormal |
|  | R6-Gal4;UAS-AGES>*w<sup>1118</sup>* | + | 32.04 | 19.56 | 52.48 | a |  |
|  | R6-Gal4;UAS-AGES>UAS-*htt*<sup>RNAi</sup> | - | 16.64 | 11.41 | 24.27 | a |  |
|  | R6-Gal4;UAS-AGES>UAS-*htt*<sup>RNAi</sup> | + | 30.90 | 17.94 | 53.22 | a |  |
| Activity Index | *w<sup>1118</sup>* >UAS-*htt*<sup>RNAi</sup> | + | 2.15 | 2.01 | 2.30 | c | lognormal |
|  | R6-Gal4;UAS-AGES>*w<sup>1118</sup>* | + | 2.37 | 2.23 | 2.53 | bc |  |
|  | R6-Gal4;UAS-AGES>UAS-*htt*<sup>RNAi</sup> | - | 2.53 | 2.27 | 2.81 | b |  |
|  | R6-Gal4;UAS-AGES>UAS-*htt*<sup>RNAi</sup> | + | **3.04** | **2.72** | **3.39** | **a** |  |
| Activity Index Daytime | *w<sup>1118</sup>* >UAS-*htt*<sup>RNAi</sup> | + | 2.01 | 1.83 | 2.21 | c | lognormal |
|  | R6-Gal4;UAS-AGES>*w<sup>1118</sup>* | + | 2.39 | 2.21 | 2.57 | b |  |
|  | R6-Gal4;UAS-AGES>UAS-*htt*<sup>RNAi</sup> | - | 2.38 | 2.12 | 2.67 | b |  |
|  | R6-Gal4;UAS-AGES>UAS-*htt*<sup>RNAi</sup> | + | **3.13** | **2.74** | **3.56** | **a** |  |
| Activity Index Nighttime | *w<sup>1118</sup>* >UAS-*htt*<sup>RNAi</sup> | + | 2.36 | 2.22 | 2.50 | c | lognormal |
|  | R6-Gal4;UAS-AGES>*w<sup>1118</sup>* | + | 2.42 | 2.26 | 2.59 | bc |  |
|  | R6-Gal4;UAS-AGES>UAS-*htt*<sup>RNAi</sup> | - | 2.69 | 2.40 | 3.01 | ab |  |
|  | R6-Gal4;UAS-AGES>UAS-*htt*<sup>RNAi</sup> | + | 2.79 | 2.53 | 3.07 | a |  |

**Supplementary Table 5.** Sleep parameters in female flies with adult-specific *htt* downregulation in sLNvs (Ethoscope system, 5-min rule). Related to Figure 3 G-I. Values represent mean ± standard deviation (SD) for each variable. The presence (+) or absence (-) of Auxin to induce adult-specific transgene expression via the AGES system is indicated. Statistical analysis was performed using non-parametric methods; different letters in the Significance column denote statistically significant differences (*p<0.05*) among genotypes within each variable, determined by Kruskal-Wallis test followed by Dunn’s post-hoc test for multiple comparisons. Sample sizes: n=82-88 flies per genotype, N=3 independent experiments.

| Variable | Genotype | Auxin | Mean | SD | Significance |
| --- | --- | --- | --- | --- | --- |
| Total Sleep<br>(mins) | *w<sup>1118</sup>*>UAS-*htt*<sup>RNAi</sup> | + | 306.48 | 203.77 | a |
|  | R6-Gal4;UAS-AGES>*w<sup>1118</sup>* | + | 203.77 | 145.61 | b |
|  | R6-Gal4;UAS-AGES>UAS-*htt*<sup>RNAi</sup> | - | 75.92 | 120.80 | c |
|  | R6-Gal4;UAS-AGES>UAS-*htt*<sup>RNAi</sup> | + | 361.30 | 203.05 | a |
| Daytime Sleep<br>(mins) | *w<sup>1118</sup>*>UAS-*htt*<sup>RNAi</sup> | + | 114.59 | 92.68 | a |
|  | R6-Gal4;UAS-AGES>*w<sup>1118</sup>* | + | 57.78 | 58.86 | b |
|  | R6-Gal4;UAS-AGES>UAS-*htt*<sup>RNAi</sup> | - | 27.28 | 52.70 | c |
|  | R6-Gal4;UAS-AGES>UAS-*htt*<sup>RNAi</sup> | + | 144.93 | 106.79 | a |
| Nighttime Sleep<br>(mins) | *w<sup>1118</sup>*>UAS-*htt*<sup>RNAi</sup> | + | 192.02 | 136.97 | b |
|  | R6-Gal4;UAS-AGES>*w<sup>1118</sup>* | + | 146.05 | 98.84 | c |
|  | R6-Gal4;UAS-AGES>UAS-*htt*<sup>RNAi</sup> | - | 48.75 | 78.21 | d |
|  | R6-Gal4;UAS-AGES>UAS-*htt*<sup>RNAi</sup> | + | **216.89** | **121.66** | **a** |
| Bout Duration<br>(mins) | *w<sup>1118</sup>*>UAS-*htt*<sup>RNAi</sup> | + | 10.31 | 3.58 | b |
|  | R6-Gal4;UAS-AGES>*w<sup>1118</sup>* | + | 9.89 | 2.27 | b |
|  | R6-Gal4;UAS-AGES>UAS-*htt*<sup>RNAi</sup> | - | 9.13 | 2.89 | b |
|  | R6-Gal4;UAS-AGES>UAS-*htt*<sup>RNAi</sup> | + | **11.66** | **2.86** | **a** |
| Bout Number | *w<sup>1118</sup>*>UAS-*htt*<sup>RNAi</sup> | + | 28.16 | 13.09 | a |
|  | R6-Gal4;UAS-AGES>*w<sup>1118</sup>* | + | 19.93 | 12.50 | b |
|  | R6-Gal4;UAS-AGES>UAS-*htt*<sup>RNAi</sup> | - | 9.58 | 10.32 | c |
|  | R6-Gal4;UAS-AGES>UAS-*htt*<sup>RNAi</sup> | + | 31.14 | 14.84 | a |
| Bout Duration Daytime<br>(mins) | *w<sup>1118</sup>*>UAS-*htt*<sup>RNAi</sup> | + | 9.45 | 3.11 | a |
|  | R6-Gal4;UAS-AGES>*w<sup>1118</sup>* | + | 7.66 | 2.65 | b |
|  | R6-Gal4;UAS-AGES>UAS-*htt*<sup>RNAi</sup> | - | 7.74 | 3.16 | b |
|  | R6-Gal4;UAS-AGES>UAS-*htt*<sup>RNAi</sup> | + | 9.49 | 2.33 | a |
| Bout Number Daytime | *w<sup>1118</sup>*>UAS-*htt*<sup>RNAi</sup> | + | 12.02 | 7.65 | a |
|  | R6-Gal4;UAS-AGES>*w<sup>1118</sup>* | + | 7.61 | 6.12 | b |
|  | R6-Gal4;UAS-AGES>UAS-*htt*<sup>RNAi</sup> | - | 4.78 | 5.40 | b |
|  | R6-Gal4;UAS-AGES>UAS-*htt*<sup>RNAi</sup> | + | 15.60 | 8.62 | a |
| Latency lights on<br>(mins) | *w<sup>1118</sup>*>UAS-*htt*<sup>RNAi</sup> | + | 2.42 | 2.51 | b |
|  | R6-Gal4;UAS-AGES>*w<sup>1118</sup>* | + | 2.76 | 2.52 | b |
|  | R6-Gal4;UAS-AGES>UAS-*htt*<sup>RNAi</sup> | - | 3.60 | 2.93 | a |
|  | R6-Gal4;UAS-AGES>UAS-*htt*<sup>RNAi</sup> | + | **1.47** | **1.77** | **c** |
| Bout Duration Nighttime<br>(mins) | *w<sup>1118</sup>*>UAS-*htt*<sup>RNAi</sup> | + | 11.36 | 9.77 | b |
|  | R6-Gal4;UAS-AGES>*w<sup>1118</sup>* | + | 10.93 | 3.38 | b |
|  | R6-Gal4;UAS-AGES>UAS-*htt*<sup>RNAi</sup> | - | 9.77 | 3.57 | b |
|  | R6-Gal4;UAS-AGES>UAS-*htt*<sup>RNAi</sup> | + | **13.01** | **4.33** | **a** |
| Bout Number Nighttime | *w<sup>1118</sup>*>UAS-*htt*<sup>RNAi</sup> | + | 16.88 | 8.58 | a |
|  | R6-Gal4;UAS-AGES>*w<sup>1118</sup>* | + | 12.95 | 7.52 | b |
|  | R6-Gal4;UAS-AGES>UAS-*htt*<sup>RNAi</sup> | - | 7.35 | 5.91 | c |
|  | R6-Gal4;UAS-AGES>UAS-*htt*<sup>RNAi</sup> | + | 16.79 | 7.38 | a |
| Latency lights off<br>(mins) | *w<sup>1118</sup>*>UAS-*htt*<sup>RNAi</sup> | + | 1.38 | 1.55 | a |
|  | R6-Gal4;UAS-AGES>*w<sup>1118</sup>* | + | 1.68 | 2.24 | a |
|  | R6-Gal4;UAS-AGES>UAS-*htt*<sup>RNAi</sup> | - | 2.45 | 2.74 | a |
|  | R6-Gal4;UAS-AGES>UAS-*htt*<sup>RNAi</sup> | + | 1.19 | 0.98 | a |
| Max Velocity | *w<sup>1118</sup>*>UAS-*htt*<sup>RNAi</sup> | + | 5.25 | 1.70 | a |
|  | R6-Gal4;UAS-AGES>*w<sup>1118</sup>* | + | 4.85 | 1.62 | a |
|  | R6-Gal4;UAS-AGES>UAS-*htt*<sup>RNAi</sup> | - | 3.29 | 0.48 | c |
|  | R6-Gal4;UAS-AGES>UAS-*htt*<sup>RNAi</sup> | + | 3.97 | 0.88 | b |
| Max Velocity Daytime | *w<sup>1118</sup>*>UAS-*htt*<sup>RNAi</sup> | + | 6.08 | 2.05 | a |
|  | R6-Gal4;UAS-AGES>*w<sup>1118</sup>* | + | 5.56 | 1.79 | a |
|  | R6-Gal4;UAS-AGES>UAS-*htt*<sup>RNAi</sup> | - | 3.42 | 0.52 | c |
|  | R6-Gal4;UAS-AGES>UAS-*htt*<sup>RNAi</sup> | + | 4.43 | 1.02 | b |
| Max Velocity Nighttime | *w<sup>1118</sup>*>UAS-*htt*<sup>RNAi</sup> | + | 4.07 | 1.27 | a |
|  | R6-Gal4;UAS-AGES>*w<sup>1118</sup>* | + | 4.03 | 2.03 | b |
|  | R6-Gal4;UAS-AGES>UAS-*htt*<sup>RNAi</sup> | - | 3.15 | 0.67 | d |
|  | R6-Gal4;UAS-AGES>UAS-*htt*<sup>RNAi</sup> | + | 3.45 | 0.86 | c |

**Supplementary Table 6.** Sleep parameters in female flies with adult-specific *htt* downregulation in sLNvs (DAM system, 25-min rule). Related to Figure 3J. Values represent Estimated Marginal Means (EMM) and 95% Confidence Intervals (95% CI) derived from Generalized Linear Mixed Models (GLMMs). The underlying error distribution specified for each behavioural variable is indicated in the Distribution column. Different letters in the Significance column denote statistically significant differences (*p<0.05*) among genotypes within each variable, determined by post-hoc pairwise comparisons of EMMs with Sidak’s p-value adjustment. Sample sizes: n=59–67 flies per genotype, N=3 independent experiments.

| Variable | Genotype | Auxin | EMM | 95% CI |  | Significance | Distribution |
| --- | --- | --- | --- | --- | --- | --- | --- |
|  |  |  |  | Lower | Upper |  |  |
| Total Sleep<br>(mins) | $w^{1118}>UAS-htt^{RNAi}$ | + | 820.55 | 684.84 | 956.27 | b | gaussian |
| | R6-Gal4;UAS-AGES>$w^{1118}$ | + | 795.18 | 672.82 | 917.54 | b | |
| | R6-Gal4;UAS-AGES>UAS-$htt^{RNAi}$ | - | 689.99 | 545.24 | 834.74 | b | |
| | R6-Gal4;UAS-AGES>UAS-$htt^{RNAi}$ | + | **993.57** | **875.34** | **1111.79** | **a** | |
| Daytime Sleep<br>(mins) | $w^{1118}>UAS-htt^{RNAi}$ | + | 336.86 | 263.11 | 410.61 | ab | gaussian |
| | R6-Gal4;UAS-AGES>$w^{1118}$ | + | 304.01 | 230.56 | 377.47 | b | |
| | R6-Gal4;UAS-AGES>UAS-$htt^{RNAi}$ | - | 271.69 | 195.44 | 347.95 | b | |
| | R6-Gal4;UAS-AGES>UAS-$htt^{RNAi}$ | + | 419.12 | 342.92 | 495.32 | a | |
| Nighttime Sleep<br>(mins) | $w^{1118}>UAS-htt^{RNAi}$ | + | 492.45 | 425.03 | 559.88 | b | beta |
| | R6-Gal4;UAS-AGES>$w^{1118}$ | + | 511.74 | 446.79 | 576.68 | b | |
| | R6-Gal4;UAS-AGES>UAS-$htt^{RNAi}$ | - | 428.84 | 348.74 | 508.95 | b | |
| | R6-Gal4;UAS-AGES>UAS-$htt^{RNAi}$ | + | **585.85** | **539.37** | **632.33** | **a** | |
| Bout Duration<br>(mins) | $w^{1118}>UAS-htt^{RNAi}$ | + | 80.98 | 66.49 | 98.62 | ab | lognormal |
| | R6-Gal4;UAS-AGES>$w^{1118}$ | + | 79.88 | 67.91 | 93.97 | ab | |
| | R6-Gal4;UAS-AGES>UAS-$htt^{RNAi}$ | - | 71.91 | 60.22 | 85.87 | b | |
| | R6-Gal4;UAS-AGES>UAS-$htt^{RNAi}$ | + | 91.66 | 77.92 | 107.83 | a | |
| Bout Number | $w^{1118}>UAS-htt^{RNAi}$ | + | 11.67 | 10.46 | 13.02 | ab | nbinom2 |
| | R6-Gal4;UAS-AGES>$w^{1118}$ | + | 10.97 | 9.81 | 12.26 | ab | |
| | R6-Gal4;UAS-AGES>UAS-$htt^{RNAi}$ | - | 10.17 | 9.01 | 11.47 | b | |
| | R6-Gal4;UAS-AGES>UAS-$htt^{RNAi}$ | + | 12.28 | 10.97 | 13.75 | a | |
| Bout Duration Daytime<br>(mins) | $w^{1118}>UAS-htt^{RNAi}$ | + | 70.69 | 57.49 | 86.91 | a | lognormal |
| | R6-Gal4;UAS-AGES>$w^{1118}$ | + | 60.23 | 50.73 | 71.51 | a | |
| | R6-Gal4;UAS-AGES>UAS-$htt^{RNAi}$ | - | 62.80 | 50.86 | 77.54 | a | |
| | R6-Gal4;UAS-AGES>UAS-$htt^{RNAi}$ | + | 74.61 | 61.30 | 90.80 | a | |
| Bout Number Daytime | $w^{1118}>UAS-htt^{RNAi}$ | + | 5.98 | 5.20 | 6.89 | ab | nbinom2 |
| | R6-Gal4;UAS-AGES>$w^{1118}$ | + | 5.83 | 5.14 | 6.62 | ab | |
| | R6-Gal4;UAS-AGES>UAS-$htt^{RNAi}$ | - | 5.26 | 4.40 | 6.29 | b | |
| | R6-Gal4;UAS-AGES>UAS-$htt^{RNAi}$ | + | 6.74 | 5.94 | 7.65 | a | |
| Latency lights on<br>(mins) | $w^{1118}>UAS-htt^{RNAi}$ | + | 118.58 | 84.90 | 165.62 | a | tweedie |
| | R6-Gal4;UAS-AGES>$w^{1118}$ | + | 105.51 | 75.50 | 147.45 | a | |
| | R6-Gal4;UAS-AGES>UAS-$htt^{RNAi}$ | - | 103.68 | 71.78 | 149.75 | a | |
| | R6-Gal4;UAS-AGES>UAS-$htt^{RNAi}$ | + | **52.38** | **34.85** | **78.73** | **b** | |
| Bout Duration Nighttime<br>(mins) | $w^{1118}>UAS-htt^{RNAi}$ | + | 109.02 | 83.57 | 142.22 | ab | lognormal |
| | R6-Gal4;UAS-AGES>$w^{1118}$ | + | 116.70 | 91.80 | 148.35 | ab | |
| | R6-Gal4;UAS-AGES>UAS-$htt^{RNAi}$ | - | 88.48 | 69.62 | 112.45 | b | |
| | R6-Gal4;UAS-AGES>UAS-$htt^{RNAi}$ | + | 129.55 | 104.39 | 160.77 | a | |
| Bout Number Nighttime | $w^{1118}>UAS-htt^{RNAi}$ | + | 6.05 | 5.21 | 7.03 | a | nbinom2 |
| | R6-Gal4;UAS-AGES>$w^{1118}$ | + | 5.46 | 4.67 | 6.39 | a | |
| | R6-Gal4;UAS-AGES>UAS-$htt^{RNAi}$ | - | 5.44 | 4.62 | 6.41 | a | |
| | R6-Gal4;UAS-AGES>UAS-$htt^{RNAi}$ | + | 5.65 | 4.81 | 6.63 | a | |
| Latency lights off<br>(mins) | $w^{1118}>UAS-htt^{RNAi}$ | + | 63.18 | 37.84 | 105.50 | a | lognormal |
| | R6-Gal4;UAS-AGES>$w^{1118}$ | + | 93.46 | 63.47 | 137.63 | a | |
| | R6-Gal4;UAS-AGES>UAS-$htt^{RNAi}$ | - | 92.09 | 55.20 | 153.66 | a | |
| | R6-Gal4;UAS-AGES>UAS-$htt^{RNAi}$ | + | 76.59 | 46.13 | 127.18 | a | |

**Supplementary Table 7.** Sleep parameters in female flies with adult-specific *htt* downregulation in sLNvs (Ethoscope system, 25-min rule). Related to Figure 3K. Values represent mean ± standard deviation (SD) for each variable. Hyphens (-) in the SD column indicate variables where only a single individual exhibited sleep bouts meeting the 25-minute criteria, precluding the calculation of a SD. The presence (+) or absence (-) of Auxin to induce adult-specific transgene expression via the AGES system is indicated. Statistical analysis was performed using non-parametric methods; different letters in the Significance column denote statistically significant differences (*p<0.05*) among genotypes within each variable, determined by Kruskal-Wallis test followed by Dunn’s post-hoc test for multiple comparisons. Sample sizes: n=82-88 flies per genotype, N=3 independent experiments.

| Variable | Genotype | Auxin | Mean | SD | Significance |
| --- | --- | --- | --- | --- | --- |
| Total Sleep<br>(mins) | *w<sup>1118</sup>*>UAS-*htt*<sup>RNAi</sup> | + | 59.02 | 116.93 | b |
|  | R6-Gal4;UAS-AGES>*w<sup>1118</sup>* | + | 27.85 | 45.33 | c |
|  | R6-Gal4;UAS-AGES>UAS-*htt*<sup>RNAi</sup> | - | 11.04 | 28.13 | d |
|  | R6-Gal4;UAS-AGES>UAS-*htt*<sup>RNAi</sup> | + | **77.55** | **93.32** | **a** |
| Daytime Sleep<br>(mins) | *w<sup>1118</sup>*>UAS-*htt*<sup>RNAi</sup> | + | 11.85 | 31.18 | b |
|  | R6-Gal4;UAS-AGES>*w<sup>1118</sup>* | + | 2.87 | 14.19 | c |
|  | R6-Gal4;UAS-AGES>UAS-*htt*<sup>RNAi</sup> | - | 2.27 | 14.72 | c |
|  | R6-Gal4;UAS-AGES>UAS-*htt*<sup>RNAi</sup> | + | **14.55** | **26.93** | **a** |
| Nighttime Sleep<br>(mins) | *w<sup>1118</sup>*>UAS-*htt*<sup>RNAi</sup> | + | 47.46 | 92.90 | b |
|  | R6-Gal4;UAS-AGES>*w<sup>1118</sup>* | + | 25.02 | 44.04 | c |
|  | R6-Gal4;UAS-AGES>UAS-*htt*<sup>RNAi</sup> | - | 8.81 | 24.93 | d |
|  | R6-Gal4;UAS-AGES>UAS-*htt*<sup>RNAi</sup> | + | **63.22** | **75.81** | **a** |
| Bout Duration<br>(mins) | *w<sup>1118</sup>*>UAS-*htt*<sup>RNAi</sup> | + | 34.55 | 9.54 | a |
|  | R6-Gal4;UAS-AGES>*w<sup>1118</sup>* | + | 35.59 | 8.42 | a |
|  | R6-Gal4;UAS-AGES>UAS-*htt*<sup>RNAi</sup> | - | 32.06 | 5.31 | a |
|  | R6-Gal4;UAS-AGES>UAS-*htt*<sup>RNAi</sup> | + | 32.82 | 6.19 | a |
| Bout Number | *w<sup>1118</sup>*>UAS-*htt*<sup>RNAi</sup> | + | 3.47 | 3.17 | a |
|  | R6-Gal4;UAS-AGES>*w<sup>1118</sup>* | + | 1.94 | 1.03 | a |
|  | R6-Gal4;UAS-AGES>UAS-*htt*<sup>RNAi</sup> | - | 1.75 | 1.04 | a |
|  | R6-Gal4;UAS-AGES>UAS-*htt*<sup>RNAi</sup> | + | 3.41 | 2.37 | a |
| Bout Duration Daytime<br>(mins) | *w<sup>1118</sup>*>UAS-*htt*<sup>RNAi</sup> | + | 30.07 | 5.23 | a |
|  | R6-Gal4;UAS-AGES>*w<sup>1118</sup>* | + | 41.00 | 1.41 | a |
|  | R6-Gal4;UAS-AGES>UAS-*htt*<sup>RNAi</sup> | - | 31.39 | - | a |
|  | R6-Gal4;UAS-AGES>UAS-*htt*<sup>RNAi</sup> | + | 29.26 | 11.30 | a |
| Bout Number Daytime | *w<sup>1118</sup>*>UAS-*htt*<sup>RNAi</sup> | + | 2.00 | 1.20 | a |
|  | R6-Gal4;UAS-AGES>*w<sup>1118</sup>* | + | 1.50 | 0.71 | a |
|  | R6-Gal4;UAS-AGES>UAS-*htt*<sup>RNAi</sup> | - | 3.00 | - | a |
|  | R6-Gal4;UAS-AGES>UAS-*htt*<sup>RNAi</sup> | + | 1.69 | 0.85 | a |
| Latency lights on<br>(mins) | *w<sup>1118</sup>*>UAS-*htt*<sup>RNAi</sup> | + | 4.15 | 2.73 | a |
|  | R6-Gal4;UAS-AGES>*w<sup>1118</sup>* | + | 3.36 | 0.07 | a |
|  | R6-Gal4;UAS-AGES>UAS-*htt*<sup>RNAi</sup> | - | 2.12 | - | a |
|  | R6-Gal4;UAS-AGES>UAS-*htt*<sup>RNAi</sup> | + | 4.30 | 2.99 | a |
| Bout Duration Nighttime<br>(mins) | *w<sup>1118</sup>*>UAS-*htt*<sup>RNAi</sup> | + | 37.39 | 16.99 | a |
|  | R6-Gal4;UAS-AGES>*w<sup>1118</sup>* | + | 35.01 | 8.52 | a |
|  | R6-Gal4;UAS-AGES>UAS-*htt*<sup>RNAi</sup> | - | 32.15 | 5.73 | a |
|  | R6-Gal4;UAS-AGES>UAS-*htt*<sup>RNAi</sup> | + | 34.45 | 6.99 | a |
| Bout Number Nighttime | *w<sup>1118</sup>*>UAS-*htt*<sup>RNAi</sup> | + | 2.94 | 2.16 | a |
|  | R6-Gal4;UAS-AGES>*w<sup>1118</sup>* | + | 1.88 | 1.09 | a |
|  | R6-Gal4;UAS-AGES>UAS-*htt*<sup>RNAi</sup> | - | 1.57 | 0.98 | a |
|  | R6-Gal4;UAS-AGES>UAS-*htt*<sup>RNAi</sup> | + | 2.85 | 1.75 | a |
| Latency lights off<br>(mins) | *w<sup>1118</sup>*>UAS-*htt*<sup>RNAi</sup> | + | 3.00 | 1.97 | a |
|  | R6-Gal4;UAS-AGES>*w<sup>1118</sup>* | + | 5.21 | 3.66 | a |
|  | R6-Gal4;UAS-AGES>UAS-*htt*<sup>RNAi</sup> | - | 4.19 | 3.65 | a |
|  | R6-Gal4;UAS-AGES>UAS-*htt*<sup>RNAi</sup> | + | 2.84 | 2.22 | a |

**Supplementary Table 8.** Sleep parameters in male flies with adult-specific *htt* downregulation in sLNvs (DAM system, 5-min rule). Related to Supplementary Figure 4A-C. Values represent Estimated Marginal Means (EMM) and 95% Confidence Intervals (95% CI) derived from Generalized Linear Mixed Models (GLMMs). The underlying error distribution specified for each behavioural variable is indicated in the Distribution column. Different letters in the Significance column denote statistically significant differences (*p<0.05*) among genotypes within each variable, determined by post-hoc pairwise comparisons of EMMs with Sidak’s p-value adjustment. Sample sizes: n=63-64 flies per genotype, N=2 independent experiments.

| Variable | Genotype | Auxin | EMM | 95% CI |  | Significance | Distribution |
| --- | --- | --- | --- | --- | --- | --- | --- |
|  |  |  |  | Lower | Upper |  |  |
| Total Sleep (mins) | $w^{1118}>UAS-htt^{RNAi}$ | + | 1044.35 | 1002.31 | 1083.79 | b | beta |
| | R6-Gal4;UAS-AGES>$w^{1118}$ | + | 1102.90 | 1042.36 | 1156.73 | ab | |
| | R6-Gal4;UAS-AGES>UAS-$htt^{RNAi}$ | - | 1093.18 | 1022.48 | 1155.23 | ab | |
| | R6-Gal4;UAS-AGES>UAS-$htt^{RNAi}$ | + | 1132.07 | 1082.21 | 1176.67 | a | |
| Daytime Sleep (mins) | $w^{1118}>UAS-htt^{RNAi}$ | + | 486.11 | 464.63 | 506.62 | b | beta |
| | R6-Gal4;UAS-AGES>$w^{1118}$ | + | 519.42 | 484.62 | 550.87 | ab | |
| | R6-Gal4;UAS-AGES>UAS-$htt^{RNAi}$ | - | 529.18 | 490.82 | 563.17 | a | |
| | R6-Gal4;UAS-AGES>UAS-$htt^{RNAi}$ | + | 536.67 | 510.56 | 560.55 | a | |
| Nighttime Sleep (mins) | $w^{1118}>UAS-htt^{RNAi}$ | + | 563.26 | 538.82 | 585.23 | b | beta |
| | R6-Gal4;UAS-AGES>$w^{1118}$ | + | 587.56 | 556.93 | 613.53 | ab | |
| | R6-Gal4;UAS-AGES>UAS-$htt^{RNAi}$ | - | 565.78 | 526.18 | 599.15 | ab | |
| | R6-Gal4;UAS-AGES>UAS-$htt^{RNAi}$ | + | 599.33 | 572.72 | 621.95 | a | |
| Bout Duration (mins) | $w^{1118}>UAS-htt^{RNAi}$ | + | 44.07 | 36.32 | 53.48 | a | lognormal |
| | R6-Gal4;UAS-AGES>$w^{1118}$ | + | 51.06 | 38.93 | 66.98 | a | |
| | R6-Gal4;UAS-AGES>UAS-$htt^{RNAi}$ | - | 48.73 | 35.39 | 67.09 | a | |
| | R6-Gal4;UAS-AGES>UAS-$htt^{RNAi}$ | + | 40.71 | 33.05 | 50.15 | a | |
| Bout Number | $w^{1118}>UAS-htt^{RNAi}$ | + | 29.21 | 23.39 | 36.47 | b | nbinom2 |
| | R6-Gal4;UAS-AGES>$w^{1118}$ | + | 32.58 | 25.43 | 41.74 | ab | |
| | R6-Gal4;UAS-AGES>UAS-$htt^{RNAi}$ | - | 36.85 | 28.66 | 47.38 | a | |
| | R6-Gal4;UAS-AGES>UAS-$htt^{RNAi}$ | + | 35.43 | 28.38 | 44.22 | a | |
| Bout Duration Daytime (mins) | $w^{1118}>UAS-htt^{RNAi}$ | + | 42.19 | 34.18 | 52.08 | a | lognormal |
| | R6-Gal4;UAS-AGES>$w^{1118}$ | + | 54.80 | 40.42 | 74.31 | a | |
| | R6-Gal4;UAS-AGES>UAS-$htt^{RNAi}$ | - | 49.54 | 35.14 | 69.82 | a | |
| | R6-Gal4;UAS-AGES>UAS-$htt^{RNAi}$ | + | 43.56 | 33.77 | 56.18 | a | |
| Bout Number Daytime | $w^{1118}>UAS-htt^{RNAi}$ | + | 15.04 | 11.71 | 19.32 | b | nbinom2 |
| | R6-Gal4;UAS-AGES>$w^{1118}$ | + | 15.77 | 11.95 | 20.81 | ab | |
| | R6-Gal4;UAS-AGES>UAS-$htt^{RNAi}$ | - | 19.07 | 14.39 | 25.27 | a | |
| | R6-Gal4;UAS-AGES>UAS-$htt^{RNAi}$ | + | 18.40 | 14.20 | 23.85 | ab | |
| Latency lights on (mins) | $w^{1118}>UAS-htt^{RNAi}$ | + | 55.83 | 46.86 | 64.81 | a | gaussian |
| | R6-Gal4;UAS-AGES>$w^{1118}$ | + | 36.00 | 26.35 | 45.65 | b | |
| | R6-Gal4;UAS-AGES>UAS-$htt^{RNAi}$ | - | 41.48 | 31.33 | 51.63 | b | |
| | R6-Gal4;UAS-AGES>UAS-$htt^{RNAi}$ | + | 31.52 | 24.56 | 38.49 | b | |
| Bout Duration Nighttime (mins) | $w^{1118}>UAS-htt^{RNAi}$ | + | 56.93 | 45.01 | 72.01 | a | lognormal |
| | R6-Gal4;UAS-AGES>$w^{1118}$ | + | 57.77 | 43.06 | 77.51 | a | |
| | R6-Gal4;UAS-AGES>UAS-$htt^{RNAi}$ | - | 60.88 | 41.68 | 88.92 | a | |
| | R6-Gal4;UAS-AGES>UAS-$htt^{RNAi}$ | + | 54.88 | 41.41 | 72.74 | a | |
| Bout Number Nighttime | $w^{1118}>UAS-htt^{RNAi}$ | + | 14.06 | 11.40 | 17.36 | a | nbinom1 |
| | R6-Gal4;UAS-AGES>$w^{1118}$ | + | 16.78 | 13.32 | 21.13 | a | |
| | R6-Gal4;UAS-AGES>UAS-$htt^{RNAi}$ | - | 18.00 | 14.26 | 22.71 | a | |
| | R6-Gal4;UAS-AGES>UAS-$htt^{RNAi}$ | + | 17.05 | 13.83 | 21.03 | a | |
| Latency lights off (mins) | $w^{1118}>UAS-htt^{RNAi}$ | + | 765.33 | 759.64 | 771.07 | a | t |
| | R6-Gal4;UAS-AGES>$w^{1118}$ | + | 740.63 | 732.60 | 748.74 | b | |
| | R6-Gal4;UAS-AGES>UAS-$htt^{RNAi}$ | - | 725.57 | 723.32 | 727.82 | c | |
| | R6-Gal4;UAS-AGES>UAS-$htt^{RNAi}$ | + | 731.42 | 727.42 | 735.44 | b | |
| Activity Index | $w^{1118}>UAS-htt^{RNAi}$ | + | 2.00 | 1.83 | 2.19 | bc | lognormal |
| | R6-Gal4;UAS-AGES>$w^{1118}$ | + | 2.14 | 1.94 | 2.36 | ab | |
| | R6-Gal4;UAS-AGES>UAS-$htt^{RNAi}$ | - | 2.26 | 2.04 | 2.50 | a | |
| | R6-Gal4;UAS-AGES>UAS-$htt^{RNAi}$ | + | 1.90 | 1.74 | 2.08 | c | |
| Activity Index Daytime | $w^{1118}>UAS-htt^{RNAi}$ | + | 1.75 | 1.60 | 1.92 | bc | lognormal |
| | R6-Gal4;UAS-AGES>$w^{1118}$ | + | 1.88 | 1.69 | 2.09 | ab | |
| | R6-Gal4;UAS-AGES>UAS-$htt^{RNAi}$ | - | 2.07 | 1.84 | 2.32 | a | |
| | R6-Gal4;UAS-AGES>UAS-$htt^{RNAi}$ | + | 1.72 | 1.57 | 1.89 | c | |
| Activity Index Nighttime | $w^{1118}>UAS-htt^{RNAi}$ | + | 2.30 | 2.07 | 2.56 | ab | Gamma |
| | R6-Gal4;UAS-AGES>$w^{1118}$ | + | 2.40 | 2.15 | 2.69 | a | |
| | R6-Gal4;UAS-AGES>UAS-$htt^{RNAi}$ | - | 2.39 | 2.13 | 2.67 | a | |
| | R6-Gal4;UAS-AGES>UAS-$htt^{RNAi}$ | + | 2.15 | 1.93 | 2.39 | b | |

**Supplementary Table 9.** Sleep parameters in male flies with adult-specific *htt* downregulation in sLNvs (DAM system, 25-min rule). Related to Supplementary Figure 4D. Values represent Estimated Marginal Means (EMM) and 95% Confidence Intervals (95% CI) derived from Generalized Linear Mixed Models (GLMMs). The underlying error distribution specified for each behavioural variable is indicated in the Distribution column. Different letters in the Significance column denote statistically significant differences (*p<0.05*) among genotypes within each variable, determined by post-hoc pairwise comparisons of EMMs with Sidak’s p-value adjustment. Sample sizes: n=63-64 flies per genotype, N=2 independent experiments.

| Variable and distribution | Genotype | Auxin | EMM | 95% CI |  | Significance | Distribution |
| --- | --- | --- | --- | --- | --- | --- | --- |
|  |  |  |  | Lower | Upper |  |  |
| Total Sleep (mins) | *w<sup>1118</sup>*>UAS-*htt*<sup>RNAi</sup> | + | 827.09 | 731.06 | 919.37 | a | beta |
|  | R6-Gal4;UAS-AGES>*w<sup>1118</sup>* | + | 841.18 | 721.93 | 953.96 | a |  |
|  | R6-Gal4;UAS-AGES>UAS-*htt*<sup>RNAi</sup> | - | 798.69 | 668.86 | 923.53 | a |  |
|  | R6-Gal4;UAS-AGES>UAS-*htt*<sup>RNAi</sup> | + | 857.70 | 753.23 | 956.49 | a |  |
| Daytime Sleep (mins) | *w<sup>1118</sup>*>UAS-*htt*<sup>RNAi</sup> | + | 362.24 | 291.49 | 432.81 | a | beta |
|  | R6-Gal4;UAS-AGES>*w<sup>1118</sup>* | + | 394.39 | 323.03 | 463.12 | a |  |
|  | R6-Gal4;UAS-AGES>UAS-*htt*<sup>RNAi</sup> | - | 376.78 | 304.31 | 447.92 | a |  |
|  | R6-Gal4;UAS-AGES>UAS-*htt*<sup>RNAi</sup> | + | 375.79 | 304.14 | 446.21 | a |  |
| Nighttime Sleep (mins) | *w<sup>1118</sup>*>UAS-*htt*<sup>RNAi</sup> | + | 473.55 | 422.41 | 520.06 | a | beta |
|  | R6-Gal4;UAS-AGES>*w<sup>1118</sup>* | + | 462.29 | 399.75 | 518.77 | a |  |
|  | R6-Gal4;UAS-AGES>UAS-*htt*<sup>RNAi</sup> | - | 435.50 | 366.49 | 499.16 | a |  |
|  | R6-Gal4;UAS-AGES>UAS-*htt*<sup>RNAi</sup> | + | 486.92 | 430.45 | 537.06 | a |  |
| Bout Duration (mins) | *w<sup>1118</sup>*>UAS-*htt*<sup>RNAi</sup> | + | 87.32 | 74.57 | 102.25 | b | lognormal |
|  | R6-Gal4;UAS-AGES>*w<sup>1118</sup>* | + | 115.96 | 91.61 | 146.78 | a |  |
|  | R6-Gal4;UAS-AGES>UAS-*htt*<sup>RNAi</sup> | - | 98.26 | 77.81 | 124.09 | ab |  |
|  | R6-Gal4;UAS-AGES>UAS-*htt*<sup>RNAi</sup> | + | 93.59 | 76.96 | 113.81 | ab |  |
| Bout Number | *w<sup>1118</sup>*>UAS-*htt*<sup>RNAi</sup> | + | 11.21 | 10.23 | 12.29 | ab | nbinom2 |
|  | R6-Gal4;UAS-AGES>*w<sup>1118</sup>* | + | 9.59 | 8.50 | 10.82 | b |  |
|  | R6-Gal4;UAS-AGES>UAS-*htt*<sup>RNAi</sup> | - | 10.28 | 8.96 | 11.78 | ab |  |
|  | R6-Gal4;UAS-AGES>UAS-*htt*<sup>RNAi</sup> | + | 11.34 | 10.28 | 12.52 | a |  |
| Bout Duration Daytime (mins) | *w<sup>1118</sup>*>UAS-*htt*<sup>RNAi</sup> | + | 102.98 | 85.98 | 123.34 | b | lognormal |
|  | R6-Gal4;UAS-AGES>*w<sup>1118</sup>* | + | 174.87 | 126.10 | 242.51 | a |  |
|  | R6-Gal4;UAS-AGES>UAS-*htt*<sup>RNAi</sup> | - | 120.69 | 89.82 | 162.17 | ab |  |
|  | R6-Gal4;UAS-AGES>UAS-*htt*<sup>RNAi</sup> | + | 118.17 | 91.61 | 152.43 | ab |  |
| Bout Number Daytime | *w<sup>1118</sup>*>UAS-*htt*<sup>RNAi</sup> | + | 4.77 | 4.14 | 5.51 | a | nbinom2 |
|  | R6-Gal4;UAS-AGES>*w<sup>1118</sup>* | + | 4.08 | 3.49 | 4.76 | a |  |
|  | R6-Gal4;UAS-AGES>UAS-*htt*<sup>RNAi</sup> | - | 4.82 | 4.14 | 5.62 | a |  |
|  | R6-Gal4;UAS-AGES>UAS-*htt*<sup>RNAi</sup> | + | 4.75 | 4.10 | 5.52 | a |  |
| Latency lights on (mins) | *w<sup>1118</sup>*>UAS-*htt*<sup>RNAi</sup> | + | 125.47 | 98.65 | 159.59 | a | lognormal |
|  | R6-Gal4;UAS-AGES>*w<sup>1118</sup>* | + | 113.79 | 85.39 | 151.64 | a |  |
|  | R6-Gal4;UAS-AGES>UAS-*htt*<sup>RNAi</sup> | - | 130.80 | 96.82 | 176.72 | a |  |
|  | R6-Gal4;UAS-AGES>UAS-*htt*<sup>RNAi</sup> | + | 133.73 | 102.94 | 173.72 | a |  |
| Bout Duration Nighttime (mins) | *w<sup>1118</sup>*>UAS-*htt*<sup>RNAi</sup> | + | 65.17 | 53.00 | 80.15 | a | t |
|  | R6-Gal4;UAS-AGES>*w<sup>1118</sup>* | + | 66.91 | 53.90 | 83.05 | a |  |
|  | R6-Gal4;UAS-AGES>UAS-*htt*<sup>RNAi</sup> | - | 59.73 | 48.83 | 73.07 | a |  |
|  | R6-Gal4;UAS-AGES>UAS-*htt*<sup>RNAi</sup> | + | 60.25 | 50.16 | 72.36 | a |  |
| Bout Number Nighttime | *w<sup>1118</sup>*>UAS-*htt*<sup>RNAi</sup> | + | 6.44 | 5.71 | 7.27 | a | nbinom1 |
|  | R6-Gal4;UAS-AGES>*w<sup>1118</sup>* | + | 5.58 | 4.90 | 6.35 | a |  |
|  | R6-Gal4;UAS-AGES>UAS-*htt*<sup>RNAi</sup> | - | 5.53 | 4.82 | 6.36 | a |  |
|  | R6-Gal4;UAS-AGES>UAS-*htt*<sup>RNAi</sup> | + | 6.59 | 5.82 | 7.46 | a |  |
| Latency lights off (mins) | *w<sup>1118</sup>*>UAS-*htt*<sup>RNAi</sup> | + | 771.93 | 764.55 | 779.39 | a | t |
|  | R6-Gal4;UAS-AGES>*w<sup>1118</sup>* | + | 772.78 | 757.38 | 788.50 | a |  |
|  | R6-Gal4;UAS-AGES>UAS-*htt*<sup>RNAi</sup> | - | 729.37 | 719.09 | 739.79 | c |  |
|  | R6-Gal4;UAS-AGES>UAS-*htt*<sup>RNAi</sup> | + | 746.93 | 738.13 | 755.84 | b |  |

**Supplementary Table 10.** Sleep parameters in male flies with adult-specific *htt* downregulation in sLNvs (Ethoscope system, 5-min rule). Related to Supplementary Figure 4E-F. Values represent mean ± standard deviation (SD) for each variable. The presence (+) or absence (-) of Auxin to induce adult-specific transgene expression via the AGES system is indicated. Statistical analysis was performed using non-parametric methods; different letters in the Significance column denote statistically significant differences (*p<0.05*) among genotypes within each variable, determined by Kruskal-Wallis test followed by Dunn’s post-hoc test for multiple comparisons. Sample sizes: n=82-89 flies per genotype, N=3 independent experiments.

| Variable | Genotype | Auxin | Mean | SD | Significance |
| --- | --- | --- | --- | --- | --- |
| Total Sleep<br>(mins) | *w<sup>1118</sup>*>UAS-*htt*<sup>RNAi</sup> | + | 771.31 | 231.90 | a |
|  | R6-Gal4;UAS-AGES>*w<sup>1118</sup>* | + | 748.86 | 192.10 | b |
|  | R6-Gal4;UAS-AGES>UAS-*htt*<sup>RNAi</sup> | - | 615.05 | 231.80 | d |
|  | R6-Gal4;UAS-AGES>UAS-*htt*<sup>RNAi</sup> | + | 664.26 | 195.96 | c |
| Daytime Sleep<br>(mins) | *w<sup>1118</sup>*>UAS-*htt*<sup>RNAi</sup> | + | 350.07 | 116.21 | a |
|  | R6-Gal4;UAS-AGES>*w<sup>1118</sup>* | + | 338.42 | 107.82 | a |
|  | R6-Gal4;UAS-AGES>UAS-*htt*<sup>RNAi</sup> | - | 292.39 | 120.91 | a |
|  | R6-Gal4;UAS-AGES>UAS-*htt*<sup>RNAi</sup> | + | 327.03 | 107.85 | a |
| Nighttime Sleep<br>(mins) | *w<sup>1118</sup>*>UAS-*htt*<sup>RNAi</sup> | + | 421.25 | 145.87 | a |
|  | R6-Gal4;UAS-AGES>*w<sup>1118</sup>* | + | 410.38 | 121.33 | a |
|  | R6-Gal4;UAS-AGES>UAS-*htt*<sup>RNAi</sup> | - | 322.77 | 145.52 | b |
|  | R6-Gal4;UAS-AGES>UAS-*htt*<sup>RNAi</sup> | + | 337.22 | 131.59 | b |
| Bout Duration<br>(mins) | *w<sup>1118</sup>*>UAS-*htt*<sup>RNAi</sup> | + | 13.28 | 2.98 | a |
|  | R6-Gal4;UAS-AGES>*w<sup>1118</sup>* | + | 12.46 | 2.30 | a |
|  | R6-Gal4;UAS-AGES>UAS-*htt*<sup>RNAi</sup> | - | 12.25 | 2.45 | a |
|  | R6-Gal4;UAS-AGES>UAS-*htt*<sup>RNAi</sup> | + | 12.36 | 2.55 | a |
| Bout Number | *w<sup>1118</sup>*>UAS-*htt*<sup>RNAi</sup> | + | 57.88 | 13.99 | b |
|  | R6-Gal4;UAS-AGES>*w<sup>1118</sup>* | + | 59.68 | 10.53 | a |
|  | R6-Gal4;UAS-AGES>UAS-*htt*<sup>RNAi</sup> | - | 48.88 | 14.18 | d |
|  | R6-Gal4;UAS-AGES>UAS-*htt*<sup>RNAi</sup> | + | 53.53 | 11.63 | c |
| Bout Duration Daytime<br>(mins) | *w<sup>1118</sup>*>UAS-*htt*<sup>RNAi</sup> | + | 12.13 | 3.23 | a |
|  | R6-Gal4;UAS-AGES>*w<sup>1118</sup>* | + | 11.44 | 2.39 | a |
|  | R6-Gal4;UAS-AGES>UAS-*htt*<sup>RNAi</sup> | - | 11.17 | 2.67 | a |
|  | R6-Gal4;UAS-AGES>UAS-*htt*<sup>RNAi</sup> | + | 12.37 | 3.02 | a |
| Bout Number Daytime | *w<sup>1118</sup>*>UAS-*htt*<sup>RNAi</sup> | + | 28.81 | 8.16 | a |
|  | R6-Gal4;UAS-AGES>*w<sup>1118</sup>* | + | 29.30 | 8.06 | a |
|  | R6-Gal4;UAS-AGES>UAS-*htt*<sup>RNAi</sup> | - | 25.93 | 9.57 | a |
|  | R6-Gal4;UAS-AGES>UAS-*htt*<sup>RNAi</sup> | + | 26.84 | 8.26 | a |
| Latency lights on<br>(mins) | *w<sup>1118</sup>*>UAS-*htt*<sup>RNAi</sup> | + | 1.42 | 0.98 | a |
|  | R6-Gal4;UAS-AGES>*w<sup>1118</sup>* | + | 1.22 | 0.96 | a |
|  | R6-Gal4;UAS-AGES>UAS-*htt*<sup>RNAi</sup> | - | 1.29 | 1.17 | a |
|  | R6-Gal4;UAS-AGES>UAS-*htt*<sup>RNAi</sup> | + | 1.23 | 0.96 | a |
| Bout Duration Nighttime<br>(mins) | *w<sup>1118</sup>*>UAS-*htt*<sup>RNAi</sup> | + | 15.98 | 8.00 | a |
|  | R6-Gal4;UAS-AGES>*w<sup>1118</sup>* | + | 13.56 | 3.78 | b |
|  | R6-Gal4;UAS-AGES>UAS-*htt*<sup>RNAi</sup> | - | 13.65 | 4.64 | b |
|  | R6-Gal4;UAS-AGES>UAS-*htt*<sup>RNAi</sup> | + | **12.43** | **3.36** | **c** |
| Bout Number Nighttime | *w<sup>1118</sup>*>UAS-*htt*<sup>RNAi</sup> | + | 29.81 | 8.25 | a |
|  | R6-Gal4;UAS-AGES>*w<sup>1118</sup>* | + | 30.50 | 6.83 | a |
|  | R6-Gal4;UAS-AGES>UAS-*htt*<sup>RNAi</sup> | - | 23.05 | 7.38 | c |
|  | R6-Gal4;UAS-AGES>UAS-*htt*<sup>RNAi</sup> | + | 26.89 | 8.11 | b |
| Latency lights off<br>(mins) | *w<sup>1118</sup>*>UAS-*htt*<sup>RNAi</sup> | + | 1.02 | 1.07 | a |
|  | R6-Gal4;UAS-AGES>*w<sup>1118</sup>* | + | 0.39 | 0.39 | b |
|  | R6-Gal4;UAS-AGES>UAS-*htt*<sup>RNAi</sup> | - | 0.75 | 0.76 | a |
|  | R6-Gal4;UAS-AGES>UAS-*htt*<sup>RNAi</sup> | + | 0.64 | 0.57 | a |
| Max Velocity | *w<sup>1118</sup>*>UAS-*htt*<sup>RNAi</sup> | + | 10.77 | 3.08 | a |
|  | R6-Gal4;UAS-AGES>*w<sup>1118</sup>* | + | 10.82 | 3.72 | a |
|  | R6-Gal4;UAS-AGES>UAS-*htt*<sup>RNAi</sup> | - | 7.35 | 2.55 | b |
|  | R6-Gal4;UAS-AGES>UAS-*htt*<sup>RNAi</sup> | + | 7.78 | 2.89 | b |
| Max Velocity Daytime | *w<sup>1118</sup>*>UAS-*htt*<sup>RNAi</sup> | + | 11.02 | 3.22 | a |
|  | R6-Gal4;UAS-AGES>*w<sup>1118</sup>* | + | 11.78 | 3.83 | a |
|  | R6-Gal4;UAS-AGES>UAS-*htt*<sup>RNAi</sup> | - | 7.90 | 3.05 | b |
|  | R6-Gal4;UAS-AGES>UAS-*htt*<sup>RNAi</sup> | + | 8.83 | 3.45 | b |
| Max Velocity Nighttime | *w<sup>1118</sup>*>UAS-*htt*<sup>RNAi</sup> | + | 10.66 | 3.49 | a |
|  | R6-Gal4;UAS-AGES>*w<sup>1118</sup>* | + | 9.45 | 4.04 | a |
|  | R6-Gal4;UAS-AGES>UAS-*htt*<sup>RNAi</sup> | - | 6.76 | 2.38 | b |
|  | R6-Gal4;UAS-AGES>UAS-*htt*<sup>RNAi</sup> | + | 6.72 | 2.85 | b |

**Supplementary Table 11.**
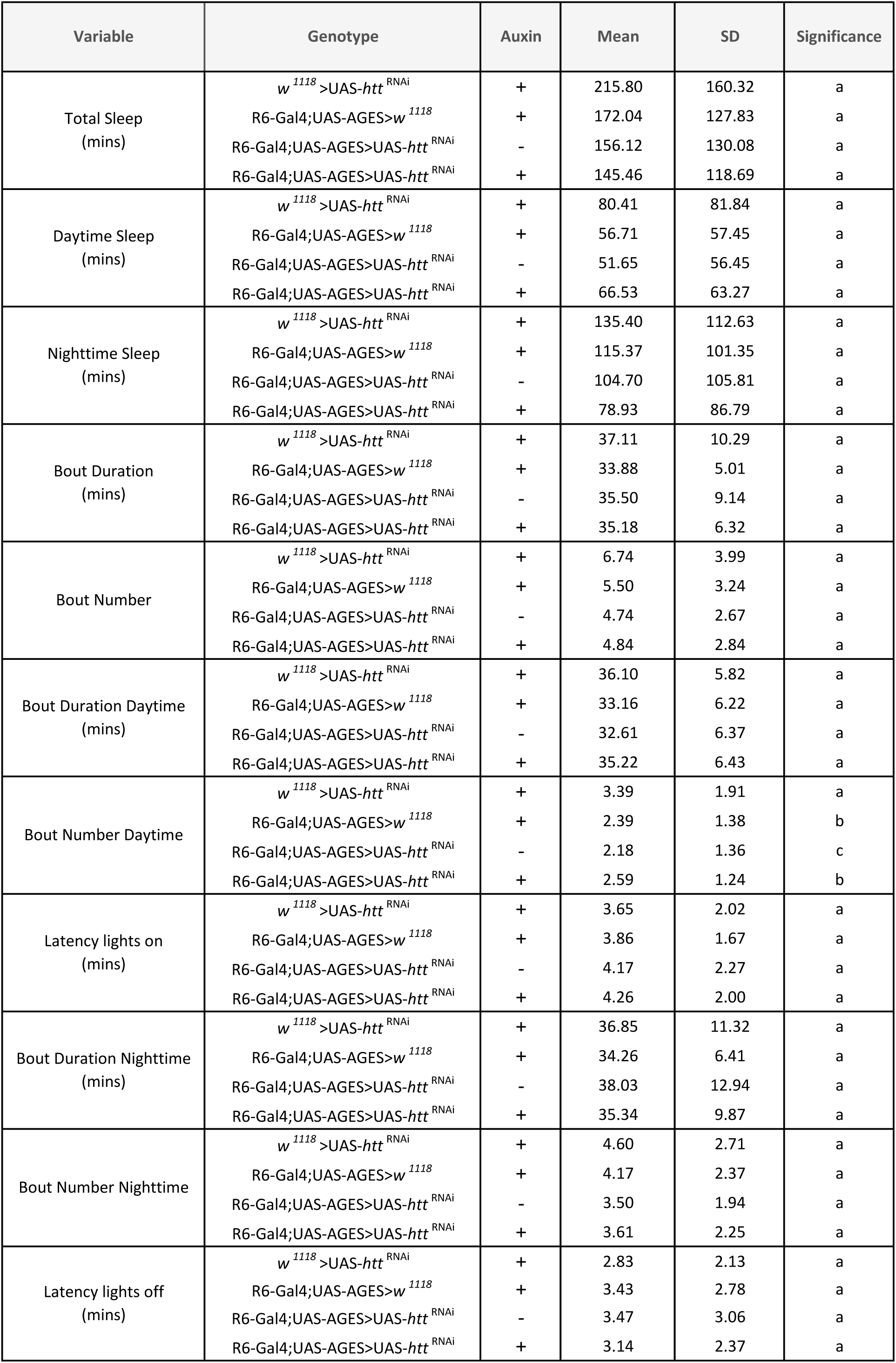
Sleep parameters in male flies with adult-specific *htt* downregulation in sLNvs (Ethoscope system, 25-min rule). Related to Supplementary Figure 4G. Values represent mean ± standard deviation (SD) for each variable. The presence (+) or absence (-) of Auxin to induce adult-specific transgene expression via the AGES system is indicated. Statistical analysis was performed using non-parametric methods; different letters in the Significance column denote statistically significant differences (*p<0.05*) among genotypes within each variable, determined by Kruskal-Wallis test followed by Dunn’s post-hoc test for multiple comparisons. Sample sizes: n=82-89 flies per genotype, N=3 independent experiments.

**Supplementary Table 12.** Sleep parameters in female flies with constitutive *htt* downregulation in sLNvs (DAM system, 25-min rule). Related to Supplementary Figure 3. Values represent Estimated Marginal Means (EMM) and 95% Confidence Intervals (95% CI) derived from Generalized Linear Mixed Models (GLMMs). The underlying error distribution specified for each behavioural variable is indicated in the Distribution column. Different letters in the Significance column denote statistically significant differences (*p<0.05*) among genotypes within each variable, determined by post-hoc pairwise comparisons of EMMs with Sidak’s p-value adjustment. Sample sizes: n=113-125 flies per genotype, N=5 independent experiments.

| Variable | Genotype | EMM | 95% CI |  | Significance | Distribution |
| --- | --- | --- | --- | --- | --- | --- |
|  |  |  | Lower | Upper |  |  |
| Total Sleep<br>(mins) | *w<sup>1118</sup>*>UAS-*htt*<sup>RNAi</sup> | 490.70 | 397.05 | 593.86 | b | beta |
|  | R6-Gal4>*w<sup>1118</sup>* | 411.35 | 324.60 | 510.67 | b |  |
|  | R6-Gal4>UAS-*htt*<sup>RNAi</sup> | **821.46** | **710.79** | **927.44** | **a** |  |
| Daytime Sleep<br>(mins) | *w<sup>1118</sup>*>UAS-*htt*<sup>RNAi</sup> | 179.67 | 130.40 | 239.99 | b | beta |
|  | R6-Gal4>*w<sup>1118</sup>* | 149.74 | 105.95 | 205.57 | b |  |
|  | R6-Gal4>UAS-*htt*<sup>RNAi</sup> | **337.43** | **267.72** | **408.89** | **a** |  |
| Nighttime Sleep<br>(mins) | *w<sup>1118</sup>*>UAS-*htt*<sup>RNAi</sup> | 333.37 | 286.03 | 381.66 | b | beta |
|  | R6-Gal4>*w<sup>1118</sup>* | 309.57 | 259.98 | 361.19 | b |  |
|  | R6-Gal4>UAS-*htt*<sup>RNAi</sup> | **503.84** | **456.04** | **546.28** | **a** |  |
| Bout Duration<br>(mins) | *w<sup>1118</sup>*>UAS-*htt*<sup>RNAi</sup> | 54.83 | 50.27 | 59.80 | b | lognormal |
|  | R6-Gal4>*w<sup>1118</sup>* | 50.88 | 46.40 | 55.79 | b |  |
|  | R6-Gal4>UAS-*htt*<sup>RNAi</sup> | **74.67** | **68.00** | **81.99** | **a** |  |
| Bout Number | *w<sup>1118</sup>*>UAS-*htt*<sup>RNAi</sup> | 9.55 | 8.22 | 11.08 | b | nbinom2 |
|  | R6-Gal4>*w<sup>1118</sup>* | 8.41 | 7.15 | 9.88 | b |  |
|  | R6-Gal4>UAS-*htt*<sup>RNAi</sup> | **12.04** | **10.57** | **13.71** | **a** |  |
| Bout Duration Daytime<br>(mins) | *w<sup>1118</sup>*>UAS-*htt*<sup>RNAi</sup> | 50.55 | 44.82 | 57.02 | b | lognormal |
|  | R6-Gal4>*w<sup>1118</sup>* | 44.89 | 39.61 | 50.88 | b |  |
|  | R6-Gal4>UAS-*htt*<sup>RNAi</sup> | **61.68** | **54.95** | **69.22** | **a** |  |
| Bout Number Daytime | *w<sup>1118</sup>*>UAS-*htt*<sup>RNAi</sup> | 4.10 | 3.39 | 4.97 | b | nbinom2 |
|  | R6-Gal4>*w<sup>1118</sup>* | 3.76 | 3.04 | 4.66 | b |  |
|  | R6-Gal4>UAS-*htt*<sup>RNAi</sup> | **6.58** | **5.52** | **7.84** | **a** |  |
| Latency lights on<br>(mins) | *w<sup>1118</sup>*>UAS-*htt*<sup>RNAi</sup> | 190.39 | 151.63 | 239.06 | a | lognormal |
|  | R6-Gal4>*w<sup>1118</sup>* | 190.87 | 148.42 | 245.47 | a |  |
|  | R6-Gal4>UAS-*htt*<sup>RNAi</sup> | **92.21** | **58.27** | **145.91** | **b** |  |
| Bout Duration Nighttime<br>(mins) | *w<sup>1118</sup>*>UAS-*htt*<sup>RNAi</sup> | 59.17 | 52.98 | 66.08 | b | lognormal |
|  | R6-Gal4>*w<sup>1118</sup>* | 53.46 | 48.13 | 59.40 | b |  |
|  | R6-Gal4>UAS-*htt*<sup>RNAi</sup> | **98.79** | **85.01** | **114.81** | **a** |  |
| Bout Number Nighttime | *w<sup>1118</sup>*>UAS-*htt*<sup>RNAi</sup> | 5.88 | 5.13 | 6.74 | a | nbinom2 |
|  | R6-Gal4>*w<sup>1118</sup>* | 5.56 | 4.83 | 6.40 | a |  |
|  | R6-Gal4>UAS-*htt*<sup>RNAi</sup> | 5.83 | 5.12 | 6.63 | a |  |
| Latency lights off<br>(mins) | *w<sup>1118</sup>*>UAS-*htt*<sup>RNAi</sup> | 38.52 | 24.73 | 60.01 | ab | lognormal |
|  | R6-Gal4>*w<sup>1118</sup>* | 54.12 | 34.43 | 85.08 | a |  |
|  | R6-Gal4>UAS-*htt*<sup>RNAi</sup> | 29.64 | 19.12 | 45.95 | b |  |

**Supplementary Table 13.**
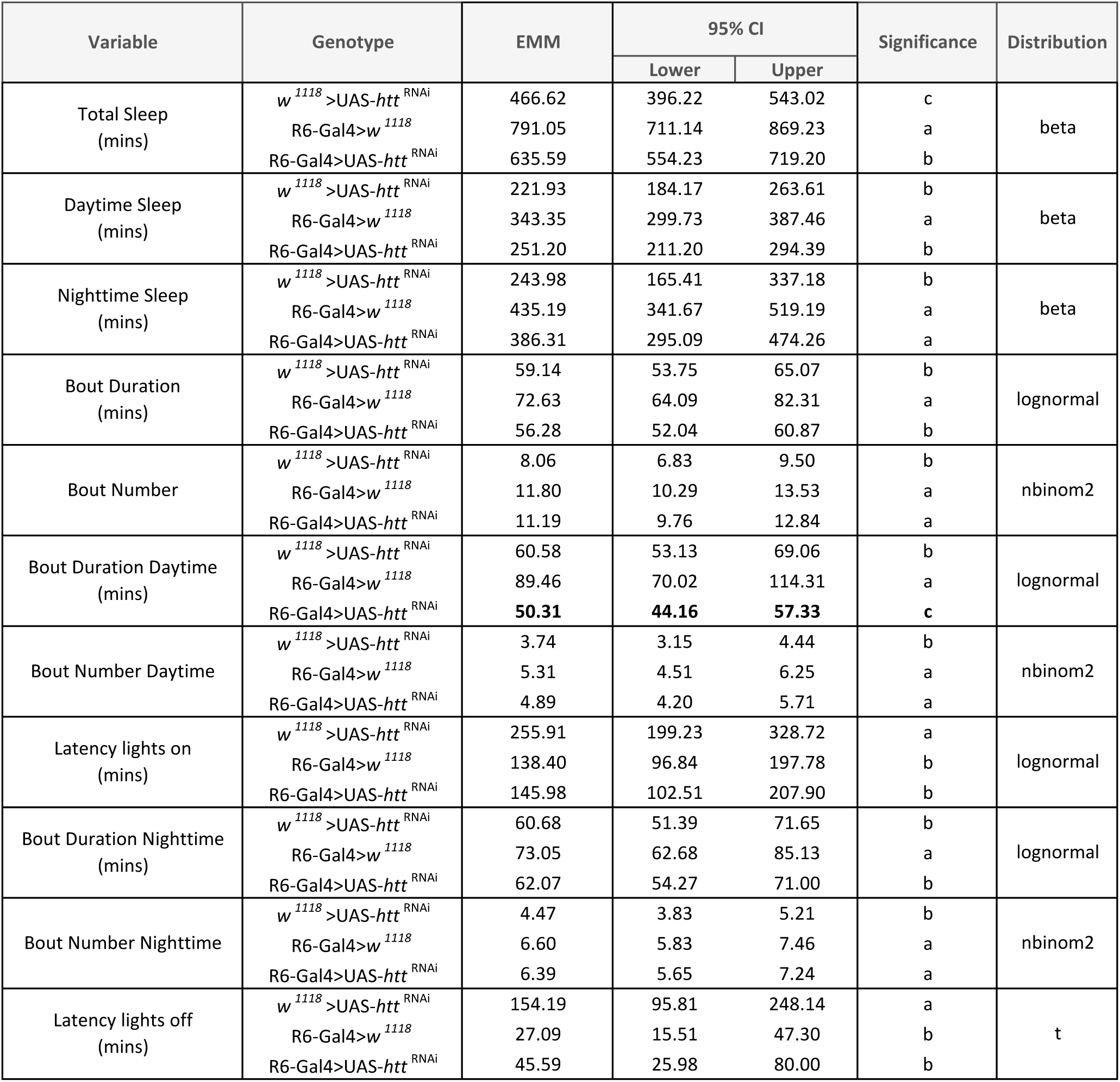
Sleep parameters in male flies with constitutive *htt* downregulation in sLNvs (DAM system, 25-min rule). Related to Supplementary Figure 2D. Values represent Estimated Marginal Means (EMM) and 95% Confidence Intervals (95% CI) derived from Generalized Linear Mixed Models (GLMMs). The underlying error distribution specified for each behavioural variable is indicated in the Distribution column. Different letters in the Significance column denote statistically significant differences (*p<0.05*) among genotypes within each variable, determined by post-hoc pairwise comparisons of EMMs with Sidak’s p-value adjustment. Sample sizes: n=63-64 flies per genotype, N=2 independent experiments.

**Supplementary Table 14.** Sleep parameters in female flies with constitutive *htt* downregulation and ANF-Emerald expression in sLNvs (DAM system, 5-min rule). Related to Supplementary Figure 5. Values represent Estimated Marginal Means (EMM) and 95% Confidence Intervals (95% CI) derived from Generalized Linear Mixed Models (GLMMs). The underlying error distribution specified for each behavioural variable is indicated in the Distribution column. Different letters in the Significance column denote statistically significant differences (*p<0.05*) among genotypes within each variable, determined by post-hoc pairwise comparisons of EMMs with Sidak’s p-value adjustment. Sample sizes: n=31-64 flies per genotype, N=2 independent experiments.

| Variable | Genotype | EMM | 95% CI |  | Significance | Distribution |
| --- | --- | --- | --- | --- | --- | --- |
|  |  |  | Lower | Upper |  |  |
| Total Sleep (mins) | *w<sup>1118</sup>*>UAS-preproANF-Emerald;UAS-*htt<sup>RNAi</sup>* | 759.11 | 675.18 | 842.00 | b | beta |
|  | R6-Gal4>*w<sup>1118</sup>* | 844.82 | 766.05 | 920.62 | b |  |
|  | R6-Gal4>UAS-preproANF-Emerald | 799.88 | 717.98 | 879.74 | b |  |
|  | R6-Gal4>UAS-preproANF-Emerald;UAS-*htt<sup>RNAi</sup>* | **1088.05** | **1027.11** | **1142.60** | **a** |  |
|  | R6-Gal4>UAS-*htt<sup>RNAi</sup>* | **1045.70** | **931.47** | **1142.47** | **a** |  |
| Daytime Sleep (mins) | *w<sup>1118</sup>*>UAS-preproANF-Emerald;UAS-*htt<sup>RNAi</sup>* | 269.04 | 228.08 | 312.68 | b | beta |
|  | R6-Gal4>*w<sup>1118</sup>* | 326.70 | 283.32 | 371.07 | b |  |
|  | R6-Gal4>UAS-preproANF-Emerald | 275.25 | 233.96 | 319.05 | b |  |
|  | R6-Gal4>UAS-preproANF-Emerald;UAS-*htt<sup>RNAi</sup>* | **448.00** | **404.28** | **489.12** | **a** |  |
|  | R6-Gal4>UAS-*htt<sup>RNAi</sup>* | **455.94** | **392.96** | **513.18** | **a** |  |
| Nighttime Sleep (mins) | *w<sup>1118</sup>*>UAS-preproANF-Emerald;UAS-*htt<sup>RNAi</sup>* | 502.49 | 450.15 | 548.54 | c | beta |
|  | R6-Gal4>*w<sup>1118</sup>* | 541.83 | 494.38 | 582.08 | bc |  |
|  | R6-Gal4>UAS-preproANF-Emerald | 543.68 | 495.57 | 584.29 | bc |  |
|  | R6-Gal4>UAS-preproANF-Emerald;UAS-*htt<sup>RNAi</sup>* | 649.39 | 624.06 | 668.58 | a |  |
|  | R6-Gal4>UAS-*htt<sup>RNAi</sup>* | 601.11 | 543.15 | 642.78 | ab |  |
| Bout Duration (mins) | *w<sup>1118</sup>*>UAS-preproANF-Emerald;UAS-*htt<sup>RNAi</sup>* | 21.25 | 17.53 | 25.76 | ab | lognormal |
|  | R6-Gal4>*w<sup>1118</sup>* | 19.71 | 16.61 | 23.39 | b |  |
|  | R6-Gal4>UAS-preproANF-Emerald | 17.89 | 15.28 | 20.96 | b |  |
|  | R6-Gal4>UAS-preproANF-Emerald;UAS-*htt<sup>RNAi</sup>* | 27.60 | 23.07 | 33.02 | a |  |
|  | R6-Gal4>UAS-*htt<sup>RNAi</sup>* | 24.74 | 17.75 | 34.47 | ab |  |
| Bout Number | *w<sup>1118</sup>*>UAS-preproANF-Emerald;UAS-*htt<sup>RNAi</sup>* | 43.55 | 37.86 | 50.10 | a | nbinom2 |
|  | R6-Gal4>*w<sup>1118</sup>* | 48.84 | 43.56 | 54.75 | a |  |
|  | R6-Gal4>UAS-preproANF-Emerald | 50.58 | 44.55 | 57.42 | a |  |
|  | R6-Gal4>UAS-preproANF-Emerald;UAS-*htt<sup>RNAi</sup>* | 47.50 | 41.39 | 54.50 | a |  |
|  | R6-Gal4>UAS-*htt<sup>RNAi</sup>* | 55.52 | 44.90 | 68.65 | a |  |
| Bout Duration Daytime (mins) | *w<sup>1118</sup>*>UAS-preproANF-Emerald;UAS-*htt<sup>RNAi</sup>* | 15.64 | 13.05 | 18.73 | a | lognormal |
|  | R6-Gal4>*w<sup>1118</sup>* | 15.06 | 11.93 | 19.02 | a |  |
|  | R6-Gal4>UAS-preproANF-Emerald | 10.82 | 9.45 | 12.39 | b |  |
|  | R6-Gal4>UAS-preproANF-Emerald;UAS-*htt<sup>RNAi</sup>* | 16.76 | 13.93 | 20.17 | a |  |
|  | R6-Gal4>UAS-*htt<sup>RNAi</sup>* | 17.86 | 12.71 | 25.09 | a |  |
| Bout Number Daytime | *w<sup>1118</sup>*>UAS-preproANF-Emerald;UAS-*htt<sup>RNAi</sup>* | 19.73 | 16.98 | 22.93 | c | nbinom2 |
|  | R6-Gal4>*w<sup>1118</sup>* | 25.43 | 21.97 | 29.43 | b |  |
|  | R6-Gal4>UAS-preproANF-Emerald | 26.49 | 22.90 | 30.65 | ab |  |
|  | R6-Gal4>UAS-preproANF-Emerald;UAS-*htt<sup>RNAi</sup>* | 31.16 | 26.99 | 35.98 | a |  |
|  | R6-Gal4>UAS-*htt<sup>RNAi</sup>* | 32.33 | 26.72 | 39.14 | a |  |
| Latency lights on (mins) | *w<sup>1118</sup>*>UAS-preproANF-Emerald;UAS-*htt<sup>RNAi</sup>* | 2.21 | 1.38 | 3.52 | a | lognormal |
|  | R6-Gal4>*w<sup>1118</sup>* | 1.36 | 0.88 | 2.10 | b |  |
|  | R6-Gal4>UAS-preproANF-Emerald | 1.65 | 0.99 | 2.75 | ab |  |
|  | R6-Gal4>UAS-preproANF-Emerald;UAS-*htt<sup>RNAi</sup>* | 0.68 | 0.38 | 1.21 | c |  |
|  | R6-Gal4>UAS-*htt<sup>RNAi</sup>* | 0.56 | 0.20 | 1.55 | bc |  |
| Bout Duration Nighttime (mins) | *w<sup>1118</sup>*>UAS-preproANF-Emerald;UAS-*htt<sup>RNAi</sup>* | 37.46 | 25.95 | 54.08 | b | lognormal |
|  | R6-Gal4>*w<sup>1118</sup>* | 35.52 | 26.07 | 48.40 | b |  |
|  | R6-Gal4>UAS-preproANF-Emerald | 32.25 | 24.71 | 42.10 | b |  |
|  | R6-Gal4>UAS-preproANF-Emerald;UAS-*htt<sup>RNAi</sup>* | 84.28 | 58.25 | 121.92 | a |  |
|  | R6-Gal4>UAS-*htt<sup>RNAi</sup>* | 42.39 | 27.02 | 66.49 | b |  |
| Bout Number Nighttime | *w<sup>1118</sup>*>UAS-preproANF-Emerald;UAS-*htt<sup>RNAi</sup>* | 23.87 | 19.92 | 28.62 | a | nbinom2 |
|  | R6-Gal4>*w<sup>1118</sup>* | 23.47 | 20.37 | 27.04 | a |  |
|  | R6-Gal4>UAS-preproANF-Emerald | 24.05 | 20.64 | 28.02 | a |  |
|  | R6-Gal4>UAS-preproANF-Emerald;UAS-*htt<sup>RNAi</sup>* | 16.25 | 12.76 | 20.70 | b |  |
|  | R6-Gal4>UAS-*htt<sup>RNAi</sup>* | 23.37 | 17.77 | 30.72 | ab |  |
| Latency lights off (mins) | *w<sup>1118</sup>*>UAS-preproANF-Emerald;UAS-*htt<sup>RNAi</sup>* | 0.57 | 0.42 | 0.77 | a | lognormal |
|  | R6-Gal4>*w<sup>1118</sup>* | 0.49 | 0.31 | 0.79 | a |  |
|  | R6-Gal4>UAS-preproANF-Emerald | 0.35 | 0.24 | 0.50 | a |  |
|  | R6-Gal4>UAS-preproANF-Emerald;UAS-*htt<sup>RNAi</sup>* | 0.38 | 0.23 | 0.61 | a |  |
|  | R6-Gal4>UAS-*htt<sup>RNAi</sup>* | 0.28 | 0.15 | 0.53 | a |  |
| Activity Index | *w<sup>1118</sup>*>UAS-preproANF-Emerald;UAS-*htt<sup>RNAi</sup>* | 2.04 | 1.95 | 2.14 | c | lognormal |
|  | R6-Gal4>*w<sup>1118</sup>* | 2.04 | 1.94 | 2.14 | c |  |
|  | R6-Gal4>UAS-preproANF-Emerald | 1.75 | 1.68 | 1.82 | d |  |
|  | R6-Gal4>UAS-preproANF-Emerald;UAS-*htt<sup>RNAi</sup>* | **2.34** | **2.16** | **2.54** | **b** |  |
|  | R6-Gal4>UAS-*htt<sup>RNAi</sup>* | **2.75** | **2.46** | **3.07** | **a** |  |
| Activity Index Daytime | *w<sup>1118</sup>*>UAS-preproANF-Emerald;UAS-*htt<sup>RNAi</sup>* | 1.94 | 1.85 | 2.05 | c | lognormal |
|  | R6-Gal4>*w<sup>1118</sup>* | 1.93 | 1.83 | 2.03 | c |  |
|  | R6-Gal4>UAS-preproANF-Emerald | 1.63 | 1.55 | 1.71 | d |  |
|  | R6-Gal4>UAS-preproANF-Emerald;UAS-*htt<sup>RNAi</sup>* | **2.29** | **2.09** | **2.52** | **b** |  |
|  | R6-Gal4>UAS-*htt<sup>RNAi</sup>* | **2.78** | **2.43** | **3.17** | **a** |  |
| Activity Index Nighttime | *w<sup>1118</sup>*>UAS-preproANF-Emerald;UAS-*htt<sup>RNAi</sup>* | 2.26 | 2.08 | 2.46 | bc | lognormal |
|  | R6-Gal4>*w<sup>1118</sup>* | 2.36 | 2.15 | 2.60 | b |  |
|  | R6-Gal4>UAS-preproANF-Emerald | 2.10 | 1.94 | 2.27 | c |  |
|  | R6-Gal4>UAS-preproANF-Emerald;UAS-*htt<sup>RNAi</sup>* | 2.41 | 2.19 | 2.65 | ab |  |
|  | R6-Gal4>UAS-*htt<sup>RNAi</sup>* | 2.75 | 2.41 | 3.14 | a |  |

**Supplementary Table 15.**
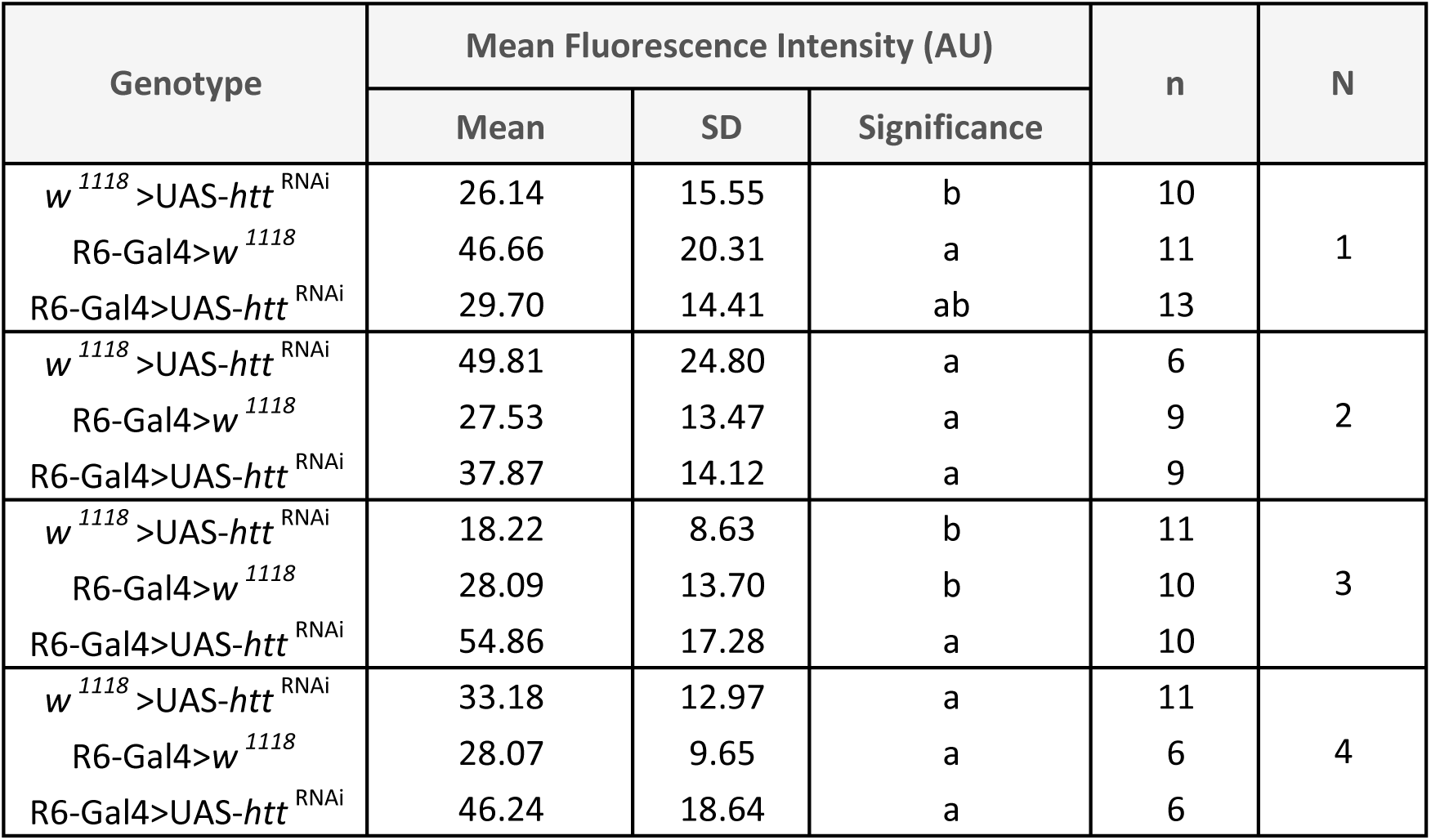
Quantification of total PDF immunoreactivity (mean fluorescence intensity) in the sLNvs terminals at ZT2. Related to Figure 4A. Different letters denote statistically significant differences (p<0.05) within each experimental replicate (N), determined by One-way ANOVA followed by Tukey’s post hoc test. SD: standard deviation; AU: arbitrary units; n: sample size.

| Genotype | Mean Fluorescence Intensity (AU) |  |  | n | N |
| --- | --- | --- | --- | --- | --- |
|  | Mean | SD | Significance |  |  |
| *w<sup>1118</sup>*>UAS-*htt*<sup>RNAi</sup> | 26.14 | 15.55 | b | 10 | 1 |
| R6-Gal4>*w<sup>1118</sup>* | 46.66 | 20.31 | a | 11 |  |
| R6-Gal4>UAS-*htt*<sup>RNAi</sup> | 29.70 | 14.41 | ab | 13 |  |
| *w<sup>1118</sup>*>UAS-*htt*<sup>RNAi</sup> | 49.81 | 24.80 | a | 6 | 2 |
| R6-Gal4>*w<sup>1118</sup>* | 27.53 | 13.47 | a | 9 |  |
| R6-Gal4>UAS-*htt*<sup>RNAi</sup> | 37.87 | 14.12 | a | 9 |  |
| *w<sup>1118</sup>*>UAS-*htt*<sup>RNAi</sup> | 18.22 | 8.63 | b | 11 | 3 |
| R6-Gal4>*w<sup>1118</sup>* | 28.09 | 13.70 | b | 10 |  |
| R6-Gal4>UAS-*htt*<sup>RNAi</sup> | 54.86 | 17.28 | a | 10 |  |
| *w<sup>1118</sup>*>UAS-*htt*<sup>RNAi</sup> | 33.18 | 12.97 | a | 11 | 4 |
| R6-Gal4>*w<sup>1118</sup>* | 28.07 | 9.65 | a | 6 |  |
| R6-Gal4>UAS-*htt*<sup>RNAi</sup> | 46.24 | 18.64 | a | 6 |  |

**Supplementary Table 16.**
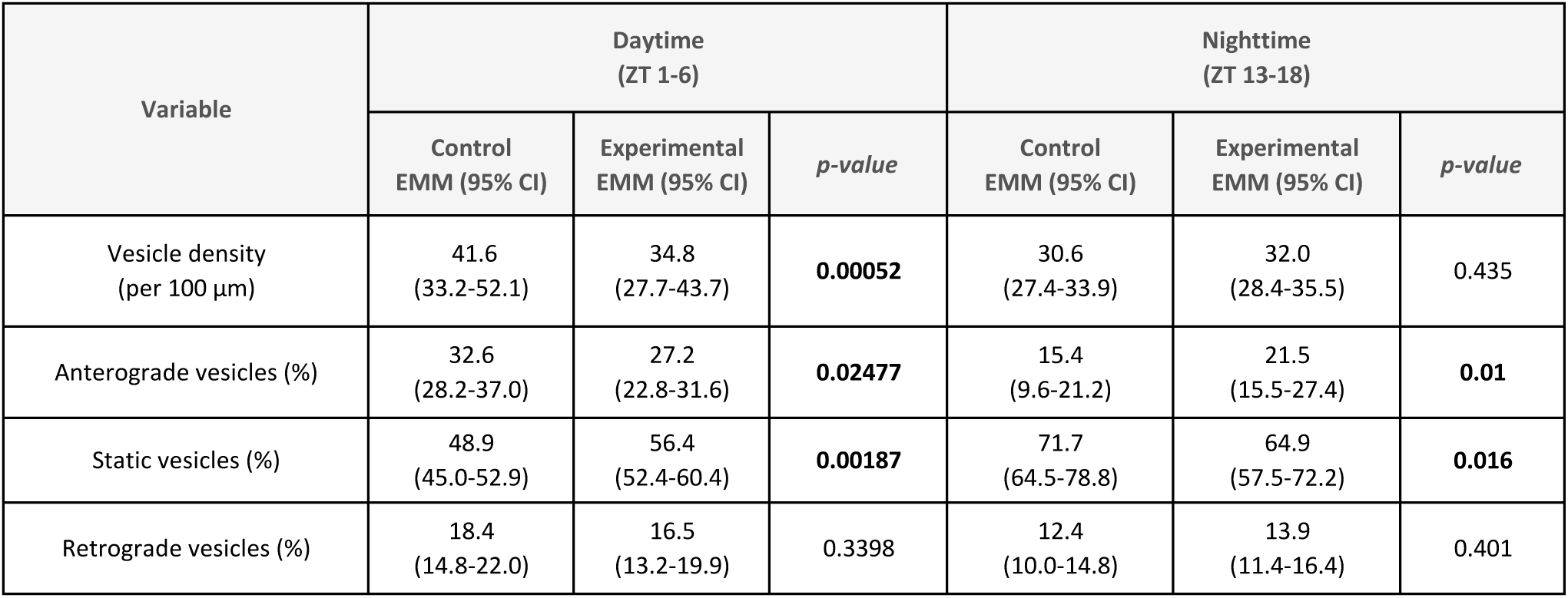
Vesicle transport parameters in sLNv axons during daytime (ZT 1–6, related to. Figure 4 C-E**) and nighttime (ZT 13–18, related to** Figure 4 H-J). Values represent Estimated Marginal Means (EMMs) with 95% Confidence Intervals (95% CI) in parentheses. Statistical significance (p-values) was derived from hierarchical Generalized Linear Mixed Models (GLMMs) with fly and hemisphere included as nested random effects. Statistically significant p-values (*p <0.05*) are highlighted in bold. Genotypes: control (*R6-Gal4>UAS-preproANF-Emerald*) and experimental (*R6-Gal4>UAS-preproANF-Emerald;;UAS-htt*^RNAi^).

| Variable | Daytime<br>(ZT 1-6) |  |  | Nighttime<br>(ZT 13-18) |  |  |
| --- | --- | --- | --- | --- | --- | --- |
|  | Control<br>EMM (95% CI) | Experimental<br>EMM (95% CI) | *p-value* | Control<br>EMM (95% CI) | Experimental<br>EMM (95% CI) | *p-value* |
| Vesicle density<br>(per 100 $\mu\text{m}$) | 41.6<br>(33.2-52.1) | 34.8<br>(27.7-43.7) | **0.00052** | 30.6<br>(27.4-33.9) | 32.0<br>(28.4-35.5) | 0.435 |
| Anterograde vesicles (%) | 32.6<br>(28.2-37.0) | 27.2<br>(22.8-31.6) | **0.02477** | 15.4<br>(9.6-21.2) | 21.5<br>(15.5-27.4) | **0.01** |
| Static vesicles (%) | 48.9<br>(45.0-52.9) | 56.4<br>(52.4-60.4) | **0.00187** | 71.7<br>(64.5-78.8) | 64.9<br>(57.5-72.2) | **0.016** |
| Retrograde vesicles (%) | 18.4<br>(14.8-22.0) | 16.5<br>(13.2-19.9) | 0.3398 | 12.4<br>(10.0-14.8) | 13.9<br>(11.4-16.4) | 0.401 |

**Supplementary Table 17.**
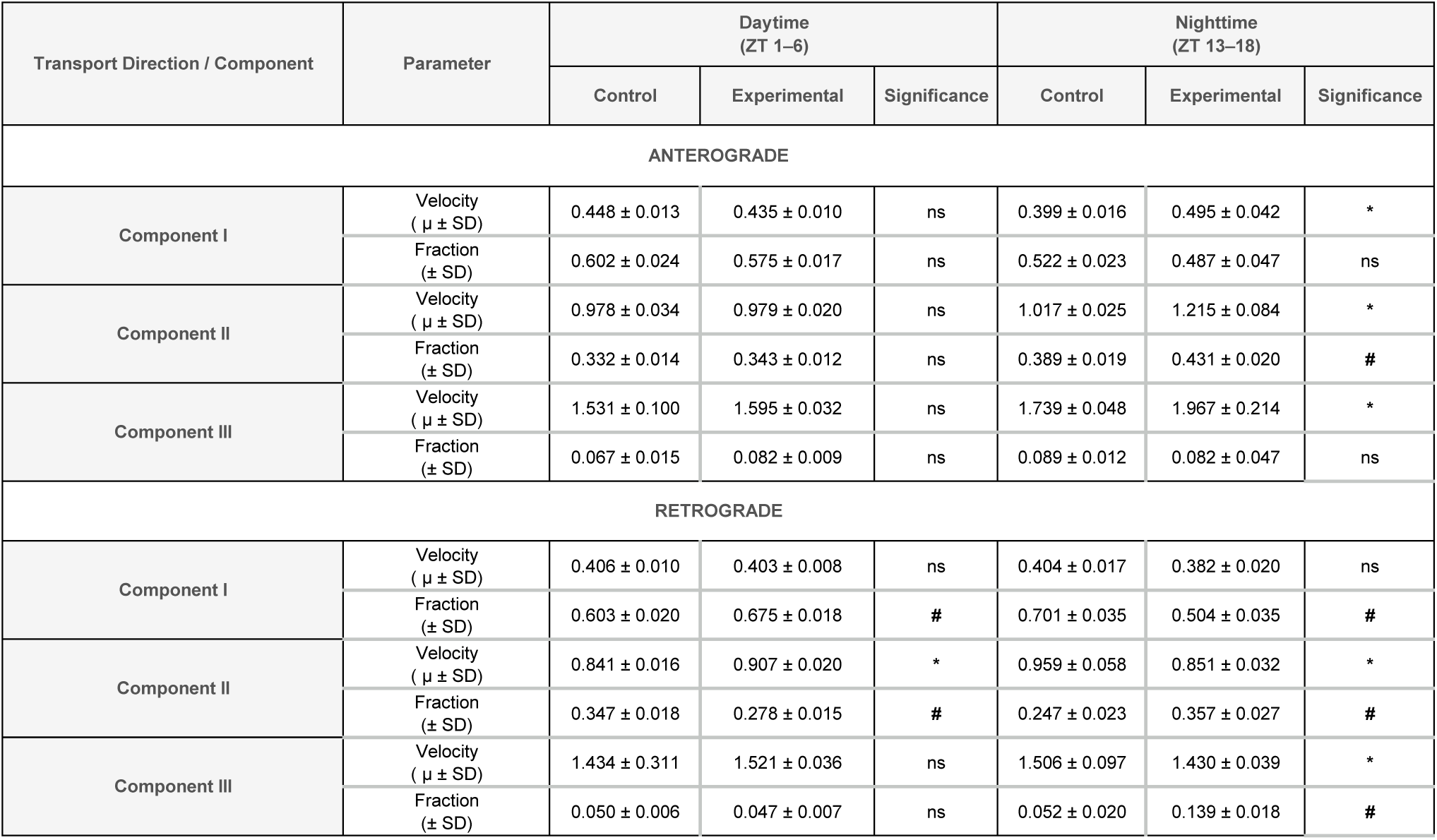
Segmental velocity components of moving vesicles in sLNv axons during daytime (ZT 1–6, related to Figure 4 F,G) and nighttime (ZT 13–18, related to Figure 4 K,L). Data represent the mean ± standard deviation (SD) derived from a triple-Gaussian Mixture Model (GMM) fitting, describing slow (Component I), middle (Component II), and fast (Component III) transport modes. Statistical significance was determined via two-tailed Z-tests comparing the parameters (component velocities μ and fractions θ) and their respective standard errors (SE) derived from the GMM uncertainty matrix between genotypes. In the significance columns, asterisks (*) indicate significant differences in velocity component (μ), while numeral symbols (#) indicate significant differences in fraction component (θ); ns: non-significant (*p>0.05*). Genotypes: control (*R6-Gal4>UAS-preproANF-Emerald*) and experimental (*R6-Gal4>UAS-preproANF-Emerald;;UAS-htt*^RNAi^).

## MOVIE LEGENDS

**Movie 1. Dense-core vesicle axonal transport in a control sLNv**. Representative movie showing the axonal transport of ANF::Emerald loaded vesicles in real time in the brain of a R6-Gal4>preproANF-Emerald fly during the light-phase (ZT4). Movie showing time in seconds (5 frames/s) oriented so that right– to left-moving vesicles correspond to anterograde transport and left to right to retrograde movement. Box shows the ascending portion of the axon, where transport was assessed. Scale bar, 10 µm. [View online: https://drive.google.com/file/d/13LbwcThcFHOstvGhvqL-HOagWsc8QJ8d/view?usp=sharing]

**Movie 2. Dense-core vesicle axonal transport in an *htt*^RNAi^ sLNv**. Representative movie showing the axonal transport of ANF::Emerald loaded vesicles in real time in the brain of a R6-Gal4>preproANF-Emerald;;UAS-*htt*^RNAi^ fly during the light-phase (ZT4). Movie showing time in seconds (5 frames/s) oriented so that right– to left-moving vesicles correspond to anterograde transport and left to right to retrograde movement. Box shows the ascending portion of the axon, where transport was assessed. Scale bar, 10 µm. [View online: https://drive.google.com/file/d/13gj67bfm9YYN4OwaWV7X1212xnEkVV27/view?usp=sharing]

## ACKNOWLEDGEMENTS

We thank Dr. Ignacio Spiousas for their assistance with the statistical analysis, Dr. Shermali Gunawardena for sharing reagents and Biochem. Alejandra Attorresi for microscopy support. This work was funded by CONICET doctoral scholarships (ABV, CAY, LAD, FFC, IML, ID, PLB), AGENCIA I+D+i-FONCYT grant PICT 2018-2030 (NIM), AGENCIA I+D+i-FONCYT grant PICT 2020-1645 (NIM), CONICET grant PIP 2022-2024 GI 11220210100779 CO (NIM), Chan Zuckerberg Initiative grant CP2-1-0000000310 (NIM), Pew Innovation Fund GR127197 (TLF), UBACyT 20020220200095BA (TLF), CONICET grant PIP 2021-2023 11220200101831 CO (EJB), PIBAA 2022-2023 28720210100905 CO (EJB), AGENCIA I+D+i-FONCYT grants PICT-2020-SERIE A-01240 and PICT-PRH-2021-00009 (EJB), and FOCEM-Mercosur COF 03/11 (IBioBA).

## AUTHOR CONTRIBUTIONS

N.I.M., A.B-V., A.R., T.L.F. and E.J.B. conceived and designed the experiments (Conceptualization). A.B-V., C.A., F.F-C., A.R., L.A.D., I.D., I.M.L., P.L.B., M.P-L., C.S.F. and E.J.B. performed the experiments and data collection (Investigation) with the aid of C.N.P. for microscopy support. A.B-V adapted the custom analysis code (Software) and performed the statistical analysis (Formal Analysis). A.B-V., C.A. and C.S.F curated the raw imaging data (Data Curation). A.B-V., C.A., T.L.F. and N.I.M. developed the DCV dynamics visualization in the axons of adult *Drosophila* sLNvs techinique (Methodology). N.I.M., T.L.F. and E.J.B. supervised the project and acquired the funding (Supervision; Funding Acquisition). N.I.M. wrote most of the original draft of the manuscript (Writing – Original Draft). All authors reviewed, edited, and approved the final version (Writing – Review & Editing).

